# Calibrated rare variant genetic risk scores for complex disease prediction using large exome sequence repositories

**DOI:** 10.1101/2020.02.03.931519

**Authors:** Ricky Lali, Michael Chong, Arghavan Omidi, Pedrum Mohammadi-Shemirani, Ann Le, Guillaume Paré

## Abstract

Rare variants are collectively numerous and may underlie a considerable proportion of complex disease risk. However, identifying genuine rare variant associations is challenging due to small effect sizes, presence of technical artefacts, and heterogeneity in population structure. We hypothesized that rare variant burden over a large number of genes can be combined into predictive rare variant genetic risk score (RVGRS). We propose a novel method (RV-EXCALIBER) that leverages summary-level data from a large public exome sequencing database (gnomAD) as controls and robustly calibrates rare variant burden to account for the aforementioned biases. A RVGRS was found to strongly associate with coronary artery disease (CAD) in European and South Asian populations. Calibrated RVGRS capture the aggregate effect of rare variants through a polygenic model of inheritance, identifies 1.5% of the population with substantial risk of early CAD, and confers risk even when adjusting for known Mendelian CAD genes, clinical risk factors, and common variant gene scores.

## INTRODUCTION

Rare variants have been hypothesized to contribute significantly to the missing heritability of complex diseases^1,2^, raising the possibility of complementing common variants polygenic scores with rare variant scores. However, the ability to identify robust associations between rare variants and complex diseases is inherently limited by sample size. Though novel statistical techniques have been developed to conduct rare variant association, which involve collapsing all rare variants within a given genetic unit (i.e. a gene or gene-set) into a single count^3–7^, the power to detect rare variant associations of modest effect remains limited. Large publicly available sequencing consortia can be employed as control databases in order to markedly increase power. However, there is currently no method to calibrate rare variant burden between test samples and public databases despite both population and sequencing differences, thus limiting the ability to construct well-powered rare variant genetic risk score.

The recent public releases of the Exome Aggregation Consortium (ExAC)^8^ and genome Aggregation Database (gnomAD)^9^ have allowed users to obtain precise allele frequency estimates in many different ethnic populations. In fact, ExAC and gnomAD are often used to distinguish between benign and pathogenic variants present in single cases or families based on rarity^10^. The summary-level data contained within these public sequencing databases can also be leveraged beyond single case and pedigree analysis. Specifically, the allele frequency information in these databases is representative of the expected allele frequencies in the general population, which can be utilized as a control distribution for rare variant association studies. However, incorporating external databases into rare variant association testing presents many population-specific and technical challenges. Specifically, populations in both ExAC and gnomAD are geographically diverse (even within single continents) and consist of sample pools that utilize different exome capture chemistries and variant calling algorithms which may be distinct from those used in a local case population. Consequently, adjustments to gene-based expected frequencies are required in order to mitigate artefactual association signals. These adjustments are especially vital for rare variants as they have shown to exhibit a higher degree of geographical specificity compared to common variants ^11–13^, and accuracy of calling rarer variants largely depends upon sample size for joint variant calling algorithms^14^. As such, sources of heterogeneity in both population substructure and sequencing technology must be addressed to leverage the full capabilities of publicly available sequencing datasets as external control datasets.

Here, we propose a novel framework, Rare Variant Exome CALIBration using External Repositories (RV-EXCALIBER), which leverages the large sample size of gnomAD as a control dataset to conduct rare variant association testing. In brief, RV-EXCALIBER works by instituting 1) a correction that accounts for the increased variance due the presence of rare variants in LD, 2) an individual-level correction factor (iCF) that accounts for global variations in population substructure and sequencing technology and 3) a gene-level correction factor (gCF) that accounts for granular deviations in mutation burden bias amongst gene sets. As such, the iCF and gCF act by calibrating the mutation loads in gnomAD (control) to an input case dataset on a sample-by-sample basis on the whole-exome level and gene-set level, respectively. Through simulations, we demonstrate that RV-EXCALIBER adequately adjusts for sequencing false negatives and population-specific effects while also partially mitigating power loss induced by sequencing false-positives. We also demonstrate the necessity for rare variant calibration by comparing deviations in mutation load between Genome In A Bottle (GIAB) consortium reference samples and gnomAD. Lastly, we demonstrate a novel application for our framework through constructing individually calibrated rare variant gene risk scores (RVGRS) for a prevalent complex disorder, coronary artery disease (CAD).

## RESULTS

### Single gene simulations using the correction factors (CF)

In order to evaluate the effect of incorporating the CFs on rare variant gene-based association, we conducted a series of simulations and determined the CF’s effect on **1)** estimated odds ratio (OR) under strong confounding conditions of sequencing false negatives (SFN), sequencing false positives (SFP) and population-specific factors (PSF) (Figure 1 A-C) and **2)** power to detect association according to SFN (0 to 1), SFN (0 to 1), and PSF (0 to 2) under a model of association assuming a true OR of 1.3 (Figure 1 D-F). To accomplish this, we performed 100,000 simulations of aggregate rare allele count across 1,000 cases for a single hypothetical gene by varying the baseline aggregate probability of mutation (P) (i.e. 5%, which is a reasonable estimate given empirically-derived expected counts) based on true OR, SFN, SFP, and PSF. We found that incorporation of the CFs fully and significantly corrected the estimated OR to match the true OR when including either sequencing false negatives or population structure (Figure 1A-B). Conversely, we observed that incorporation of the CF over-corrects (i.e. estimated OR < true OR) when sequencing false positives are present (Figure 1C). We hold that the adjustment is still effective in the latter scenario given that CF-adjusted effect size of the gene-based association will be conservative.

**Figure 1:**
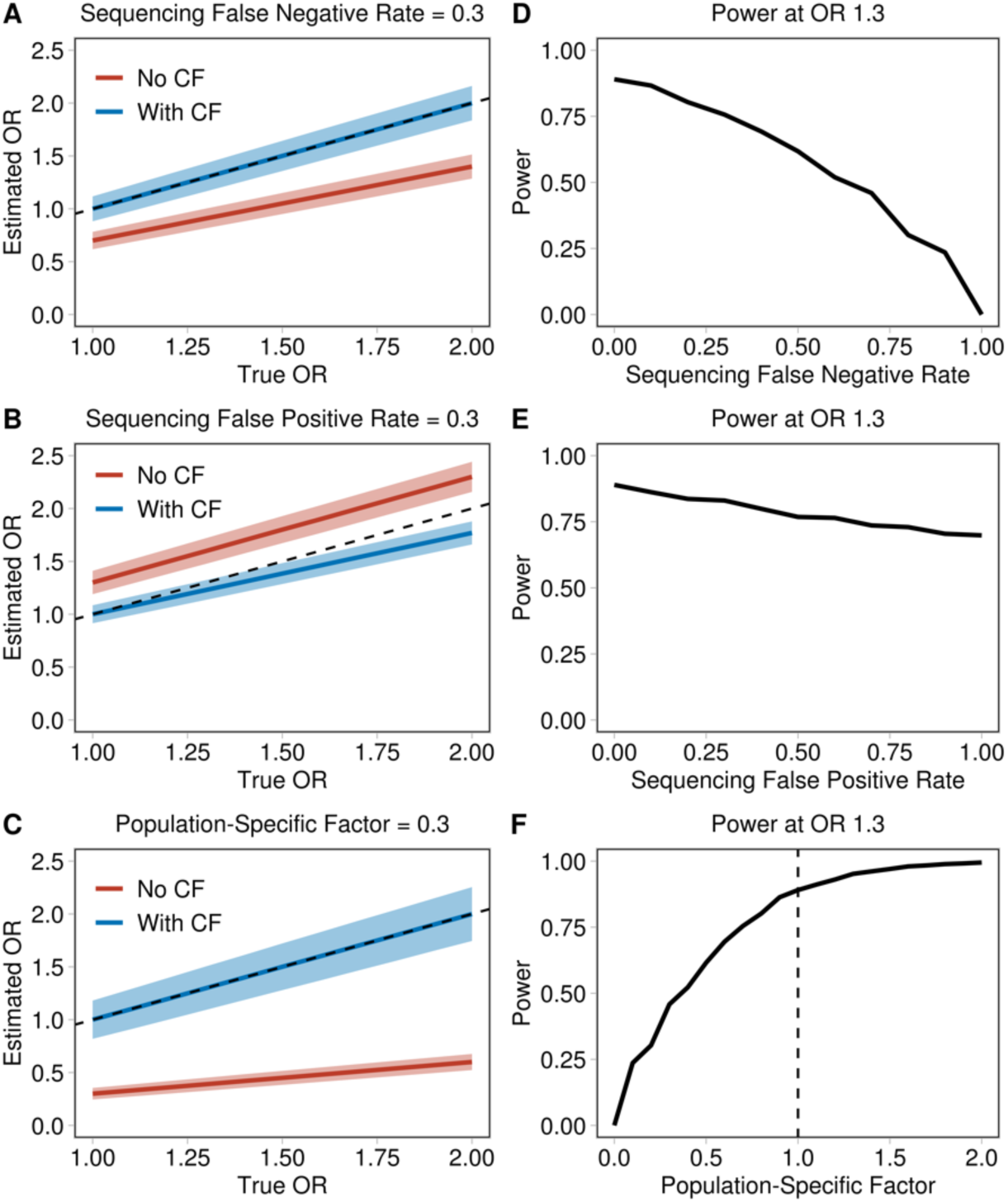
Simulated effects of SFN rate, SFP rate, and PSF on estimated OR and association power. Probability of mutation (P) within a gene was calculated using equation 2 by simulating true OR values from 1 to 2 in 0.1 stage intervals while keeping SFN, SFP and PSF fixed. Thereafter, an estimated OR was calculated for every unique P (**A-C; red lines**). CF values were calculated according to equation 3 and were used to determine an adjusted P (P*) (equation 4) while keeping SFN, SFP, and PSF fixed at the same values. Using P*, a CF-adjusted estimated OR was calculated (**A-C; blue lines**). Dashed lines in **A-C** indicate the line of expectation. CF-adjusted power curves were generated using different ranges of SFN (0 to 1), SFP (0 to 1), and PSF (0 to 2) at a fixed true OR value of 1.3 (**D-F**). Power for every value of SFN, SFP, and PSF was calculated as the proportion of all simulations with a p-value < 0.05. Dashed lines in **F** represents a null deviation in PSF. OR indicates odds ratio, SFN indicates sequencing false negatives, SFP indicates sequencing false positives, PSF indicates population-specific factor.

To estimate the power to detect a gene-based association, we similarly simulated 100,000 observed rare allele counts assuming a true OR of 1.3, which was the effect size necessary to achieve 80% power when no confounding factors exist (i.e. SFN & SFP rates of 0% and a PSF of 1) using our model (Figure 1D-F). Under the setting where sequencing false negatives are present, we expect a decrease in mutation rate and thus, power to detect association. Incorporation of the CF mitigates the reduction in power as we observe increased power to detect association across all SFN rates in the CF-adjusted compared to the unadjusted model. When only sequencing false positives are present, we show that incorporation of the CF ameliorates the spurious increase in power observed as mutation rate rises due to increasing rates of SFPs. It is important to note that the artificial increase in power observed in the unadjusted model is accredited solely to sequencing artefacts and not true variants, and thus would translate directly into increased type I error. Lastly, we observe an increase in power as the PSF becomes greater (i.e. the variant count in a test sample is greater than a reference sample due to reasons such as population substructure). In this setting, the power increase is expected as the mutation rate is increasing due to variants that are truly present because of population genetic factors, as opposed to variant artefacts observed in the un-adjusted model including SFP.

### GIAB consensus sequences highlights the role of population substructure in rare variant mutation load

As rare variants demonstrate a higher degree of geographical specificity and are under greater selective pressure than common variants, we sought to assess the impact of population substructure on rare variant burden through use of gold-standard consensus GIAB sequences. When comparing the Ashkenazi Jewish and Central European consensus sequences with the non-Finnish European (NFE) population in gnomAD, we observed significant deviations in iCF for rare variants (0 < MAF ≤ 0.01) (Ashkenazi: CF = 1.37; 95% CI, 1.22 to 1.55; p < 0.05 and Central European: CF = 0.60; 95% CI, 0.54 to 0.68; p < 0.05), where a CF=1 represents equal mutational loads. Deviations in variants burden were attenuated with higher MAF for the Ashkenazi Jewish consensus sequence while it remained significantly different across all MAF threshold bins for the Central European consensus sample, albeit less so at higher MAF (Figure 2 and Supplementary Table 4). To test whether CF deviations for rare variants would dissipate when calibrated against a more closely matched comparator population, we evaluated the CF between the Ashkenazi Jewish consensus sequence and the gnomAD ASJ population and the East Asian consensus sequence with the gnomAD EAS population. In these scenarios, the observed CFs closely approximated the expected CF across all MAF threshold bins, but rare variants still deviated most from the expected CF relative to the remaining MAF bins (Figure 2B, D and Supplementary Table 4).

**Figure 2:**
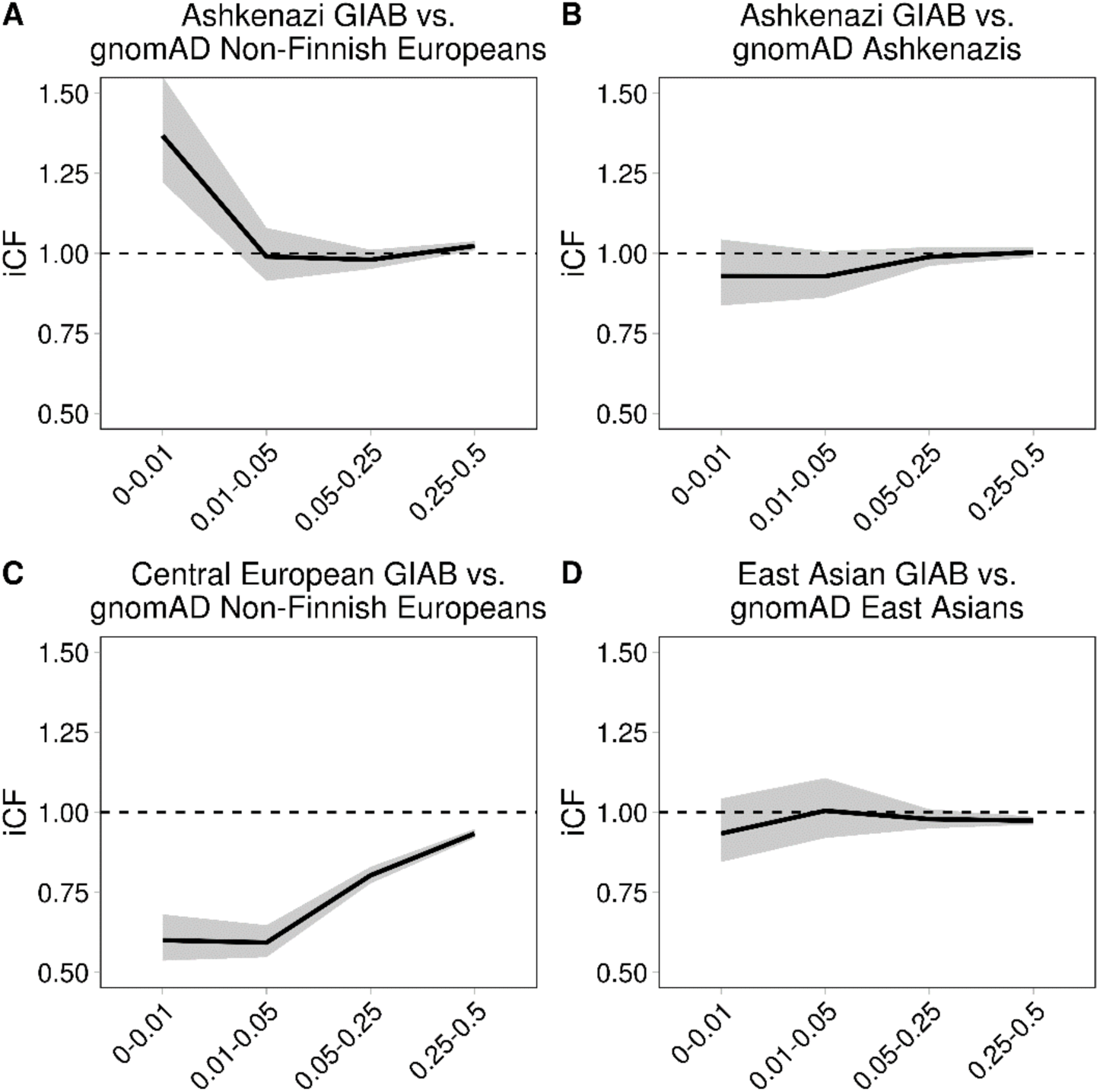
Comparison of iCF values for consensus GIAB samples stratified by allele frequency bin. Ethnic comparisons include the Ashkenazi GIAB sample (NA24385) with gnomAD Non-Finnish Europeans and gnomAD Ashkenazis (**A** and **B**), the Central European GIAB sample (NA12878) with gnomAD Non-Finnish Europeans (**C**), and the East Asian GIAB sample (NA24631) with gnomAD East Asians population (**D**). iCF values were calculated according to equation 6. AF bins were chosen to best stratify variants according to rare [0-0.01], low-frequency [0.01-0.05], common [0.5-0.25] and very common [0.25-0.5]. AFs were standardized to the minor allele. All shaded regions depict the 95% CI. Dashed lines indicate an iCF representing equal mutation loads between the GIAB sample and gnomAD (i.e. iCF = 1) across all protein-coding genes. GIAB indicates Genome In A Bottle.

Since the CFs deviated mostly from the expected values among rare variants for every population comparison, we wanted to rule out the possibility that the population effects driving these deviations were restricted to non-constrained genes as this would preclude use of a single CF to equilibrate mutation loads at the exome-wide level. To accomplish this, we allocated genes into deciles according to their missense constraint score^8^ and assessed whether there was significant heterogeneity among the CFs across all deciles. For each consensus sequence versus gnomAD comparison, we observed no significant heterogeneity after accounting for multiple hypotheses testing (pHet > 0.0125 (0.05/4) for all comparisons) (Supplementary Figure 2).

### iCF distribution among healthy controls participants from MIGen

A total of 8 WES cohorts were selected from the MIGen Exome Sequencing Study (Supplementary Table 1) as a discovery set for rare variant burden association (N=5,910 for cases and N=6,082 for controls). All analyses concerning the MIGen discovery cohorts were conducted using rare pathogenic alleles (i.e variants with a minor allele frequency <0.001 and nonsynonymous single nucleotide variants (SNV) with a Mendelian clinically applicable pathogenicity score >0.025 or loss-of-function variants; see Data Supplement). iCFs were calculated for healthy control participants from each cohort and their distributions were plotted to identify whether there were marked inter and intra-cohort differences in exome-wide rare variant mutation load. Across the 8 MIGen cohorts, the highest and lowest median iCF values were from the Italian Atherosclerosis Thrombosis and Vascular Biology (ATVB) (median=1.135; IQR, 0.909-1.363) and the Malmö Diet Cancer (MDC) studies (median=0.664; IQR, 0.487-0.841), respectively (Supplementary Table 5). All pairwise between cohorts comparison of iCF among control participants were found to be significantly different, even after adjusting for sex and the first 20 principle components, which were calculated based on common single nucleotide variants (Figure 3A). iCF distributions were also compared between cases and controls of individual MIGen cohorts, where significant differences were identified in 2 out of 8 studies. Specifically, iCF values were significantly lower among controls from the Registre Gironi del Cor (REGICOR) cohort (OR=0.33 per 1 unit change in iCF; 95%CI, 0.12-0.92; *P*=0.0352), but higher for MDC controls (OR=2.44; 95%CI, 1.01-5.96; *P*=0.0492) (Supplementary Figure 3 and Supplementary Table 6).

**Figure 3:**
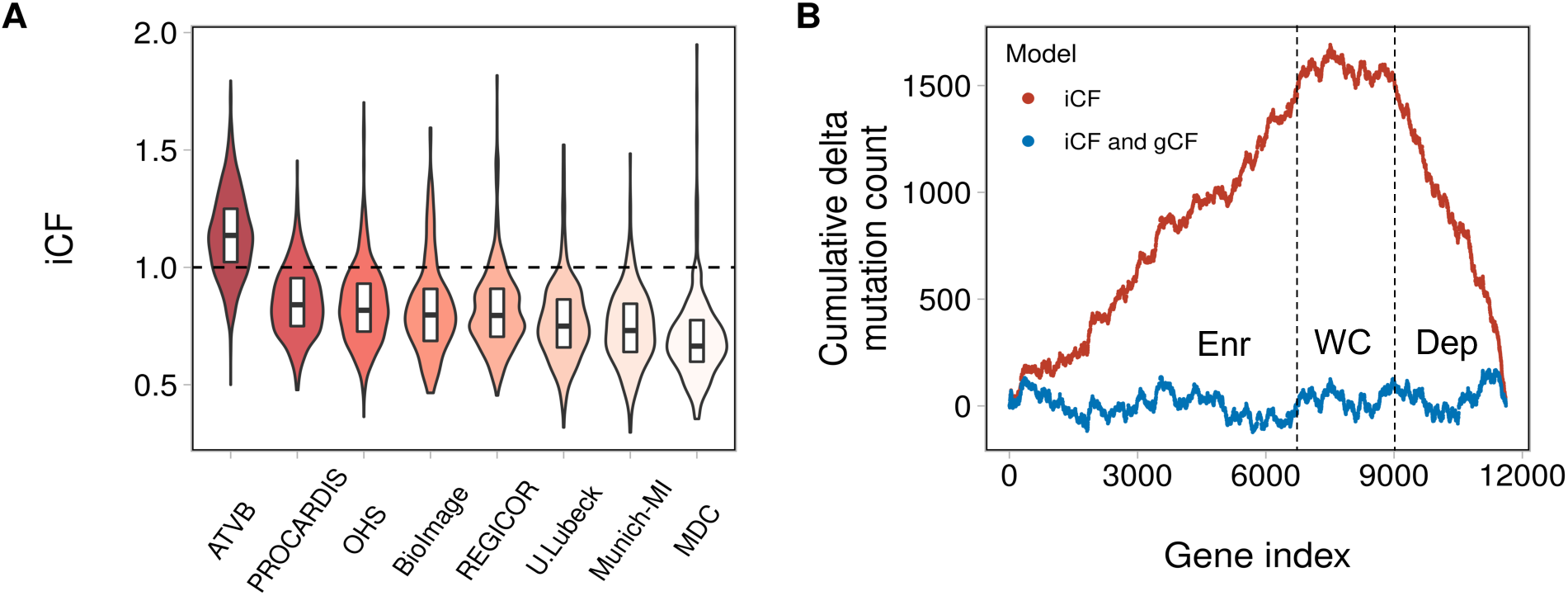
Correction factor adjustment between studies and across genes. Distribution of iCFs for 6,082 healthy controls from 8 cohorts in MIGen were determined across all protein-coding genes **(A).** Dashed line represents an iCF corresponding to equal mutation loads between a MIGen participant and gnomAD (i.e. iCF = 1). The violins demonstrate the spread of iCF among all 8 MIGen cohorts. The horizontal line in each boxplot indicate the median iCF values while the top and bottom lines represent the 75^th^ and 25^th^ percentiles of the iCF distribution, respectively. Length of boxplot represents the inter-quartile range of iCF values. The cumulative delta mutation counts (i.e. the cumulative difference in mutation count between healthy control participants in MIGen and gnomAD per protein-coding gene) demarcates genes that are systematically enriched (Enr), well-calibrated (WC) and depleted (Dep) for rare pathogenic alleles among MIGen control participants **(B)**. The height of the mountain (red) demonstrates the degree of adjustment offered by the iCF in order to achieve calibration between MIGen controls and gnomAD. Implementing the gCF (blue) mitigates residual gene-level biases that cannot be accounted for by the iCF alone. iCF indicates individual correction factor, and gCF indicates gene correction factor.

### RV-EXCALIBER adequately calibrates associations using healthy controls participants from MIGen

The efficacy of the gCF is largely determined on the degree of similarity between a test and ranking cohort in terms of sequencing technology and population structure. In order to evaluate the adequacy of gCF adjustment in the absence of disease associations, we randomly divided healthy controls from MIGen (N=6,082) into a ranking cohort (N=2,730) in which the gCF would be calculated based on the distribution of gene-based *P*-values and expected counts and a test cohort (N=3,352) in which the gCF would be applied (see Data Supplement). 11,616 genes with observed and expected counts of ≥1 were allocated into 50 distinct gCF bins (see Data Supplement), corresponding to 232 genes per bin. gCF values ranged from 0.89 (bin 30) to 1.25 (bin 10) in the ranking cohort, with a significant correlation between bin index and gCF. In other words, genes that were depleted and enriched for pathogenic alleles in gnomAD trended toward higher and lower gCF values, respectively (R^2^=0.57; *P*<0.0001) (Supplementary Figure 4). Adjustment for gCF further attenuated deviations in observed versus expected counts as compared to what is achievable using the iCF alone (Figure 3B).

After establishing the iCF and gCF adjustments, we sought to compare the genomic inflation factors (i.e. lambda) at the median (*λ_med_*) using the RV-EXCALIBER and Testing Rare variants Against Public Data (TRAPD)^15^ methods in order to assess the presence of type I error. For TRAPD, inclusion of rare pathogenic nonsynonymous SNVs was based on the top 94^th^ percentile of quality-by-depth (QD) score according to rare synonymous SNVs in 3,352 healthy control exomes in MIGen and top 90^th^ percentile of QD scores in gnomAD, which achieved the best calibration between the 95^th^ percentile test statistics (*λ*_95_) and a uniform distribution of *P*-values (*λ_95_* = 0.99) (Supplementary Figure 7). Inclusion of insertions and deletions were based on the top 96^th^ percentile QD scores for all rare nonsynonymous SNVs in 3,352 healthy control exomes in MIGen and top 90^th^ percentile of QD scores in gnomAD (*λ*_95_ = 1.02). After applying TRAPD and RV-EXCALIBER to 4,160 protein coding genes that had observed counts ≥1 and expected counts ≥10 using both methods (to forego potential biases due to genes with low power to detect rare variant associations), RV-EXCALIBER achieved better calibration at the median of test statistics (*λ_med_* = 0.95 and 0.41 for RV-EXCALIBER and TRAPD, respectively) (Figure 4).

**Figure 4:**
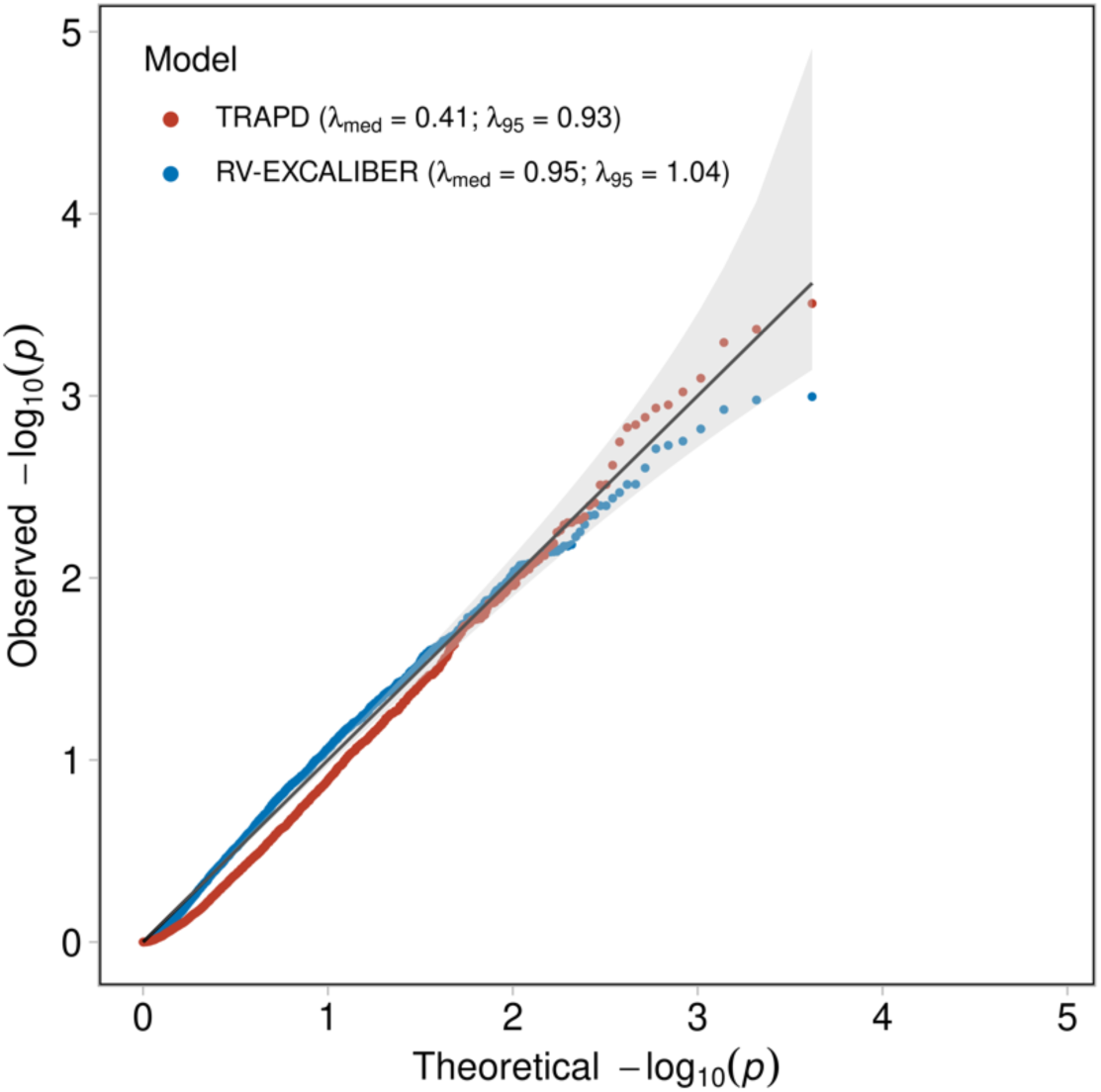
Quantile-quantile plot for gene-based test statistics generated using TRAPD and RV-EXCALIBER for 3,352 healthy control participants in MIGen. A total of 4,160 genes with observed counts ≥1 in MIGen and expected counts ≥10 in gnomAD non-Finnish Europeans were used to conduct gene burden testing according to the TRAPD and RV-EXCALIBER methods. To gauge calibration of test statistics between methods, 3,352 healthy MIGen control participants were compared to gnomAD non-Finnish Europeans. The remaining subset of 2,730 MIGen healthy controls were used as the gCF ranking cohort. The genomic inflation estimates were evaluated at the median (*λ_med_* and 95^th^ percentile (*λ*_95_) of test statistics. The solid black line indicates an expected uniform distribution of *P*-values under the null and the shaded region represents the 95% confidence interval of the expected distribution.

We next sought to assess the contribution of each of the three proposed adjustments to type I error control using the same 3,352 healthy controls in MIGen as “cases” and gnomAD non-Finnish Europeans as controls. Genomic inflation estimates at the median were better calibrated when using RV-EXCALIBER base (which only accounts for LD between rare variants and does not incorporate ‘iCF’ or ‘iCF and gCF-adjustment to the expected counts) compared to a Fisher’s Exact test across 4,815 protein-coding genes (*λ_med_* = 0.69 and 0.57, respectively). We also demonstrated that calibration of test statistics improves in a step-wise fashion upon incorporating each additional feature of RV-EXCALIBER. Specifically, *λ_med_* estimates incrementally approach 1 using i) RV-EXCALIBER base, ii) RV-EXCALIBER base + iCF adjustment, and iii) RV-CALIBER base + iCF + gCF adjustment (i.e. the fully adjusted model) (Supplementary Figures 5 and 6).

### Rare variant association

After adjusting gnomAD expected counts with RV-EXCALIBER, we conducted an exome-wide rare variant association on protein-coding genes across 5,910 discovery MI cases, testing all 7,146 genes with an observed count ≥1 and an expected count of ≥10 (again, to filter out underpowered genes). RV-EXCALIBER adequately calibrates gene-based test statistics (*λ_med_* =0.92) (Figure 5). The low-density lipoprotein receptor (*LDLR*) was identified as the only exome-wide significant signal (OR = 2.25; 95% CI, 1.70-3.05; *P* = 9.4×10^-13^), with an observed allele count of 160 and an adjusted expected count of 71.1. Among the 160 total observed alleles, 111 were located at distinct exonic sites including 99 (89.2%) missense variants, 10 (9.0%) nonsense variants, 1 (0.9%) frameshift insertion, and 1 (0.9%) frameshift deletion (Supplementary Table 7). We also noted a suggestive signal in tyrosine protein kinase Fes/Fps (*FES*) (OR = 2.00; 95% CI, 1.20-3.42; *P* = 2.5×10^-4^) which has previously been identified as a genome-wide significant loci for CAD in multiple studies (Supplementary Table 8). In fact, a total of 35 (excluding *LDLR*) of the 7,146 genes were found to harbor a genome-wide significant variant for CAD (i.e. a CAD gene set). After generating a null distribution of combined test statistics for 100,000 random gene sets of equivalent size, we identified that the probability of observing a combined test statistic greater than or equal to that of the CAD gene set was statistically significant (*P* = 0.014).

**Figure 5:**
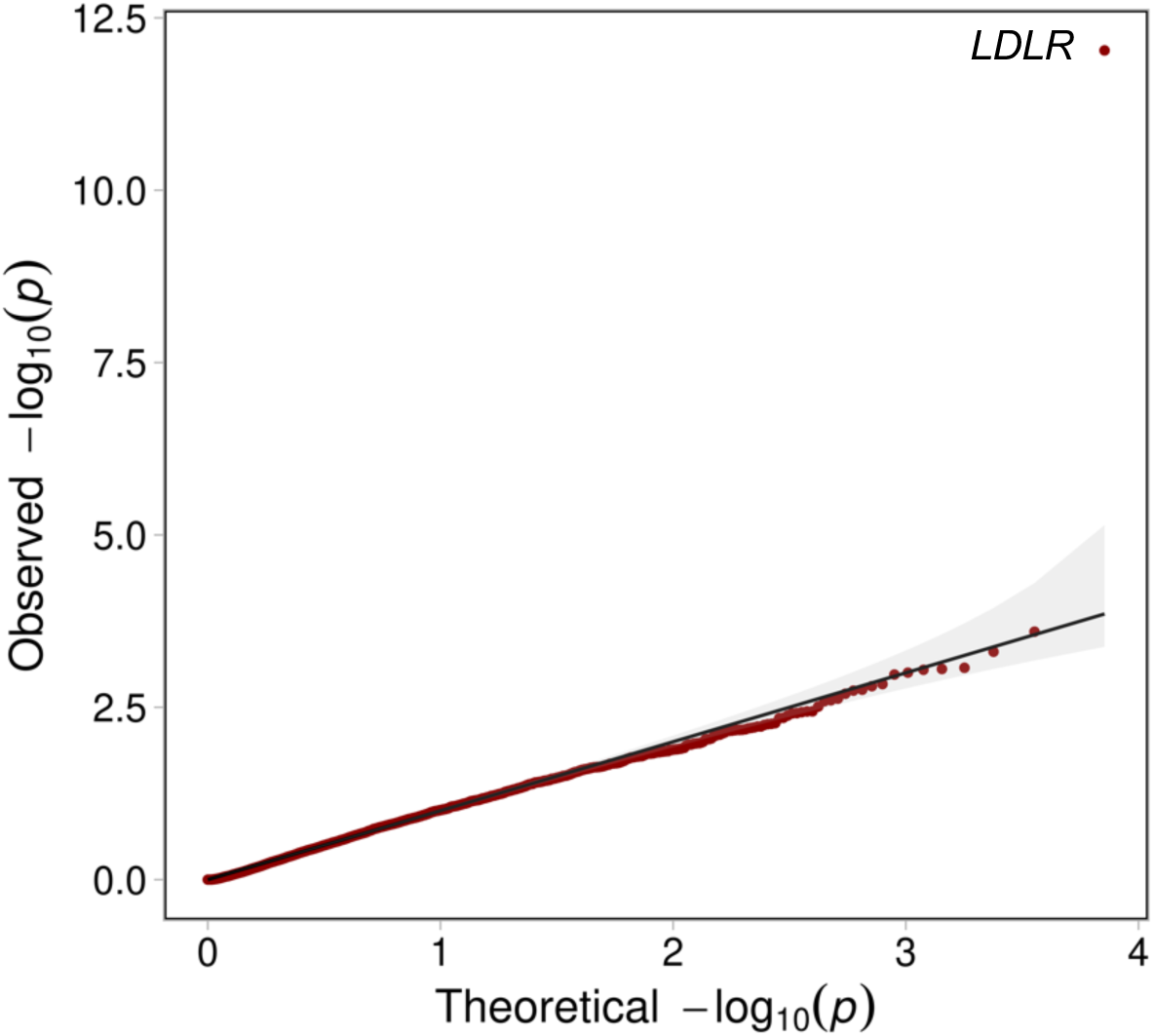
Quantile-quantile plot for discovery gene-based test statistics generated using RV-EXCALIBER for 5,910 case participants from MIGen. A total of 7,146 protein-coding genes with observed counts ≥1 in MIGen and expected counts ≥10 in gnomAD non-Finnish Europeans were used to conduct a gene burden testing with the RV-EXCALIBER method. Association testing was conducted using 5,910 MI cases in MIGen and gnomAD non-Finnish Europeans as controls. The genomic inflation estimate at the median (*λ_med_*) was determined to be 0.92. The solid black line indicates an expected uniform distribution of *P*-values under the null and the shaded region represents the 95% confidence interval of the expected distribution.

### A rare variant gene risk score for prevalent CAD has consistent predictive effects in European and South Asian populations

To test whether a calibrated rare variant gene score (RVGRS) could predict CAD, we calculated RVGRSs that weighted pathogenic alleles using adjusted gene-based effect size estimates from the MIGen rare variant association. Each RVGRS was calculated in each case (N=3,843) and control (N=42,007) in the UK Biobank European participants, incrementally including the top 10 to top 3,000 genes (based on *P*-value) from the MIGen association (increments of 10 genes; 300 total scores). The RVGRS generated from the top 950 discovery genes (RVGRS950) best discriminated participants with CAD from healthy controls, with a 1.08-fold (95% CI, 1.04-1.11; *P*=2.1×10^-5^) increased odds for developing CAD relative to healthy control subjects per 1 SD change in RVGRS (Figure 6 and 7). As a sensitivity analysis, we removed genes known to confer risk for familial hypercholesterolemia (FH) in order to ensure that the predictive estimate of the RVGRS950 was not due to single-gene Mendelian effects. After removing *LDLR* (which was the only FH gene within RVGRS950; i.e. RVGRS950*^LDLR-^*), a consistent predictive effect for CAD (OR=1.07; 95% CI, 1.03-1.10; *P*=1.5×10^-4^) was observed (Figure 6). A major pitfall of common variants gene scores is the lack of transferability across ancestries. Although RVGRS950 did not reach significance in South Asians from the Pakistan Risk of Myocardial Infarction Study (PROMIS) (N=2,946 for cases and N=3,708 for controls), there was still a similar predictive effect observed (OR=1.04; 95% CI, 0.99-1.08; *P*=0.096). Nevertheless, a RVGRS generated using the top 680 genes (RVGRS680) did validate in PROMIS (OR=1.06; 95% CI, 1.01-1.11; *P*=0.024) with a consistent effect in the UK Biobank (OR=1.06; 95% CI, 1.03-1.10; *P*=5.9×10^-4^). In fact, we identified that the effect estimates generated from all RVGRS (i.e. top 10 to top 3000 genes) did not significantly differ between the UK Biobank and PROMIS according to a test for heterogeneity (Figure 7 and Supplementary Table 9). As a negative control, we tested the same 300 RVGRS on UK Biobank participants with irritable bowel disease (N=3,058) and identified no significant difference in distribution of RVGRS between case and control subjects.

**Figure 6:**
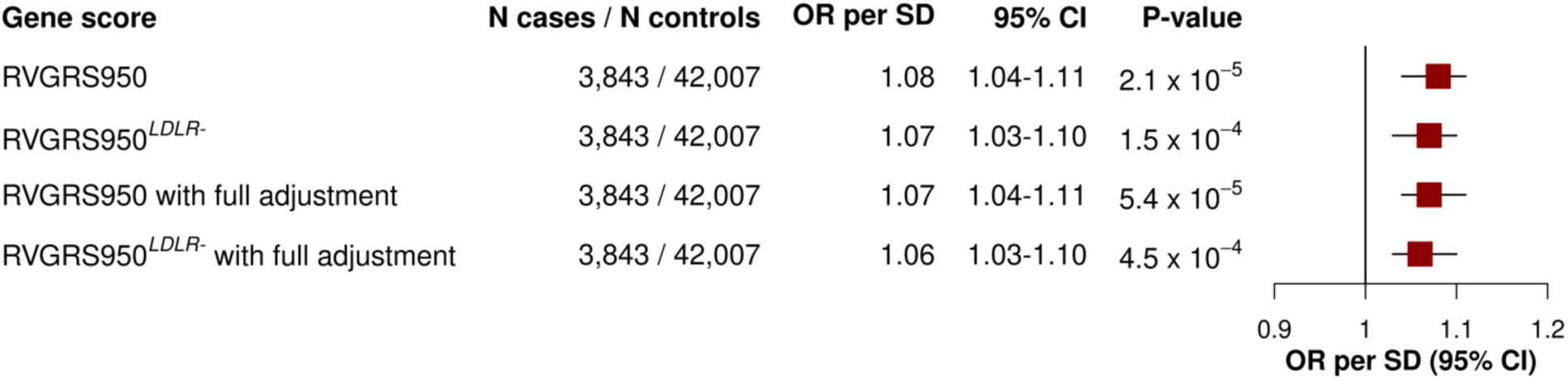
Effects of RVGRS950 and RVGRS950*^LDLR-^* on CAD among UK Biobank participants. Predictive effects of RVGRS950 and RVGRS950*^LDLR-^* (i.e. RVGRS950 without *LDLR*) on CAD are shown for UK Biobank European participants. The same RVGRS with additional adjustment for clinical risk factors and CVGRS (i.e. with full adjustment) are also shown. Odds ratios are expressed in terms of a 1 SD change in RVGRS and were calculated using logistic regression. All RVGRS associations are adjusted for age, age^2^, sex, and the first 20 principal components of ancestry. 95% confidence intervals were calculated from the standard error of the beta regression estimate. OR indicates odds ratio, CI indicates confidence interval, and CVGRS indicates common variant genetic risk score.

**Figure 7:**
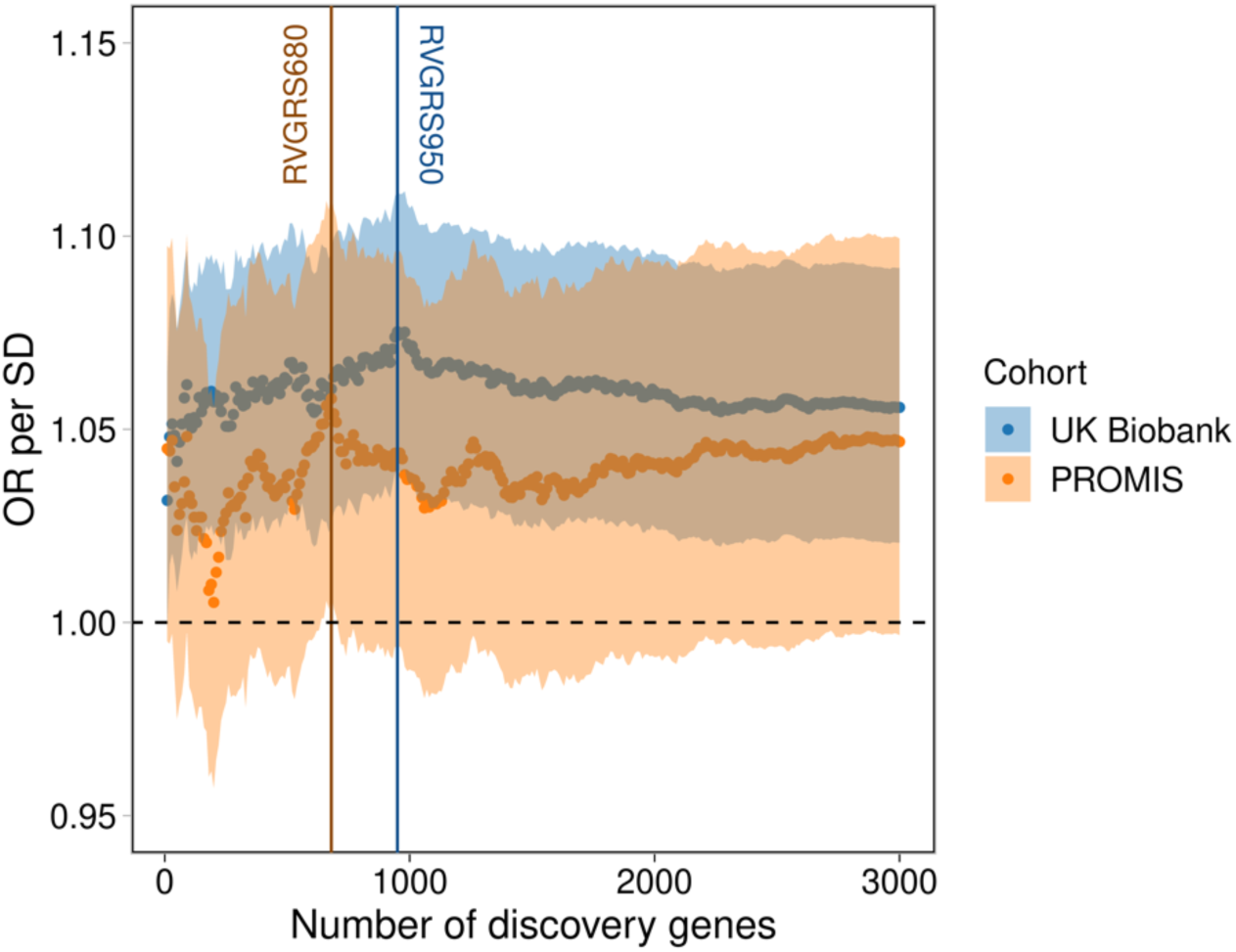
RVGRS effect estimates on CAD evaluated in UK Biobank Europeans and PROMIS South Asians. Predictive effects of 300 RVGRSs generated using the top 10 to top 3000 discovery genes (in intervals of 10) on CAD are shown for Europeans in UK Biobank and South Asians in PROMIS. RVGRS950 and RVGRS680 correspond to the RVGRS that significantly achieved the highest predictive estimate in the UK Biobank Europeans and PROMIS South Asians, respectively. Shaded regions correspond to 95% confidence intervals of effect estimates. Dashed line represents a line of no effect. Odds ratios are expressed in terms of a 1 SD change in RVGRS and were calculated using logistic regression adjusted for age, age^2^, sex, and the first 20 principal components of ancestry. 95% confidence intervals were calculated from the standard error of the beta regression estimate. OR indicates odds ratio and CI indicates confidence interval.

### RVGRS acts independently of common variant gene scores and clinical risk factors

RVGRS950 and RVGRS950*^LDLR-^* acted independently from both the aggregate effect of common variants and clinical risk factors, with a consistent predictive estimates (OR=1.07; 95% CI, 1.04-1.11; *P*=5.4×10^-5^ for RVGRS950 and OR=1.06; 95% CI, 1.03-1.10; *P*=4.5×10^-4^ for RVGRS950*^LDLR-^*) (Figure 6). Moreover, we observed a significant trend of higher RVGRS950 estimates among individuals with lower clinical risk for CAD (*P*interaction=0.015), where individuals in low, middle, and high tertiles of FRS had ORs of 1.16, 1.10, 1.03 conferred through RVGRS950 for CAD, respectively. The observed trend also remained significant when using RVGRS950*^LDLR-^* (Supplementary Table 10). Interestingly, the effect conferred through CVGRS did not significantly differ across FRS tertiles, with ORs of 1.20, 1.14, and 1.16 for low, middle, and high tertiles of FRS, respectively (*P*_interaction_=0.65) (Supplementary Table 11). We next tested whether addition of RVGRS950 could improve patient classification as compared to a base model consisting of only CVGRS and FRS. Incorporation of RVGRS950 was found to significantly improve risk prediction for CAD (net reclassification index=0.0571; *P*=9.4×10^-4^) after adjusting models for age, age^2^, sex and the first 20 principal components of ancestry. The net reclassification index also remained significant when using RVGRS950*^LDLR-^* (Supplementary Table 12).

### RVGRS demonstrates increased predictiveness for early-onset CAD

Due to the substantial heritability of early-onset CAD and the enrichment of mendelian of FH mutations in these subjects, we tested the predictive effect of RVGRS950*^LDLR-^* across 4 age categories. These included males and females with established onset of CAD or MI at age i) ≤40 and ≤45, ii) ≤45 and ≤50, iii) ≤50 and ≤55, or iv) ≤55 and ≤60, respectively. RVGRS950*^LDLR-^* was found to confer the highest risk among individuals with the earliest onset of CAD, with a 1.32-fold (95% CI, 1.01-1.69; *P*=0.035) increased odds of disease per 1 SD change in RVGRS (Figure 8). The median RVGRS950*^LDLR-^* in early CAD cases corresponded to the 66^th^ percentile of RVGRS950*^LDLR-^* for the control distribution, compared to only the 52^nd^ percentile for general CAD (Figure 9). 1 in 68 (1.5%) of participants in the UK Biobank had a RVGRS950*^LDLR-^* score corresponding to a ≥2-fold increase in odds of early disease.

**Figure 8:**
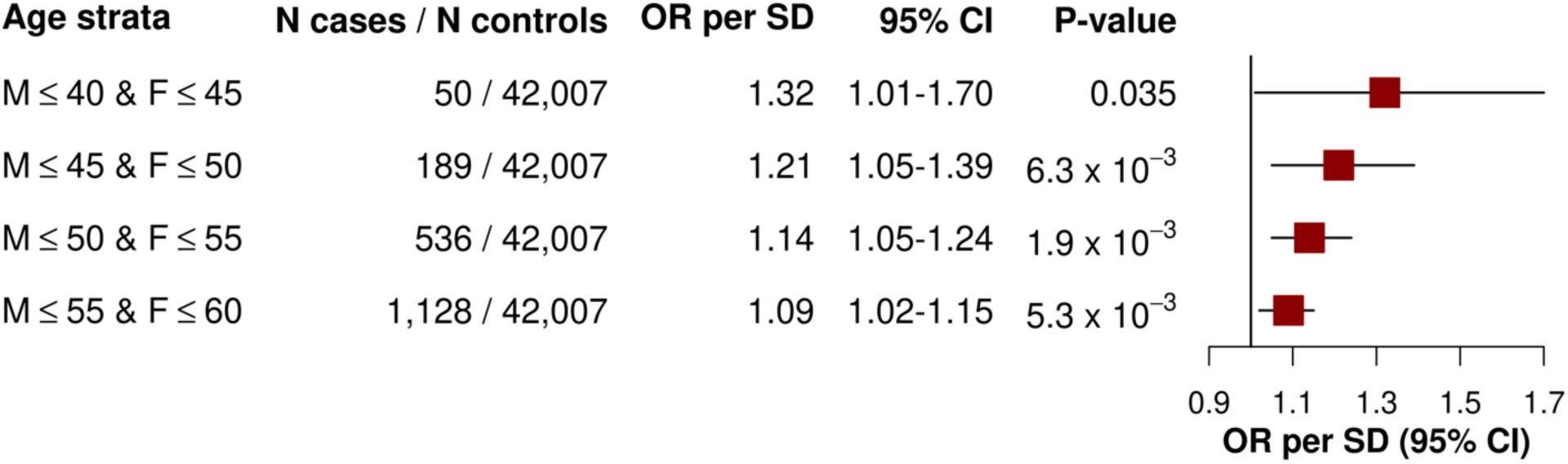
Effects of RVGRS950*^LDLR-^* among 4 age strata in UK Biobank participants. Predictive effects of RVGRS950*^LDLR-^* are shown for UK Biobank European participants across 4 age strata. Ages correspond to earliest event across all pertinent ICD-10 codes that encompassed our CAD definition. Odds ratios are expressed in terms of a 1 SD change in RVGRS and were calculated using logistic regression adjusted for sex, and the first 20 principal components of ancestry. 95% confidence intervals were calculated from the standard error of the beta regression estimate. OR indicates odds ratio, CI indicates confidence interval, M indicates male, and F indicates female.

**Figure 9:**
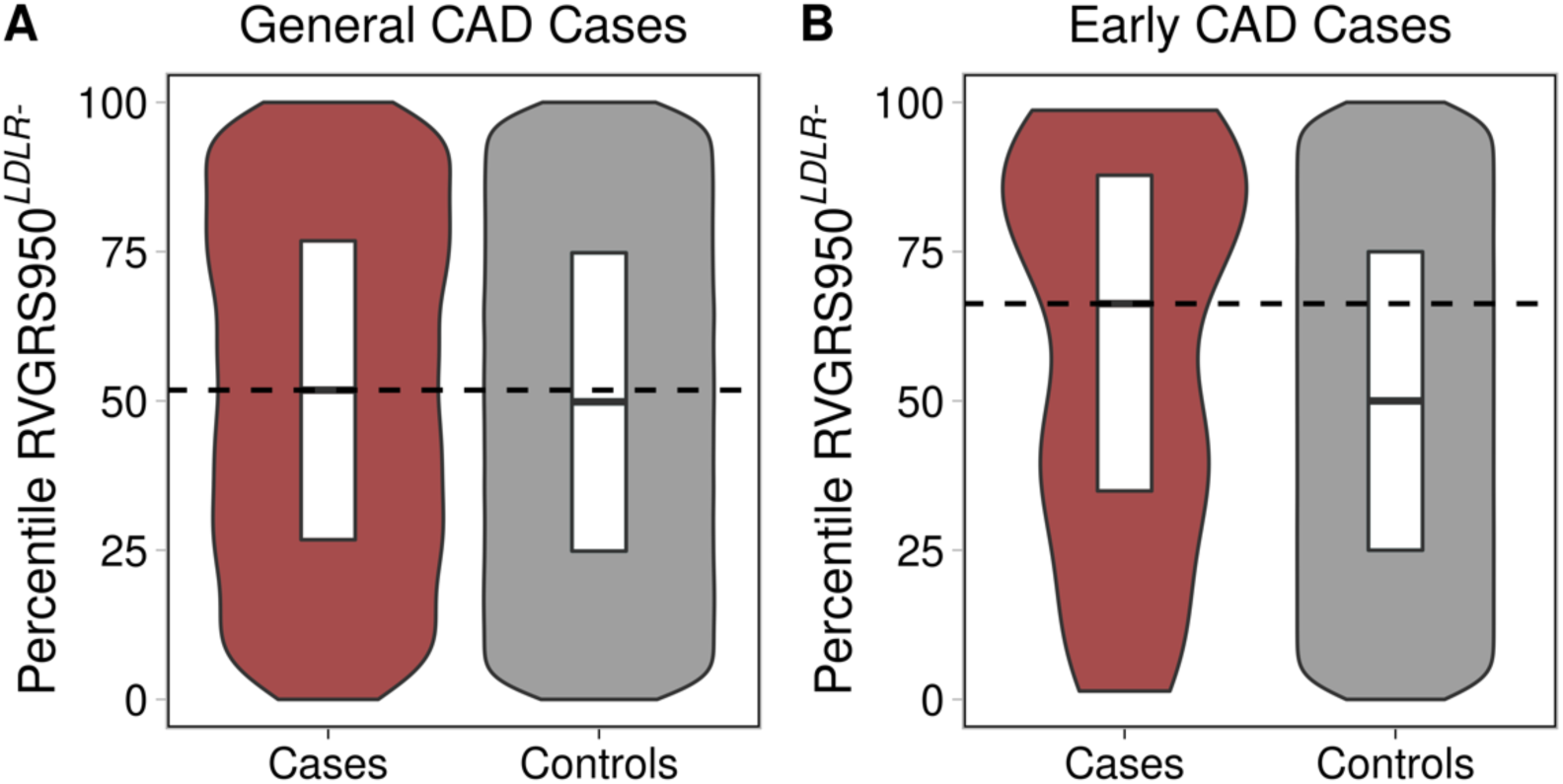
Distribution of RVGRS950*^LDLR-^* among CAD-free controls, general CAD cases, and early CAD cases in the UK Biobank. Distribution of RVGRS950*^LDLR-^* was evaluated for UK Biobank control participants and UK Biobank case participants with general CAD (i.e. any CAD irrespective of age) **(A)**, as well as UK Biobank control participants and UK Biobank participants with early CAD (i.e. males ≤ 40 and females ≤ 45) **(B)**. The violins demonstrate the spread of RVGRS950*^LDLR-^* distribution for these groups. The horizontal line in each boxplot indicate the median RVGRS950*^LDLR-^* value while the top and bottom lines represent the 75^th^ and 25^th^ percentiles of the RVGRS950 distribution, respectively. Length of boxplot represents the inter-quartile range of RVGRS950*^LDLR-^* values. The dashed lines represent where the median of each respective case group falls in the control distribution.

## DISCUSSION

We herein describe a novel framework, which leverages summary allele frequencies in gnomAD to identify complex disease risk conferred through the modest effects of rare variant burden. While methods have been developed to address the genetic contributors to Mendelian disease using external control databases, there is currently no workflow to systematically calibrate rare variant burden between test samples and reference databases. RV-EXCALIBER uses individual and gene-level mutation loads to calibrate exome sequences to summary-level allele frequencies in gnomAD while also accounting for LD between rare variants. In doing so, our method addresses the primary sources of bias that confound association signals generated using a technically and phenotypically heterogeneous reference population. By instituting our corrections, we were able to leverage a very large control sample to detect modest rare variant effects which were used to generate a rare variant gene risk scores for CAD.

Current methodologies that leverage gnomAD as controls have been tailored to demarcate variant and gene-based associations for Mendelian disorders, which are mediated through mutations with very high effect^15,16^. There is currently no framework that incorporates global and granular adjustments to gnomAD in order to detect rare variant burden associated with complex diseases, which are primarily conferred through variants of modest effects. Furthermore, existing methodologies utilizing gnomAD as controls, such as TRAPD, calibrate test statistics based on benign mutation rate by altering filtering criteria for variant inclusion^15^. While this procedure has shown to be effective in this respect^15,16^, informing variant inclusion through quality metrics such as quality-by-depth and variant quality score recalibration does not necessarily control for the major sources of bias when using public controls such as gnomAD. Such biases include 1) differences in individual-level mutation load between case subjects and gnomAD, which reflect disparities in sequencing technology and population substructure and 2) bias among genes that demonstrate systematic enrichment or depletion relative to gnomAD. In fact, when conducting a rare variant association test, RV-EXCALIBER demonstrated better calibration of gene-based test statistics relative to TRAPD according to genomic inflation estimates calculated at the median (Figure 4).

We show through simulations that individual-level differences in rare variant mutation burden can be impacted by variation in population substructure (i.e. PSF) and technological biases (i.e. SFP and SFN). Detection of rare variants through exome sequencing can vary substantially based on methods of library preparation, exome enrichment strategy, read alignment ^17^. To evaluate the effect of population factors, we used three consensus sequences from GIAB and compared to closely matched ethnic groups in gnomAD. For each comparison, the iCF deviated most from expectation in rare variant bins, especially among European populations (Figure 2). This is largely because rare variants demonstrate tremendous geographical disparity as they have either arisen recently or are older variants that are functionally deleterious and so kept at low population frequency by purifying selection^11,18–20^. As such, individual rare variants tend to cluster in smaller population groups and are not geographically ubiquitous like common variants. This is especially true in Europe, as principal component analysis has revealed consistent and distinct clustering of sub-populations^21,22^. Indeed, we show that the mutation burden in European consensus sequences (i.e. Central European or Ashkenazi) deviate substantially from the Non-Finnish Europeans in gnomAD and is more closely matched across common variants (Figure 2A & 2C). Surprisingly, differences in iCF between cases and controls were observed even within single studies. This can likely be attributed to batch effect sequencing, which provides further rationale to incorporate correction when comparing exomes to reference datasets, especially in case-control designs where samples are potentially sequenced in batches based on affection status. Although the iCF accounts for global deviation in sequencing and population bias, we instituted the gCF to specifically address the bias in mutation burden at the gene-level, which cannot otherwise be captured on an exome-wide scale. Specifically, by tracking the cumulative exome-wide mutation load, we demonstrate that the gCF mitigates bias in genes harbouring residual inflation or deflation that could not be addressed using iCF alone for global adjustment. Since genes were ranked according to both *P*-value and expected count in an independent set of healthy controls, we demarcate genes that are reproducibly “enriched”, “well-calibrated”, and “depleted” in rare mutations in relation to gnomAD non-Finnish Europeans (Figure 3B). Overall, our results validate presence of systematic gene biases specific to a given cohort and effectively illustrates the extent of adjustment. It is important to note that both iCF and gCF are implemented on our base method that adjusts for LD among rare variants, since LD cannot be estimated by summary-level data in gnomAD. Alternative methods do not adjust for LD when using gnomAD as controls^15,16^.

Common variant gene scoring approaches have limited trans-ancestral predictive value and clinical implications remain highest when discovery and validation sets are ethnically matched, which is largely the case only for European populations^18,19,23^. Moreover, empirical and simulation models have demonstrated that directional selection across ancestries can severely impair predictive accuracy in non-European validation populations, even after accounting for principal components and proportion of causal variants^18,19^. The large disparity in linkage-disequilibrium (LD) structure between ethnic groups markedly impair the predictive value of common variant GRS since causal variants remain largely unknown and thus tag variants are used. In contrast, rare variants are more likely to be functional and thus execute the same biological function regardless of ancestral origin. We show that the RVGRS demonstrates consistent predictive estimates for Europeans in UK Biobank exomes and South Asian exomes in PROMIS (Figure 7).

In order to concretely establish the RVGRS as an independent mediator of disease, we verified that the predictive estimate elicited through RVGRS remained consistent even after adjusting for CVGRS as derived using effect estimates from the CARDIoGRAMplusC4D consortia and clinical risk factors as calculated by a Framingham risk score^22,24^. In fact, adding RVGRS to a reference model consisting of CVGRS and clinical risk factors significantly reclassified 5.7% of CAD events, which represents a marked improvement given the relatively small size of our discovery population. Moreover, the RVGRS demonstrated an increase in predictive effect for individuals at low clinical risk as compared to those with a high clinical burden for developing CAD. Interestingly, no such effect gradient was observed for the CVGRS, which mitigates its ability to discern residual genetic risk for CAD among individuals that may present with limited clinical risk factors. Similar to CVGRS, we found that the risk conferred through RVGRS was strongest among very young individuals (i.e. males ≤40 and females ≤45) and that the risk markedly decreased upon inclusion of older individuals (Figure 9), which suggests a higher burden of rare variant associations in early-onset cases. This is consistent with earlier evidence which shows that early-onset cases have a higher genetic burden of disease due to limited exposure to traditional environmental risk factors^25–29^.

It is important to note that the risk estimates for high polygenic risk derived from RVGRS and CVGRS cannot be readily compared due to the extreme discrepancy in discovery sample size used to compute effect estimates to weight alleles in validation samples. In our work, we included 5,910 cases to derive beta estimates while the betas used for CVGRS in CAD/MI are obtained from the CARDIoGRAMplusC4D meta-analysis, which has a case population that is 10.3x larger. Nevertheless, our work bridges the gap between rare and common variants under a unified polygenic framework. Despite the marked difference in case sample size, we show that a RVGRS confers high risk (i.e. ≥2-fold) for early CAD in 1.5% of the population and remains independent of Mendelian effects, clinical risk factors, and CVGRS.

We herein present a method (RV-EXCALIBER) for calibrating rare pathogenic alleles between case exomes and gnomAD by mitigating major biases at the individual and gene-level while also correcting for putative LD between rare variants. RV-EXCALIBER demonstrates better calibration than existing methodologies that leverage gnomAD as controls and was empirically applied to generate a first RVGRS for CAD. The RVGRS conferred disease risk independent of Mendelian effects, clinical risk factors, and CVGRS while also correctly reclassifying ∼6% of CAD events. Our results suggest that addition of RVGRS to CVGRS and Mendelian mutations has potential clinical utility. Our results also make a strong case for large sequencing studies of common, complex traits and diseases. Additionally, leveraging publicly available data as controls in an unbiased manner may permit additional resources to be allocated to recruiting extreme disease cases, which might harbor a greater burden of rare causal variants. Lastly, we also note that RV-EXCALIBER could be extended to whole-genome sequencing, in order to better delineate the role of rare non-coding variants on disease risk.

## Acknowledgments

This research has been conducted using the MIGen and UK Biobank resources.

## Supplementary Appendix

### 1. Data acquisition and discovery study samples

#### A. genome Aggregation Database (gnomAD)

The gnomAD dataset consists of 120,393 exomes which were aggregated across 35 exome sequencing consortia^1^. Release 2.0.1 of the gnomAD dataset (gnomAD r2.0.1) was used as the reference population against all comparator sequences (described in **Supplementary Sections 1B and 1C**). After data processing (discussed in **Supplemental Sections 4A, 4B, and 4C**), there were a total of 13.1 million variant calls (93.6% single nucleotide variants (SNV); 6.4% insertions/deletions (INDEL)).

#### B. Genome In A Bottle (GIAB) consensus sequences

The GIAB consortium is hosted by the National Institute of Standards and Technology (NIST) to provide gold-standard reference samples for benchmarking human genomic sequences^2^. To evaluate the role of population-specific effects to exome-wide mutation load, we determined correction factors using high-confidence sequence variants corresponding to 3 reference GIAB samples of different ethnicities (using methodology developed by Zook *et al.* 2014)^2^ : 1) NA12878 (Central European; NIST ID HG001), 2) NA24631 (East Asian; NIST ID HG005), and 3) NA24385 (Ashkenazi Jewish; NIST ID HG002). Variants called in these reference samples were harmonized across 5 sequencing technologies, 7 read mappers and 3 variant callers to generate a “consensus” variant callset that we used as benchmark sequences to evaluate the excess or deficit in mutation load across all protein-coding genes in the gnomAD database. Consensus variant calls also allows for the identification of “difficult-to-sequence” regions across the genome, such as segmental duplications, short tandem repeats and other structural variations. Exclusion of these regions can subsequently be used to generate “high-confidence” genomic intervals. High confidence region files and variant call sets (in variant call format) for version 3.2.2 of NA12878, version 3.3 of NA24631, and version 3.3.2 of NA24385 were obtained from the GIAB ftp repository (ftp://ftp-trace.ncbi.nlm.nih.gov/giab/ftp/release/NA12878_HG001/NISTv3.3.2/), (ftp://ftp-trace.ncbi.nlm.nih.gov/giab/ftp/release/ChineseTrio/HG005_NA24631_son/NISTv3.3/), and (ftp://ftp-trace.ncbi.nlm.nih.gov/giab/ftp/release/AshkenazimTrio/HG002_NA24385_son/latest/GRCh37/) for use in our benchmarking analysis.

#### C. Myocardial Infarction Genetics (MIGen) sequences

MIGen exome sequencing datasets were obtained from the database of Genotypes and Phenotypes (dbGaP). All data handling and analyses were approved by the Hamilton Integrated Research Ethics Board and approval for data downloads was subsequently granted by the National Heart, Lung, and Blood Institute Data Authorization Committee (NHLBI DAC). Variant call files (VCF’s) and general phenotypic information were downloaded with authorized access using version 2.9 of NCBI’s Sequence Read Archive (SRA) toolkit^3^. All dbGaP study accessions used in this work are listed in Supplementary Table 1 for reference. A total of 10 MIGen cohorts were downloaded from dbGaP. Summary information on each cohort is provided in Supplementary Table 1.

**Supplementary Table 1:** Baseline and sequencing characteristics for individual cohorts used from the MIGen consortium.

| MIGen Cohort | dbGaP study accession | Ethnicity | % Reported Male | Sequencing platform | Exome enrichment kit | Post-QC Cases (N) | Post-QC Controls (N) | % Cases retained after QC | % Controls retained after QC |
| --- | --- | --- | --- | --- | --- | --- | --- | --- | --- |
| Italian Atherosclerosis Thrombosis and Vascular Biology | phs000814.v1.p1 | European | 89 | HiSeq 2000 and 2500 | SureSelect Human All Exon v.2 Kit | 1784 | 1719 | 98 | 97 |
| Malmo Diet and Cancer Study | phs001101.v1.p1 | European | 55 | HiSeq 2000 and 2500 | ICE Capture Reagent | 518 | 522 | 96 | 97 |
| BioImage Study | phs001058.v1.p1 | European | 61 | HiSeq 2000 and 2500 | ICE Capture Reagent | 128 | 269 | 84 | 77 |
| University of Lubeck | phs000990.v1.p1 | European | 61 | HiSeq 2000 and 2500 | ICE Capture Reagent | 799 | 859 | 92 | 96 |
| German Heart Centre in Munich | phs000916.v1.p1 | European | 65 | HiSeq 2000 and 2500 | ICE Capture Reagent | 386 | 394 | 97 | 99 |
| Registre Gironi del Cor | phs000902.v1.p1 | European | 77 | HiSeq 2000 and 2500 | SureSelect Human All Exon v.2 Kit | 357 | 390 | 93 | 97 |
| Precocious Coronary Artery Disease Study | phs000883.v1.p1 | European | 65 | HiSeq 2000 and 2500 | SureSelect Human All Exon v.2 Kit | 983 | 959 | 95 | 94 |
| Ottawa Heart Study | phs000806.v1.p1 | European | 67 | HiSeq 2000 and 2500 | SureSelect Human All Exon v.2 Kit | 955 | 970 | 97 | 98 |
| Pakistan Risk of Myocardial Infarction Study | phs000917.v1.p1 | South Asian | 83 | HiSeq 2000 and 2500 | ICE Capture Reagent and SureSelect Human All Exon v.2 Kit | 2946 | 3708 | 97 | 92 |

### 2. Study sample descriptions for MIGen

**Supplementary Table 2:** Details on overall study design along with pertinent inclusion criteria for cases and controls for each MIGen exome sequencing cohort.

| MIGen Cohort | Study design | Case inclusion criteria | Control inclusion criteria | Reference |
| --- | --- | --- | --- | --- |
| *Discovery* |  |  |  |  |
| Italian Atherosclerosis Thrombosis and Vascular Biology | Case-control | Men and women with MI < 45 years old with coronary angiography | Men and women without history of thromboembolic disease; matched to cases according to age, sex, and geographical origin | 4,5 |
| Malmo Diet and Cancer Study | Nested case-control | Men and women with incident non-fatal or fatal MI | Men and women with no MI on follow-up | 6,7 |
| BioImage Study | Nested case-control | Men and women that developed cardiovascular events (defined as myocardial infarction or coronary revascularization) after a median follow-up of 2.7 years from baseline | Men and women with no cardiovascular events (defined as myocardial infarction or coronary revascularization) on follow-up; matched to cases for age (+/- 2 years). Sex, smoking status, and type 2 diabetes status | 8,9 |
| University of Lubeck | Case-control | Men and women with MI < 60 years old and at least 1 first-degree relative with coronary artery disease | Consisted of a random set of individuals from the German population that met a similar age and sex proportion as cases, but unselected for phenotype | 10–13 |
| German Heart Centre in Munich | Case-control | Men with MI at < 41 years old and women with MI at < 56 years old | Men > 65 years old and women > 75 years old without personal history of MI or CAD | 14 |
| Registre Gironi del Cor | Nested case-control | Men with MI at < 50 years old; women with MI at < 60 years old | No personal history of MI or coronary revascularization between 55 and 80 years old | 15,16 |
| Precocious Coronary Artery Disease Study | Case-control | Men or women with MI, unstable angina, stable angina, or coronary revascularization at < 66 years old | No personal or sibling history of MI, unstable angina, stable angina, or coronary revascularization at < 66 years old | 17,18 |
| Ottawa Heart Study | Case-control | Men < 55 years old and women < 65 years old with personal history of CAD; CAD was defined as having an MI, coronary artery bypass surgery, or angiographic stenosis of > 50% in at least one major coronary artery | Men ≥ 65 years old and women ≥ 70 years old with no peroneal history of cardiovascular disease | 19,20 |
| *Validation* |  |  |  |  |
| UK Biobank 50,000 exomes | Prospective cohort | Definition for CAD provided Supplementary Section 7A | Definition for CAD-free controls provided in Supplementary Section 7A | 21,22 |
| Pakistan Risk of Myocardial Infarction Study | Case-control | Men and women aged 30 to 80 presenting with clinical symptomology consistent with MI, ECG changes consistent with MI, elevated troponin levels consistent with MI, and no history of cardiovascular disease | No self-reported history of cardiovascular disease | 23,24 |

### 3. Broad exome sequencing

All exome sequencing was performed at the Broad Institute of Harvard and MIT. Sample sequence capture chemistry and sequencing platforms are stated for each MIGen cohort in Supplementary Table 1. All methodology pertaining to **1)** QC of sample DNA, **2)** exome sequencing, **3)** library construction and in-solution hybrid selection, **4)** preparation of libraries for cluster amplification and sequencing, **5)** cluster amplification and sequencing, **6)** read mapping and variant analysis, **7)** sequencing QC, or **8)** variant calling has been extensively described in previous works^25–27^.

### 4. Variant-level QC for MIGen and gnomAD

#### A. Data preparation

All multi-allelic variants were broken into separate records using vcflib’s vcfmultibreak tool^28^ and were treated as separate variants. All INDELS variants were left-aligned using the bcftools norm function to standardize reference and alternate allele INDEL calls according to the hg19 reference genome^29^.

#### B. Filter sites

Only sites receiving a filter notation of “PASS” were kept in downstream analyses. As such, sites failing any variant quality metrics (**Supplementary Section 3**) were eliminated.

#### C. Allele counts

Only variant sites demonstrating at least one heterozygous carrier (i.e. an allele count of ≥ 1 were retained.

### 5. Variant-level QC for only MIGen

#### A. Hardy-Weinberg Equilibrium

Variants deviating significantly from Hardy-Weinberg equilibrium (P < 5 x 10^-6^) were removed from the analysis. It is important to note that variants considered for Hardy-Weinberg equilibrium were processed through variant level QC described in **Supplementary Sections 4A, 4B and 4C.**

#### B. Variant missingness

No variant-level missingness threshold was applied as variants exhibiting high missingness would implicitly be removed using the internal allele-frequency filter as discussed in **Supplementary Section 8E**.

### 6. Sample-level QC

#### A. Sample missingness

Rate of missing genotype calls were determined for each sample in each MIGen cohort. Since the distribution of missingness can vary between studies due to batch effects, samples exhibiting missingness > 6 SDs from the mean within each MIGen cohort were removed.

#### B. Sex check

Sex check was performed in plink version 1.9 ^30^ on a cohort-by-cohort basis. Method-of-moments F coefficients based on observed and expected homozygosity counts in the X-chromosome were calculated after removing the pseudo-autosomal regions. Reported females demonstrating F coefficients greater than 6 SDs and reported males demonstrating F coefficients less than 6 SDs were removed from the analysis.

#### C. Ethnicity check

Using the genetic complex trait analysis (GCTA) tool^31^, principal component analysis was conducted with common variants (minor allele frequency (MAF) >0.01) passing quality control from each study. Principal components were projected onto the backdrop of 1000Genomes phase 3 samples, corresponding to reference clusters of African (N=661), East Asian (N=504), European (N=503), South Asian (N=489), and Latin American (N=347) ancestry. Scatter plots of the first two principal components were inspected to identify samples with discrepant reported vs. genetic ethnicity.

#### D. Kinship

Kinship analyses were performed using KING^32^ in each cohort separately. Kinship analysis was automatically stratified by reported ethnic group since no individual cohort had a mixture of reported ancestries. Variants contributing to the kinship analysis were restricted to those that 1) were autosomal 2) were SNV’s 3) were designated ‘PASS’ sites, 4) had a MAF > 0.01, and 5) had < 10% sample missingness. Variants meeting all of the above criteria were further pruned according to linkage disequilibrium. Specifically, r^2^ values were determined in a pairwise fashion for these variants using a window size of 50 and a step size of 5. One variant in a given pair would be removed if the r2 value exceeded 0.2. According to KING documentation, pairs of samples with estimated kinship coefficient ranges of >0.354, [0.177 - 0.354], [0.0884 - 0.177] and [0.0442 - 0.0884] were designated as duplicates/MZ twins, 1^st^-degree relatives, 2^nd^-degree relative, and 3^rd^- degree relatives, respectively. As such, a single sample from a pair with a kinship coefficient ≥0.0442 (that which had less overall missingness) was kept.

#### E. Heterozygosity

A method-of-moments F-coefficient based on the observed and expected homozygous genotypes was determined using plink v1.9^30^ using the same set of variants as used in the kinship analysis. F-coefficients demonstrating values less or greater than 6 SDs from the mean were removed as likely sources of sample admixture and consanguinity, respectively.

### 7. Validation study populations

#### A. UK Biobank exomes

The prospective UK Biobank 50,000 exome release was accessed in March 2019 under application #15255 and contained individual-level genotype data for all individuals who consented for genetic analysis. The pVCF plink binary format files generated using the Functional Equivalent pipeline were downloaded using ukbfetch^33^. The UK Biobank study received approval from the National Health Service National Research Ethics Service North West.

After performing variant and sample-level QC as previously described in **Supplementary Sections 5 and 6**, we achieved 45,850 unrelated subjects of European ancestry (i.e. British and other Caucasians as ascertained by principle component analysis) and 16,051 protein-coding genes with a least singleton variant (as per variant filtering criteria described in **Supplementary Section 5**).

Coronary artery disease was evaluated using a liberal composite outcome consisting of angina, acute and chronic ischemic heart disease, myocardial infarction, or coronary revascularization. Angina pectoris, ischemic heart disease, and myocardial infarction was based on either self-report or hospital admission diagnosis. All phenotypes were defined according to ICD-10 definitions (UK Biobank data field 41270). Angina pectoris was ascertained using ICD-10 codes I20.X. Acute and chronic ischemic heart disease was ascertained using ICD-10 codes I24.0, I24.8-24.9 and I25.0-25.1, I25.6, I25.8-25.9, respectively. Myocardial infarction was ascertained using ICD-10 codes of I21.X, I22.X, I23.X, I24.1, I25.2. Coronary revascularization was assessed based on an OPCS-4 coded procedure for coronary artery bypass grafting (K40.1–40.4, K41.1–41.4, K45.1–45.5) or coronary angioplasty with or without stenting (K49.1–49.2, K49.8–49.9, K50.2, K75.1–75.4, K75.8–75.9).

Irritable bowel disease (used as a negative control phenotype) was defined according to ICD-10 codes of K85.0 or K85.9.

#### B. Pakistani Risk of Myocardial Infarction Study

The Pakistani Risk of Myocardial Infarction Study (PROMIS) was a sub-study among the entire MIGen cohort, but was selected *a priori* as a validation population. As such, the variant and sample-level QC are identical to what was described above in **Supplementary Sections 5 and 6**.

### 8. Data processing

#### A. Variant annotation

All variants underwent gene-based annotation using the ANNOVAR *geneanno* pipeline with the refGene database ^34^.Specifically, variants were classified to 1 of 9 genomic regions: 1) exonic, 2) splice donor and acceptor sites, 3) non-coding RNA (ncRNA), 4) untranslated region at the 5-prime end (UTR5), 5) untranslated region at the 3-prime end (UTR3), 6) intronic, 7) upstream, 8) downstream, or 9) intergenic. All variants were annotated to the gene harbouring the variant in question. Variants classified as exonic were further classified according to 8 mutation types: 1) frameshift insertion, 2) frameshift deletion, 3) stopgain, 4) stoploss, 5) nonframeshift insertion, 6) nonframeshift deletion, 7) nonsynonymous SNV, 8) synonymous SNV. Nonsynonymous SNVs were annotated with version 1.0 of the Mendelian Clinically Applicable Pathogenicity (M-CAP) score ^35^ for pathogenic classification. Lastly, variants were annotated according to their corresponding alternate allele frequencies in version 2.0.1 of gnomAD.

#### B. Variant pathogenicity filtering for GIAB consensus sequences and gnomAD

Due to the depletion of variants in the GIAB consensus sequences after filtering with the intersection of HCC and NIST high-confidence sites, analysis was restricted to all protein-coding SNV’s. These criteria were applied to variants present in the GIAB consensus sequences and gnomAD.

#### C. Variant frequency filtering for GIAB consensus sequences and gnomAD

Each variant within the GIAB consensus sequence was annotated with its corresponding allele frequency in a closely related population within gnomAD. Specifically, the Central European GIAB sample (NA12878) was annotated with the Non-Finnish European (NFE) population allele frequencies, the Ashkenazi Jewish GIAB sample (NA24385) was annotated with both the NFE and Ashkenazi Jewish (ASJ) population allele frequencies, and the East Asian GIAB sample (NA24631) was annotated with the East Asian (EAS) population allele frequency. Since the gnomAD supplies the alternate allele frequency (AAF) for each variant, we decided to standardize all annotated variants to the minor allele as opposed to eliminating all variants with an alternate allele frequency (AAF) > 0.5. After standardization of gnomAD allele frequency annotations, variants from the GIAB consensus sequences and gnomAD were stratified into 4 MAF threshold bins: 1) 0 ≤ MAF ≤ 0.01 (rare variants), 2) 0.01 ≤ MAF ≤ 0.05 (low-frequency variants), 3) 0.05 ≤ MAF ≤ 0.25 (common variants), and 4) 0.25 ≤ MAF ≤ 0.5 (very common variants).

#### D. Variant pathogenicity filtering for MIGen exomes, UK Biobank 50,000 exomes, and gnomAD

Percentiles were established for the M-CAP scores of all 71,561,086 nonsynonymous variants in the exome. Variants in were thereafter filtered for nonsynonymous (i.e. missense) SNVs with an M-CAP score of >0.025 (50^th^ percentile; which is also the default score for ascribing a nonsynonymous SNV as pathogenic^35^) plus disruptive variants, which included variants leading to a premature stop codon (stopgain), insertion or deletion variants altering the DNA reading frame (frameshift insertion/deletion) and SNVs within splice donor/acceptor sites (splicing).

#### E. Variant frequency filtering for MIGen exomes, UK Biobank 50,000 exomes, and gnomAD

Prior to filtering, all AAF’s annotated using gnomAD were standardized to the minor allele as stated in **Supplementary Section 8C**. To mitigate the chance of batch effects, all MIGen and UK Biobank sequence variants were also annotated with cohort-specific AAF’s standardized to the minor allele. All MIGen sequence variants with a MAF < 0.001 according to all 5 major ethnic groups in gnomAD (1) Non-Finnish European, 2) African, 3) South Asian, 4) East Asian, 5) Latin American) and cohort-specific frequencies were kept for downstream analysis. The same threshold was applied to variants in gnomAD, but these variants were not annotated with MIGen or UK Biobank cohort-specific frequencies.

### 9. Online Methods

#### A. Single gene simulations

We simulated an additive rare variant association model for a single gene with differing probabilities of mutation based on 3 confounding parameters: 1) sequencing false negative (*SFN*) rates, 2) sequencing false positive (*SFP*) rates, and 3) population-specific factors (*PSF*). *SFN* is a real number between 0 and 1 that represents the proportion of true variant calls missed by sequencing. *SFP* is a real number ≥ 0 and corresponds to the ratio of called false variants to the number of (true) variants in the reference population. The *PSF* parameter is also a real number ≥ 0 and is defined as the ratio of (true) variants in a gene relative to a reference sample. *PSF* represents the effect of population substructure, where *PSF* = 1 when the test and reference populations are perfectly matched. The probability of mutation *P* for the gene is given by equation 1:

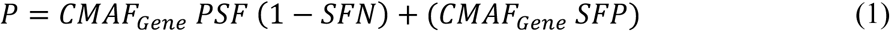

where *CMAF_Gene_* is the baseline cumulative minor allele frequency (CMAF) (defined as the aggregate sum of the MAF) of the simulated gene in the absence of any genetic effect, sequencing effects (i.e. *SFN* and *SFP*), or population effects (i.e. *PSF*). Please note use of *CMAF_Gene_* implicitly assumes variants are rare and LD negligible. When a genetic effect exists, equation 1 can be written as:

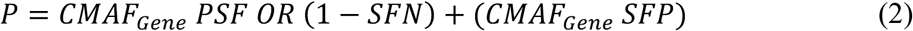

where *P* represents the probability of mutation given an individual is a case and is therefore dependent upon the *OR*, which is the true odds ratio, or genetic effect of the association. It is important to note that the *OR* represents the ratio of the frequency of rare alleles in a case population to a control population using the approximation of rare exposures:

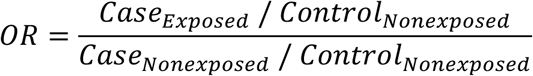

where

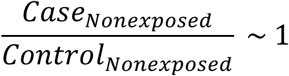

for rare exposures (i.e. the *CMAF* attributed to the aggregate rare allele frequency within a gene). The equation for *OR* thus simplifies to

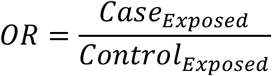

which we also empirically confirmed.

We then derived a correction factor (*CF*) to adjust for sequencing false positives, negatives, and population substructure effects. *CF* is defined as following, which is a direct simplification of equation 2 under the assumption of no genetic effect (*OR*=1):

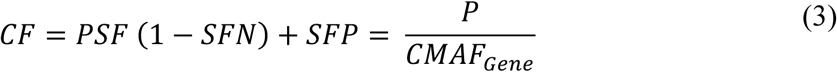

*CF* can be estimated by calculating the ratio of observed variants to expected variants in the reference population (i.e. *CMAF_Gene_*) over all protein-coding genes genome-wide (or at least a large number of genes). Since *CF* is empirically estimated across all protein-coding genes, it is assumed that the effect of truly associated genes on genome-wide estimates is negligible. Estimated 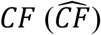 can then be used to calculate an adjusted *P\** for the gene:

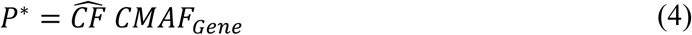

We then sought to determine whether use of *P*^∗^ (as opposed to *P* = *CMAF_Gene_*) would improve association testing in the presence of confounding (i.e. *SFN* > 0, *SFP* > 0 or *PSF* <>1). To test whether use of *P\** provides unbiased estimates of OR, we varied the true (unobserved) *OR* from 1 (null effect) to 2 (strong effect) at intervals of 0.1. Each 34 was simulated 100,000 times in 1000 individuals and the mean (SD) estimated 34 calculated. In all simulations, SPN, SPF and PSF were set at 0.3 to illustrate conditions of strong confounding. To ascertain type I and II errors we performed a second set of simulations, fixing the true (unobserved) 34 to 1.3 and varying SFN and *SFP* from 0 to 1 and *PSF* from 0 to 2. Each condition was simulated 100,000 times in 1,000 participants. *P*-values were calculated using the binomial exact test assuming either *P\** or *P*.

#### B. Sequence coverage harmonization

Differential sequence coverage driven by non-biological factors between any two cohorts can render genetic association signals prone to false positives or negatives. As such, restricting analyses to sites that are sensitive to variant detection in both gnomAD and comparator samples (i.e. GIAB and MIGen) will mitigate detection of spurious associations. Per variant coverage statistics for gnomAD release 2.0.1 were downloaded on the command line using the Google Cloud Storage gsutil tool. First, variant sites were intersected against an interval file corresponding to refGene coding regions across the genome as defined by the Reference Sequence (RefSeq) database (the interval file was retrieved from the University of Southern California (UCSC) Table Browser: https://genome.ucsc.edu/cgi-bin/hgTables). Second, the average proportion of individuals with ≥ 20X coverage was determined for each coding interval. Intervals in which ≥ 90% of individuals had ≥ 20X coverage were deemed as “high-coverage coding (HCC)” intervals. Lastly, the resulting HCC intervals were intersected against high-confidence regions (using the NIST high-confidence regions for GIAB samples and cohort-specific exome enrichment intervals for MIGen samples; see Supplementary Table 1). The resultant final intersected regions would therefore be characterized by HCC regions that are sensitive to detecting variants in gnomAD whilst still capturing high quality and enriched regions in GIAB and MIGen samples, respectively.

#### C. Benchmarking correction factors for GIAB consensus sequences

Correction factors for each individual were calculated based on the ratio of mutation loads between a GIAB comparator sample and the gnomAD dataset. Specifically, variant counts across 18410 autosomal protein-coding genes curated in the refGene coding regions file (see **Supplementary Section 8B**) were aggregated to generate observed mutation load in the consensus GIAB sample and then compared to the aggregate expected mutation load obtained from gnomAD frequencies across the same set of genes (equation 5)

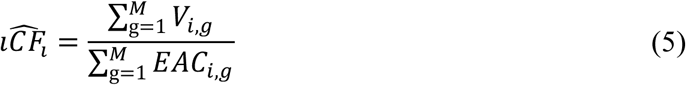

where 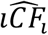 represents the estimated individual correction factor for individual *i* which is equal to the ratio between the sum of variant counts *V* across all * *M* genes in the GIAB comparator sequence to the sum of the estimated allele count in gnomAD (Q+)) (i.e. 2)*+$) for the same * genes. For simplicity, 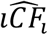 is thereafter referred as *iCF*). Mutation loads for both gnomAD and GIAB were restricted to the intersection of variant sites between gnomAD HCC and the NIST high-confidence regions for a given GIAB sample. Q+) were generated using ethnic-specific allele frequencies that best matched the ethnicity of the comparator sample in order to limit population stratification bias. Specifically, the Q+) for comparison with the Central European GIAB sample (NA12878) was generated using the Non-Finnish European AAF’s in gnomAD (see Supplementary Table 3 for all GIAB to gnomAD ethnicity matches). A 95% confidence interval (CI) was calculated for the *iCF* in each allele frequency bin using cumulative Bernoulli variance where the probability of success was denoted by the gene *CMAF*.

**Supplementary Table 3:** Ethnicities in gnomAD used to generate the Q+) for a given GIAB sample.

| ID of GIAB sample | Ethnicity of GIAB sample | Ethnicity in gnomAD used to determine the *EAC* |
| --- | --- | --- |
| NA12878 | Central European | Non-Finnish European |
| NA24631 | Ease Asian | East Asian |
| NA24385 | Ashkenazi Jewish | Non-Finnish European |
|  |  | Ashkenazi Jewish |

#### D. Determining R)iCF heterogeneity for rare variants in GIAB consensus sequences

Since a single value of R) *iCF* is to be used to equilibrate rare mutation burden across the exome, it is imperative to identify whether *iCF* differs significantly across genes stratified by different levels of constraint (i.e. a gene’s tolerance towards harbouring deleterious mutations). To test this, constraint scores calculated as previously described^36^ for all genes were downloaded from the Exome Aggregation Consortium (ExAC) release 0.3.1 ftp repository (ftp://ftp.broadinstitute.org/pub/ExAC_release/release0.3.1/functional_gene_constraint/fordist_cleaned_exac_r03/march16_z_pli_rec_null_data.txt). Genes were stratified into deciles based on their z-score for missense variant constraint and heterogeneity of the mean and standard deviation (determined from the 95% CI described in **Supplementary Section 8C**) of the *iCF* for each decile tested using *rMeta* R package^37^.

#### E. Individual correction factor (iCF) in MIGen

The individual Correction Factor *(iCF)* was calculated for each MIGen participant as per **equation 5**. Variant sites contributing to mutation load were selected from the intersection of gnomAD HCC regions and the exome enrichment intervals for each MIGen cohort (Supplementary Table 1). Upon generating an *iCF* for every MIGen participant, an *iCF-*adjusted *P* (i.e. *P*\*) was calculated per sample per gene according to:

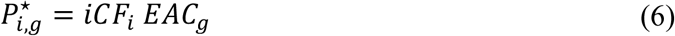

As for the GIAB analysis, the Q+) for each gene was based on the gnomAD population that was most closely related to the ethnicity of the MIGen cohort (Supplementary Table 1). For example, the Q+) for all European cohorts was generated using the NFE AAF’s whereas the Q+) for the South Asian cohort used the South Asian AAF’s in gnomAD.

#### F. Gene Correction Factor (gCF) in MIGen discovery exomes

It was observed that some genes are consistently enriched or depleted in rare mutations in healthy controls from multiple cohorts as compared to gnomAD, pointing to gene-specific sequencing false positives/negatives or gene-specific population structure effects. Hence, a gene correction factor (*gCF*) was instituted to adjust for biases in gene mutation burden not captured by (*gCF*). To determine the (*gCF*), genes were binned according to their association signals in an independent calibration cohort that was 1) sequenced using the same exome enrichment chemistry and technological platform as the test cohort and 2) absent of samples with phenotypes that were either identical to, or risk factors for the phenotype characterizing the test cohort (e.g. a cohort with samples positive for MI or type 2 diabetes mellitus could not be used to calibrate a test cohort of MI cases). The rationale for these criteria are to bin genes with similar biases in mutation burden (criteria 1), but not based on mutation burden due to presence of disease-conferring mutations (criteria 2). To fulfill these two criteria, gene bins were established using first, quintiles of expected counts (in ascending order) and second, deciles of *P*-values (described in **Supplementary Section 8G**) (from most to least significant) within a ranking cohort consisting of 6,082 disease-free controls across 8 MIGen populations (Supplementary Table 1 and Supplementary Table 2). As *P*-values are one-sided, the ranking will order genes from the one with the greatest evidence for enrichment in rare pathogenic mutations in MIGen healthy controls versus gnomAD to the one with the greatest evidence for depletion in rare, pathogenic alleles in MIGen healthy controls versus gnomAD. Additionally, by ordering by *P*-value within quintiles of expected count, we account for genes that may exhibit significant association signals by virtue of limited power to detect mutations. A *gCF* was then calculated according to equation 9:

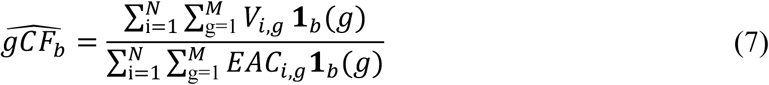

where 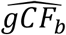 is the estimated gCF value across *N* individuals and *M* genes in bin *b* and 1*_b_(X)* is 1*_b_(X)* is the indicator function with value 1 if gene *g* is part of bin *b* and 0 otherwise. 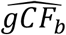 is thereafter referred as gCF for simplicity. A bin size of ∼232 (50 bins across 11,616 genes with a *V_g_* and a *EAC_g_* of at least 1 assessed in the ranking cohort) ensures that the proportion of disease-causing genes is expected to be small in each bin. Selecting smaller bin sizes (i.e ≤ 50) ultimately nullifies the per gene association signals, which precludes inclusion of a gCF. In other words, small bin sizes will tend to bias results towards the null of no association. In contrast, extensively large bin sizes will bin genes together that do not necessarily share a similar bias in mutation load, which will dilute the effect of the *gCF* and render it ineffective. An *iCF* and *gCF* adjusted *P* (i.e. *P*\*\*) was then calculated for each gene and individual according to:

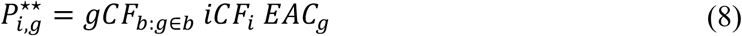

#### G. Rare variant association

To test associations of individual genes with MI, a burden test was used to evaluate the difference in aggregate counts for rare, pathogenic alleles (as defined in **Supplementary Section 8D**) between MIGen cases (N=5,910) and the *iCF* and *gCF*-adjusted *EAC*). To account for potential LD between rare variants as well as biases in mutation burden between test samples and gnomAD, we introduce a novel procedure based on individual differences between observed and expected mutation counts. Specifically, for each gene and individual, we define the delta count (*D*) as:

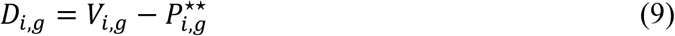

For each gene, we then test whether mean *D_i,g_*(i.e. *D_g_*) is equal or different than zero using a Z test since *D_g_* is expected to be equal to zero under the null hypothesis of no association:

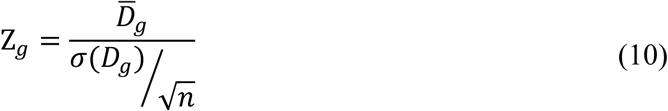

This procedure has several advantages. First, it adjusts for biases in mutation burden between the test samples and gnomAD. Second, it leverages the fact LD between variants will increase variance (i.e. σ^2^(*D_g_*)) but not expected *D_g_*. As σ(*D_g_*) is empirically derived, *a priori* knowledge of the LD structure is not necessary.

#### H. Evaluating enrichment in genes found to be genome-wide significant loci in CARDIoGRAMplusC4D

Genes from the discovery rare variant association in MIGen were cross referenced against loci harboring a genome-wide significant (GWS) variant (*P*<5×10^-8^) in the CARDIoGRAMplusC4D consortium^38^. A total of 36 CAD genes were identified among the 7,146 discovery genes, including the low-density lipoprotein receptor (*LDLR*). Due to the disproportionate association signal for *LDLR*, a combined test statistic was generated from the remaining 35 CAD genes using Fisher’s combined probability test. To test the null hypothesis that the combined test statistic for the CAD gene set was not significantly different from that of random gene sets, we generated a null distribution of combined test statistics for 100,000 randomly selected gene sets of identical size (also excluding *LDLR*) and identified the probability of observing a test statistic greater than or equivalent to that of the CAD gene set.

#### I. Rare variant association using TRAPD

The distribution of gene-base test statistics generated using RV-EXCALIBER were compared to those generated using TRAPD. Specifically, healthy control subjects from the ATVB, OHS, BioImage, and Munich-MI cohorts (N=3,352) were used as the “case” dataset and tested against non-Finnish Europeans in gnomAD, which was used as the control population. For TRAPD, quality by depth (QD) values were generated in cases by computing the ratio of the variant-level quality to the sum of the depth of coverage for all non-homozygous reference genotypes across a single variant. QD values in gnomAD were extracted directly from the INFO field present in the downloaded VCF (**Supplementary Section 1**). To remain consistent, variant frequency and pathogenicity filtering along with sequencing coverage harmonization was completed exactly as performed for RV-EXCALIBER (**Supplementary Sections 8D, 8E, and 9B**). The distribution of QD values was determined for all rare synonymous SNVs in cases to obtain variant sets corresponding to the top 50^th^ to 99^th^ percentiles of QD scores. For gnomAD we kept only the rare synonymous SNVs corresponding to the top 70^th^, 75^th^, 80^th^, 85^th^, 90^th^, and 95^th^ percentile of QD scores. Per gene *P*-values were generated for each QD percentile set from 2×2 contingency tables using a Fishers exact test followed by calculation of lambda-95 (f_gh_) estimate as previously described^39^. The percentile set with the best calibrated f_gh_ was used to inform inclusion of nonsynonymous SNVs for burden testing. Inclusion of INDELs were informed using the same procedure, but were calibrated using rare nonsynonymous SNVs instead of synonymous SNVs. For RV-EXCALIBER, gene-based test statistics were calculated as previously described where healthy controls from the PROCARDIS, U. Lubeck, REGICOR, and MDC cohorts (N=2,730) were used as the ranking set for calculation of gCF.

#### J. Rare variant genetic risk score (RVGRS)

Gene-based odds ratios and corresponding regression coefficients were generated from rare variant association analysis between the discovery MIGen samples (Supplementary Table 1) and gnomAD according to:

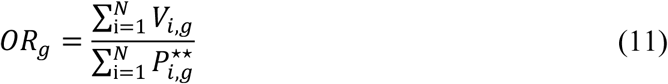

and

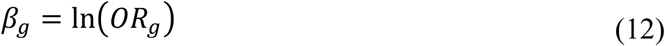

weighted rare variants genetic risk scores (RVGRS) were then derived for each individual by summing the product of D_i,g_ and β_g_ across all genes (included in the gene score):

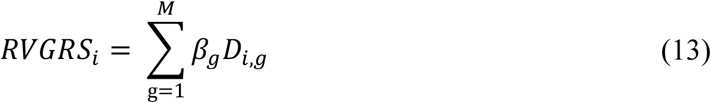

The RV-EXCALIBER model was used to evaluate the association of RVGSR_i_ with case status in validation cases independent from the derivation sample. For the UK Biobank, we selected the entire population (N=45,850) as the ranking cohort (due to it being an unselected prospective cohort study) in order to calculate the DIG term to be weighted by the discovery effect size estimates. For PROMIS, we used the same ranking cohort as was used for the discovery association test **(Supplementary Section 9F**).

Inclusion of genes from the discovery gene associations to generate the *RVGSR_i_* the validation cohorts was done in a cumulative stepwise manner. Specifically, the *RVGSR_i_* were built using the top 10 to 3000 most significant genes by including 10 additional genes per step, for a total of 300 sets of *RVGSR_i_* in the validation cohort. Logistic regression was performed to evaluate the association between *RVGSR_i_* and prevalent CAD in the UK Biobank exomes or MI in PROMIS while adjusting for age, age^2^, sex and the first 20 genetic principal components. The distribution of raw per-individual RVGRS scores were scaled to a mean of 0 and a SD of 1 prior to association testing. The OR for an increase of 1 SD in the score was obtained by calculating the natural exponential function of the regression coefficient. Significance for all trend calculations was conducted by regressing case affection status with an interaction term between the scaled RVGRS and the parameter of interest. We defined substantial risk conferred as having a RVGRS corresponding to a ≥2-fold increased odds of disease.

#### K. Calculating the Framingham Risk Score

A 10-year estimation of cardiovascular disease risk was determined using a Framingham Risk Score (FRS)^40^. The FRS incorporated age, sex, high-density lipoprotein cholesterol, total cholesterol, systolic blood pressure, antihypertensive medication, and smoking status to generate a composite score on a per individual basis. Missing clinical data was classified as “missing completely at random” and was imputed using multivariate imputation by chained equations^41^. We opted to use recommended values of 5 imputed datasets over 50 iterations using predictive mean matching. To test the hypothesis that individuals with low burden of clinical risk factors had higher genetic risk of disease, we regressed case affection status with a multiplicative interaction term consisting of standardized RVGRS and tertile of FRS while still using age, age^2^, sex, and the first 20 principle components of ancestry as covariates.

#### L. Ascertaining a common variant gene risk score (CVGRS) for UK Biobank participants

Initial common variant gene risk scores (CVGRS) were constructed as previously described^42^. LD adjustment was incorporated as published in the GraBLD study, where weights were corrected for each SNP such that all SNPs can be included in the gene score^43^. The default value of 300 SNPs upstream and downstream from the target SNP is considered.

PLINK’s --score function^30^ was the main method of generating the CVGRS, which accordingly applies a linear scoring system to the inputted genotype matrix. SNPs were matched between UKB and GWAS data from the CARDIoGRAMplusC4D consortium with 1000 genomes imputation^38^, after which all duplicate and triallelic SNPs were removed. All scores were corrected for threshold and corrected for LD. Initially, various *P*-value thresholds and LD windows values were tested to seek for an optimal value. A default value of 0.1 for threshold and LD window size of 300 was utilized for all initial scores. All scores are then standardized to a mean of 0 and SD of 1. After LD pruning, a total of 838,897 SNPs were used to construct the CVGRS.

Interaction between standardized CVGRS and FRS tertile was conducted as described in **Supplementary Section 9L.**

#### M. Evaluating net reclassification improvement index using RVGRS

We evaluated the net reclassification improvement (NRI) index when incorporating RVGRS into a base model consisting of CVGRS and FRS. Specifically, we first generated a logistic regression model where case affection status was regressed against CVGRS, FRS, age, age^2^, sex, and the first 20 principle components. A second model was then generated by regressing affection status against RVGRS and the aforementioned independent variables. Both models were thereafter imported into the *Hmisc* R package^44^ to ascertain the NRI index.

#### N. Stratifying risk conferred by RVGRS based on age of CAD onset

Age of onset for each ICD-10 phenotype was defined according to date of first in-patient diagnosis (UK Biobank data field 41280). Age of CAD onset was ascertained according to the first occurring event in our composite CAD definition (**Supplementary Section 7A**). Age of onset was stratified into 4 categories: i) males ≤40 and females ≤45, ii) males ≤40 and females ≤45, iii) males ≤45 and females ≤50, and iv) males ≤55 and females ≤60. Enrichment of RVGRS in cases from the 4 age strata were evaluated against the UK Biobank control population. Association with CAD was evaluated using sex and the first 20 principle components as covariates. Age and age^2^ were not used as covariates since the age distribution between early-onset cases and controls is necessarily significant.

## SUPPLEMENTAL RESULTS

**Supplementary Table 4:** iCF values for every GIAB vs. gnomAD ethnicity across 4 allele frequency bins.

| AF bin | GIAB ethnicity | gnomAD population | iCF | 95% CI |
| --- | --- | --- | --- | --- |
| 0-0.01 | Central European | Non-Finnish European | 0.60 | 0.54-0.68 |
|  | East Asian | East Asian | 0.93 | 0.84-1.04 |
|  | Ashkenazi | Ashkenazi | 0.93 | 0.84-1.04 |
|  | Ashkenazi | Non-Finnish European | 1.37 | 1.22-1.55 |
| 0.01-0.05 | Central European | Non-Finnish European | 0.59 | 0.55-0.65 |
|  | East Asian | East Asian | 1.00 | 0.92-1.11 |
|  | Ashkenazi | Ashkenazi | 0.93 | 0.86-1.01 |
|  | Ashkenazi | Non-Finnish European | 0.98 | 0.91-1.08 |
| 0.05-0.25 | Central European | Non-Finnish European | 0.80 | 0.78-0.83 |
|  | East Asian | East Asian | 0.98 | 0.95-1.01 |
|  | Ashkenazi | Ashkenazi | 0.99 | 0.96-1.02 |
|  | Ashkenazi | Non-Finnish European | 0.98 | 0.95-1.01 |
| 0.25-0.50 | Central European | Non-Finnish European | 0.93 | 0.92-0.95 |
|  | East Asian | East Asian | 0.97 | 0.96-0.99 |
|  | Ashkenazi | Ashkenazi | 1.00 | 0.99-1.02 |
|  | Ashkenazi | Non-Finnish European | 1.02 | 1.01-1.04 |

**Supplementary Table 5:** Median iCF values for healthy control participants within 8 MIGen cohorts.

| MIGen cohort | Median iCF (IQR) |
| --- | --- |
| ATVB | 1.135 (0.909-1.363) |
| PROCARDIS | 0.841 (0.636-1.045) |
| OHS | 0.817 (0.613-1.022) |
| BioImage | 0.797 (0.576-1.019) |
| U. Lubeck | 0.795 (0.591-1.000) |
| Munich-MI | 0.750 (0.545-0.954) |
| REGICOR | 0.731 (0.525-0.936) |
| MDC | 0.664 (0.487-0.841) |

IQR=inter-quartile range

**Supplementary Table 6:** Effect size estimates of iCF distribution in MIGen control versus case participants.

| **MIGen cohort** | **OR (95% CI)\*** | **P-value** |
| --- | --- | --- |
| ATVB | 0.83 (0.56-1.25) | 0.381 |
| PROCARDIS | 0.75 (0.40-1.40) | 0.370 |
| OHS | 0.73 (0.37-1.44) | 0.364 |
| BioImage | 0.86 (0.17-4.49) | 0.857 |
| U. Lubeck | 0.96 (0.48-1.92) | 0.911 |
| Munich-MI | 0.39 (0.14-1.10) | 0.0745 |
| REGICOR<sup>‡</sup> | 0.33 (0.12-0.92) | 0.0352 |
| MDC<sup>‡</sup> | 2.44 (1.01-5.96) | 0.0492 |

\*OR is expressed as controls relative to cases
<sup>‡</sup> Nominal significance (P<0.05)

**Supplementary Table 7:**
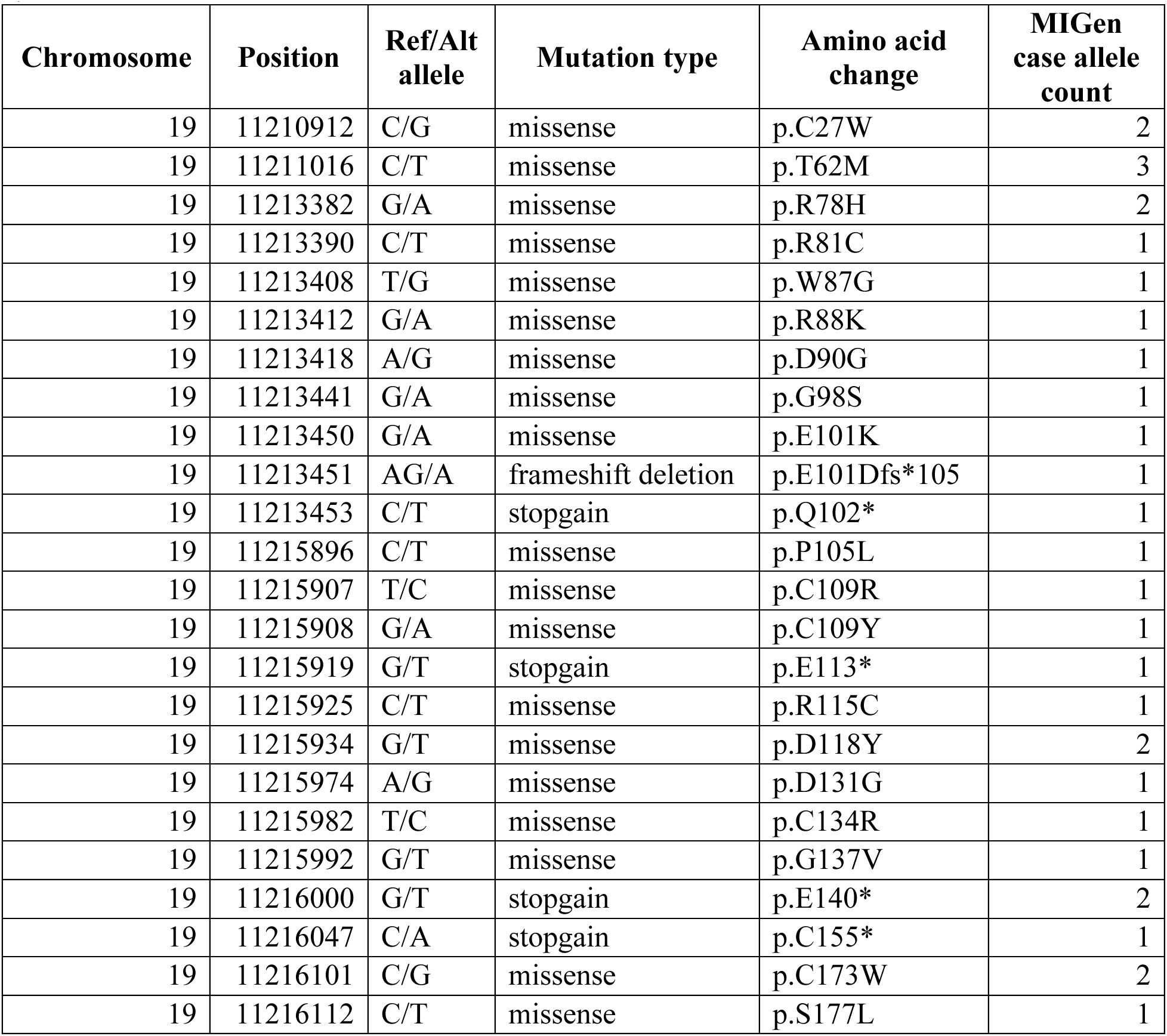
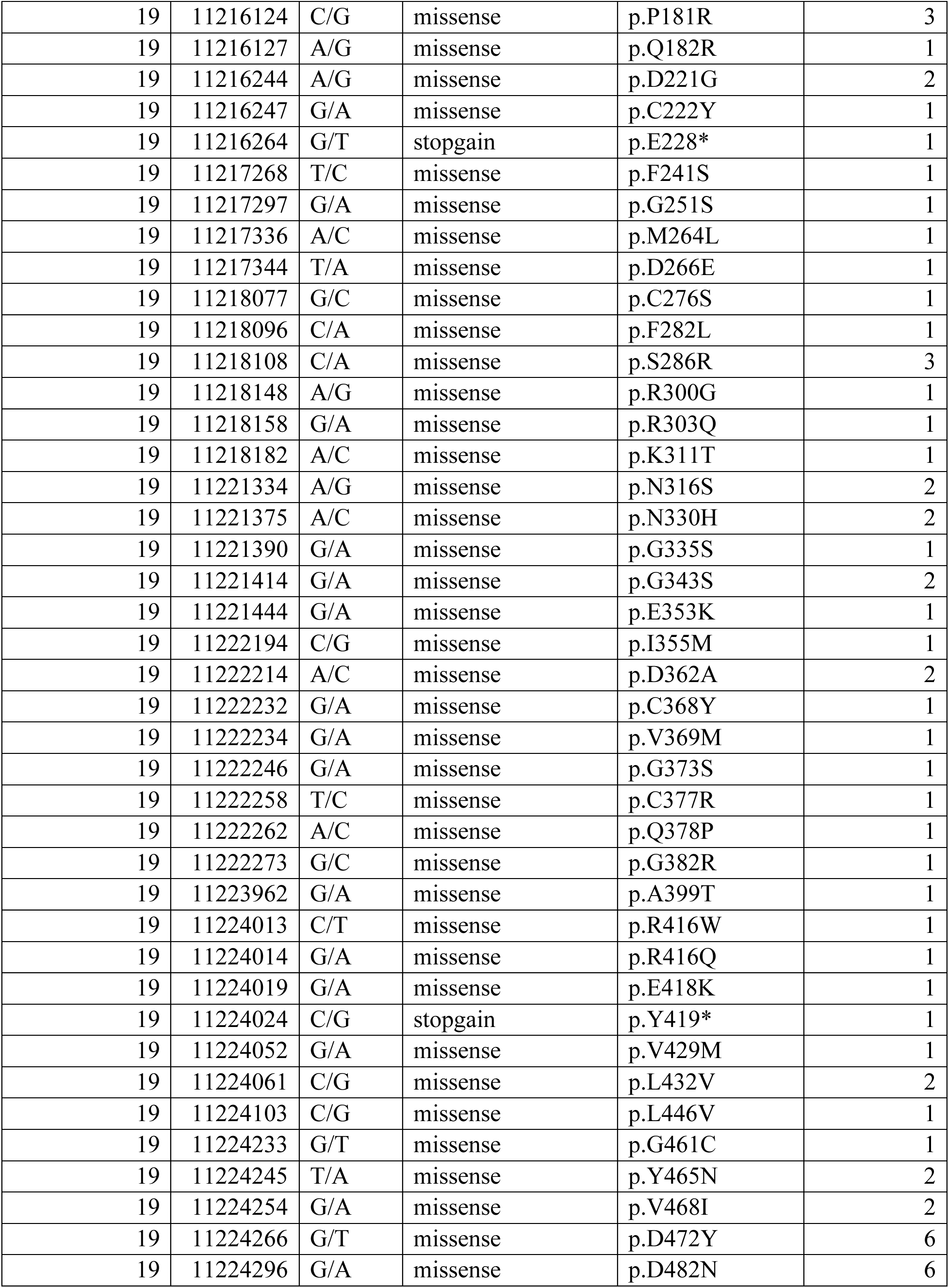
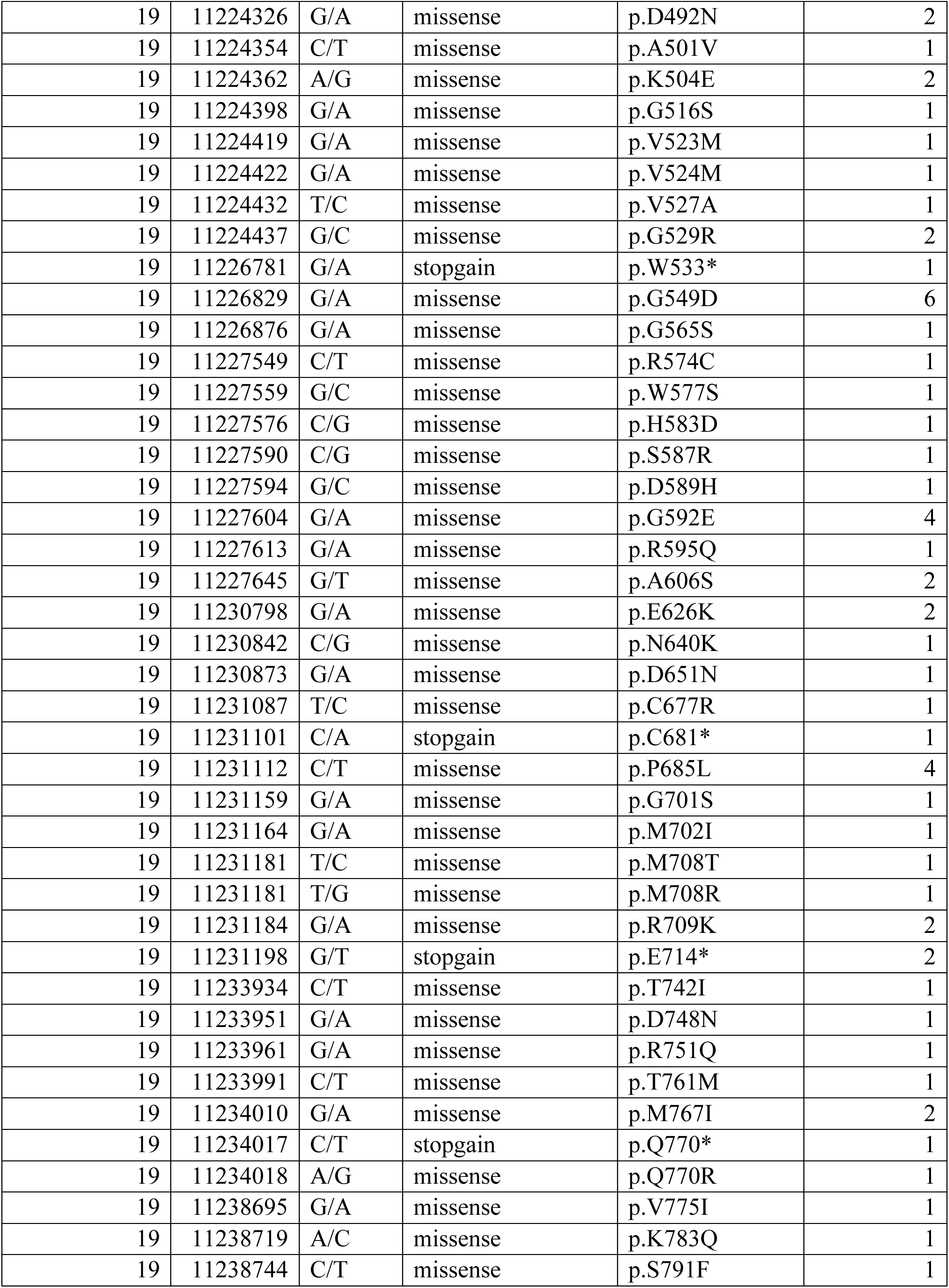
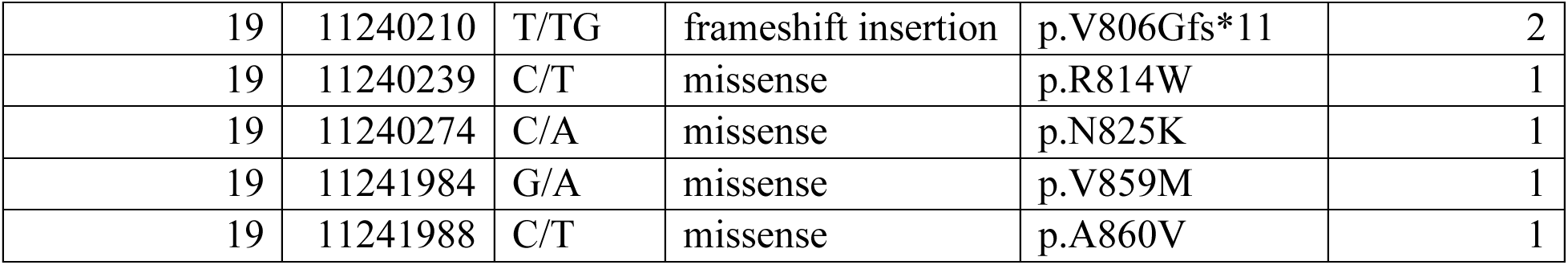
Characteristics of rare pathogenic alleles observed in *LDLR* among 5,910 cases from the MIGen consortium.

**Supplementary Table 8:**
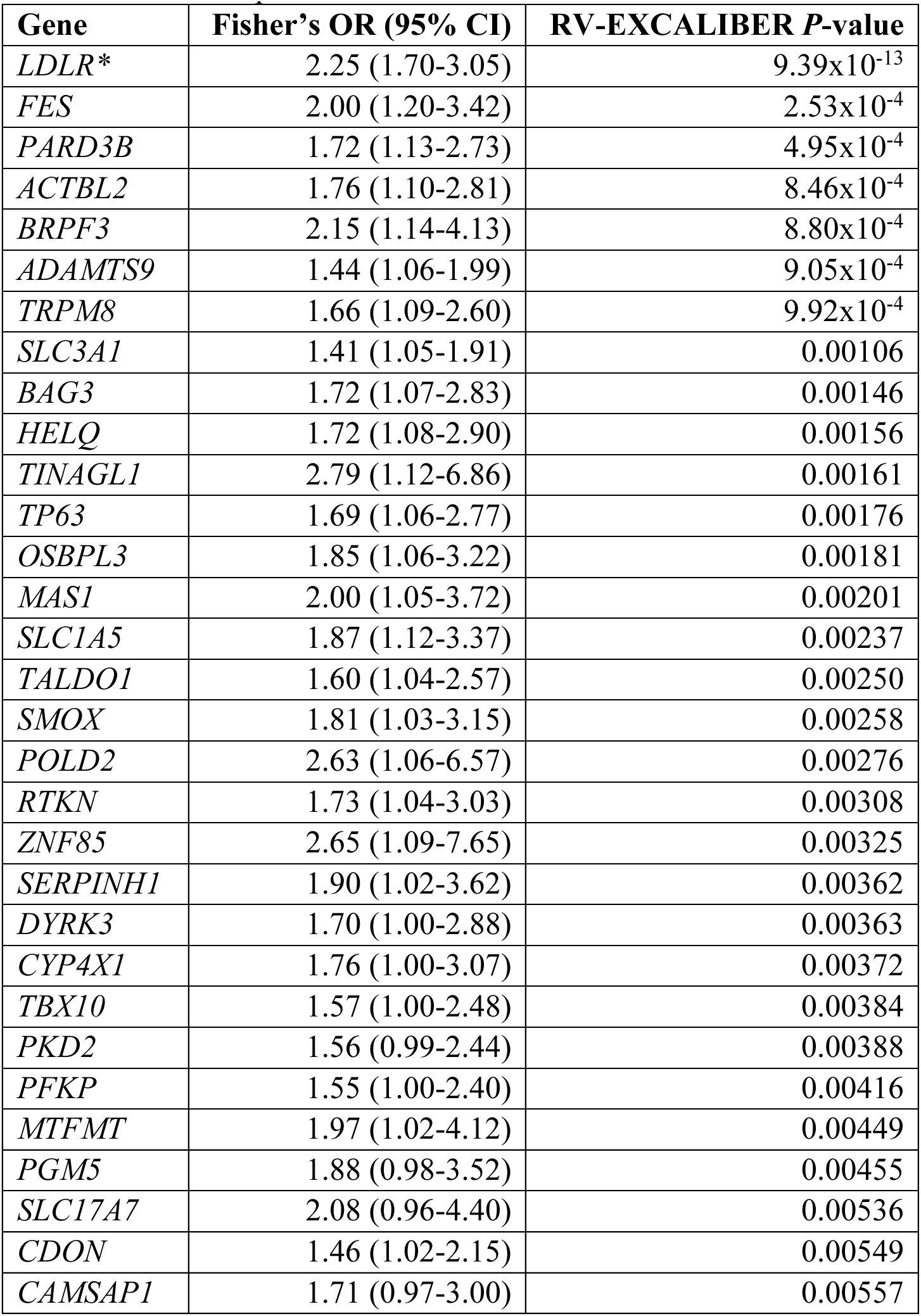
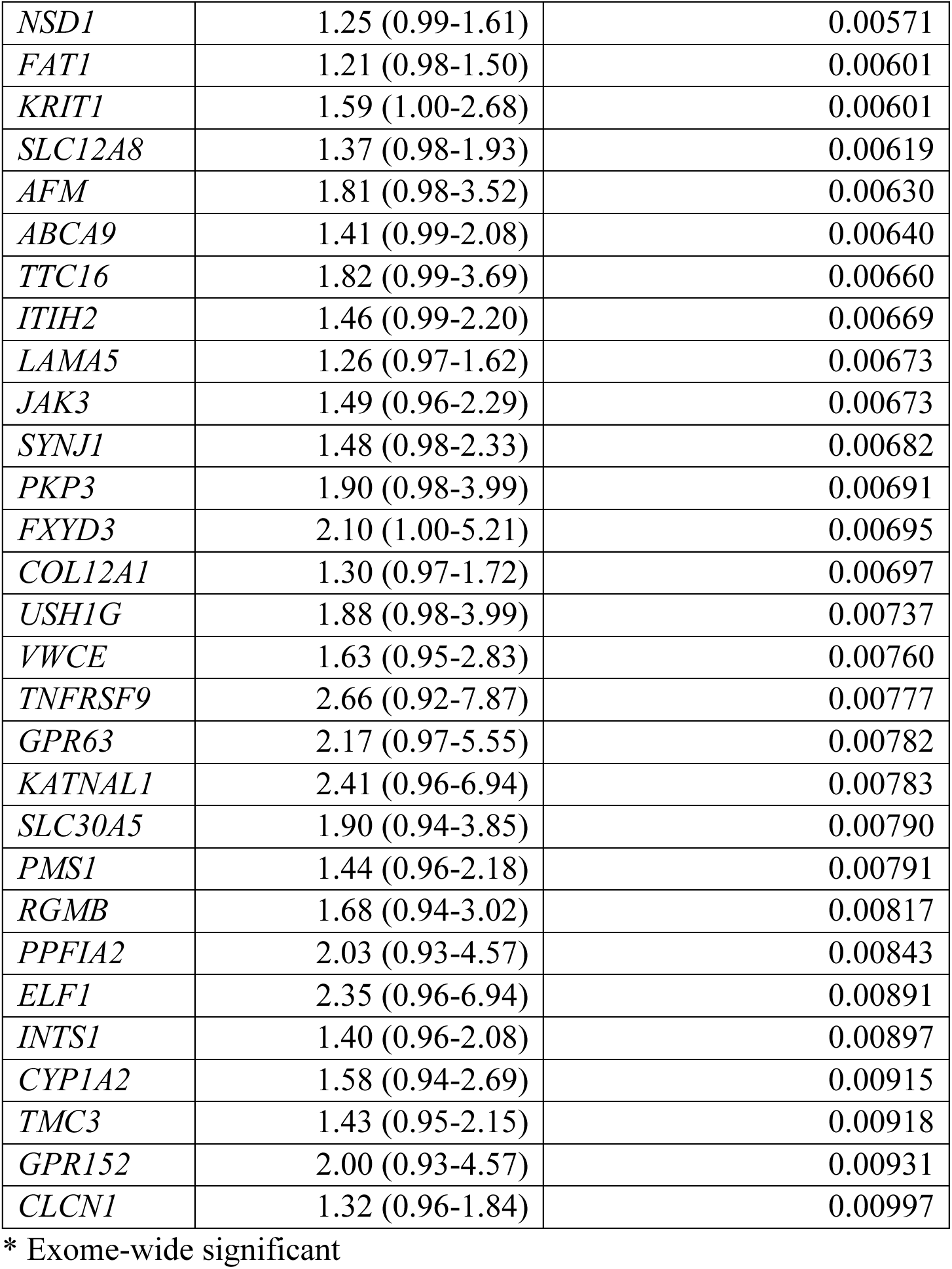
Top gene-based associations using expected counts that were adjusted with RV-EXCALIBER. Variants included were nonsynonymous SNVs with an M-CAP score >0.025 and all disruptive variants.

**Supplementary Table 9:**
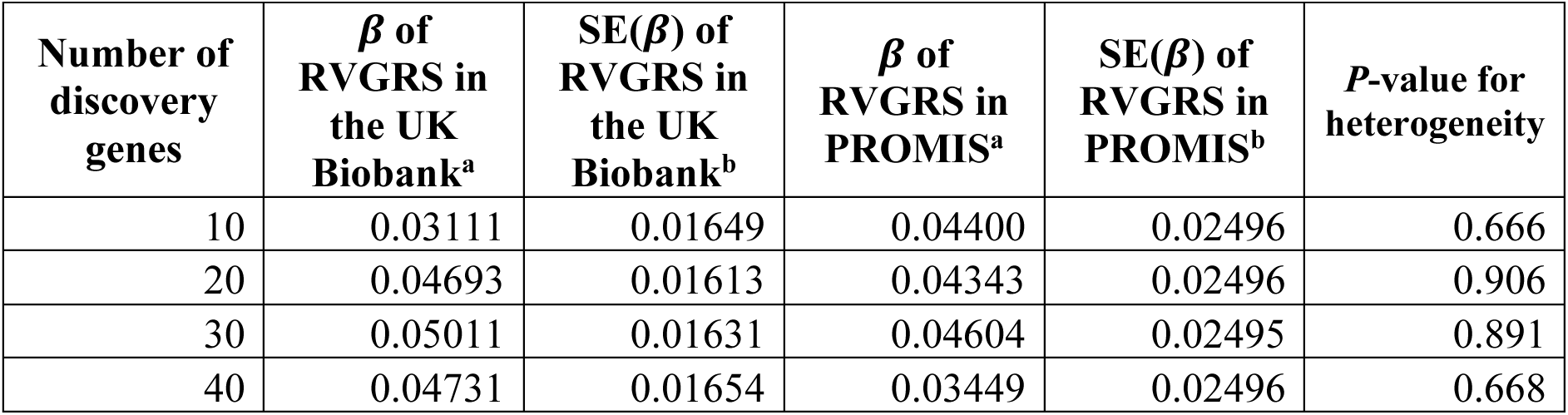
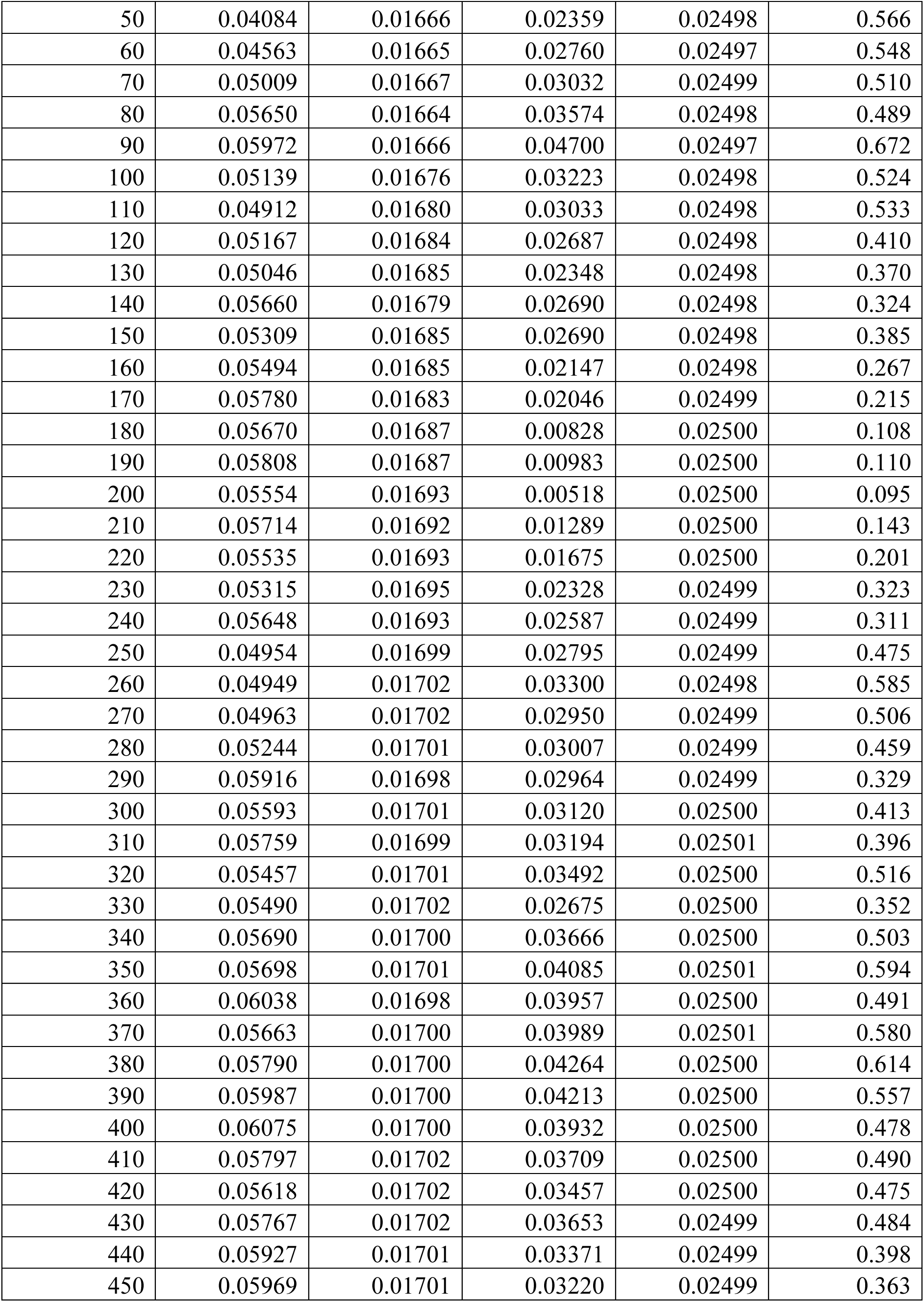
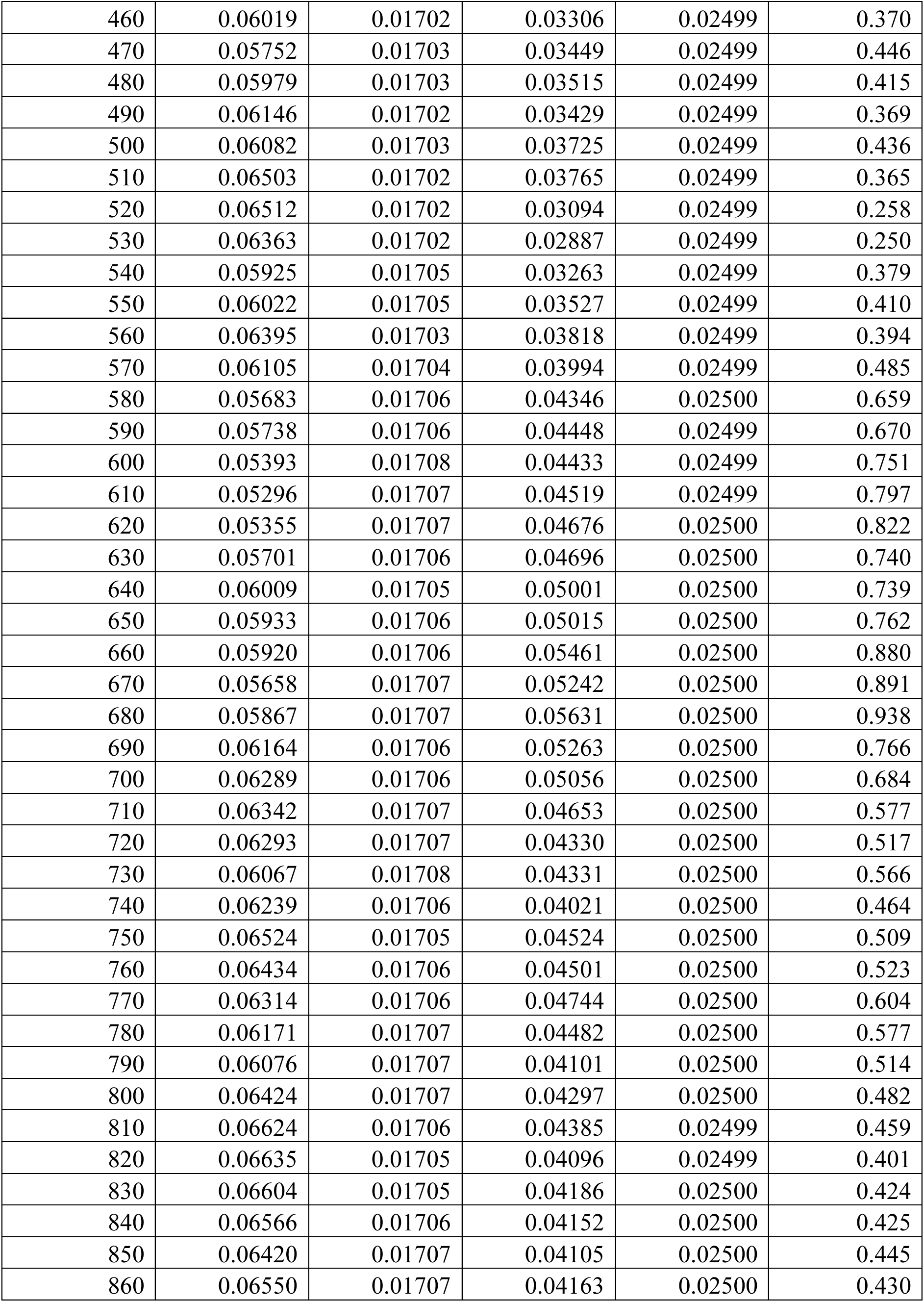
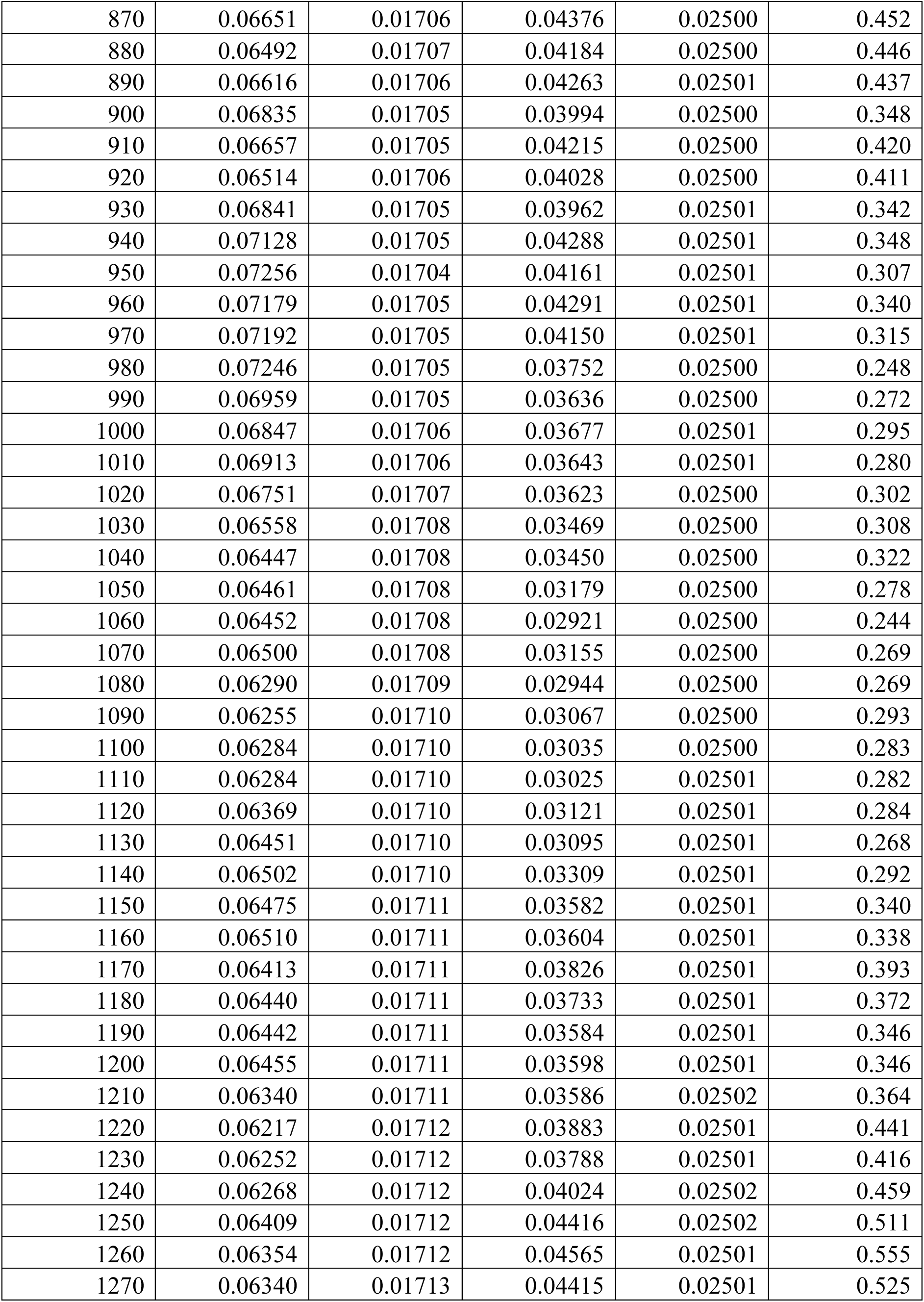
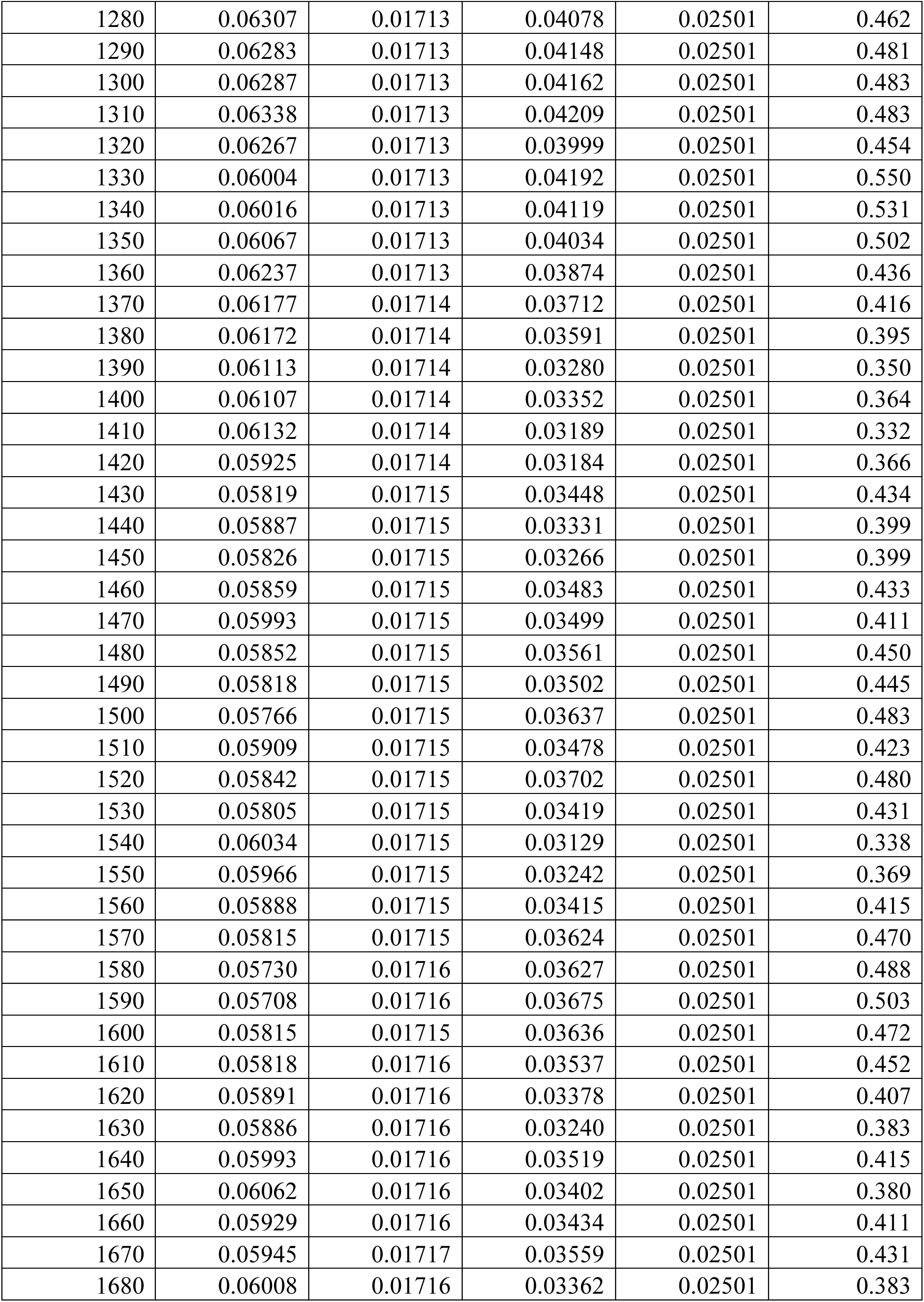
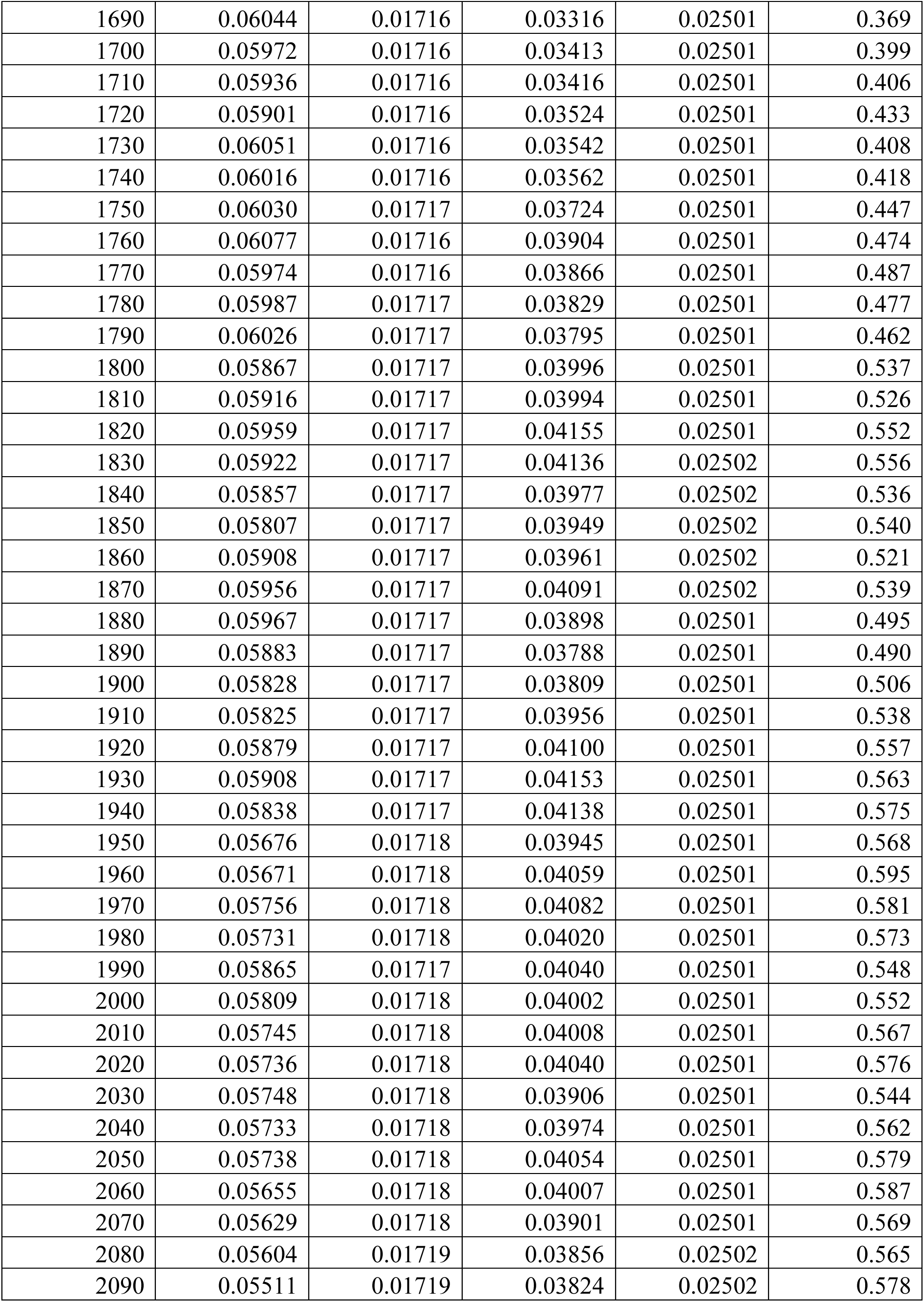
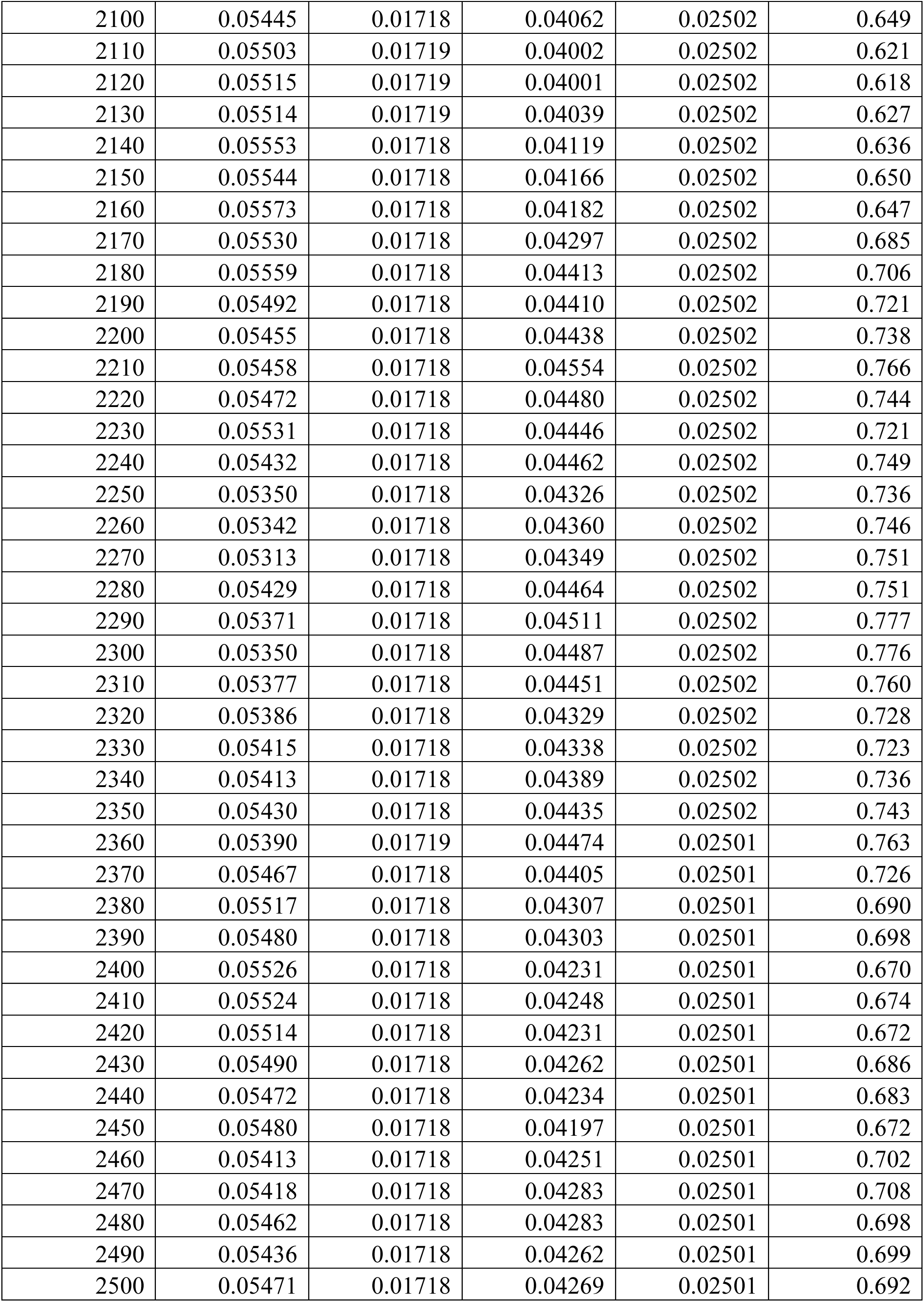
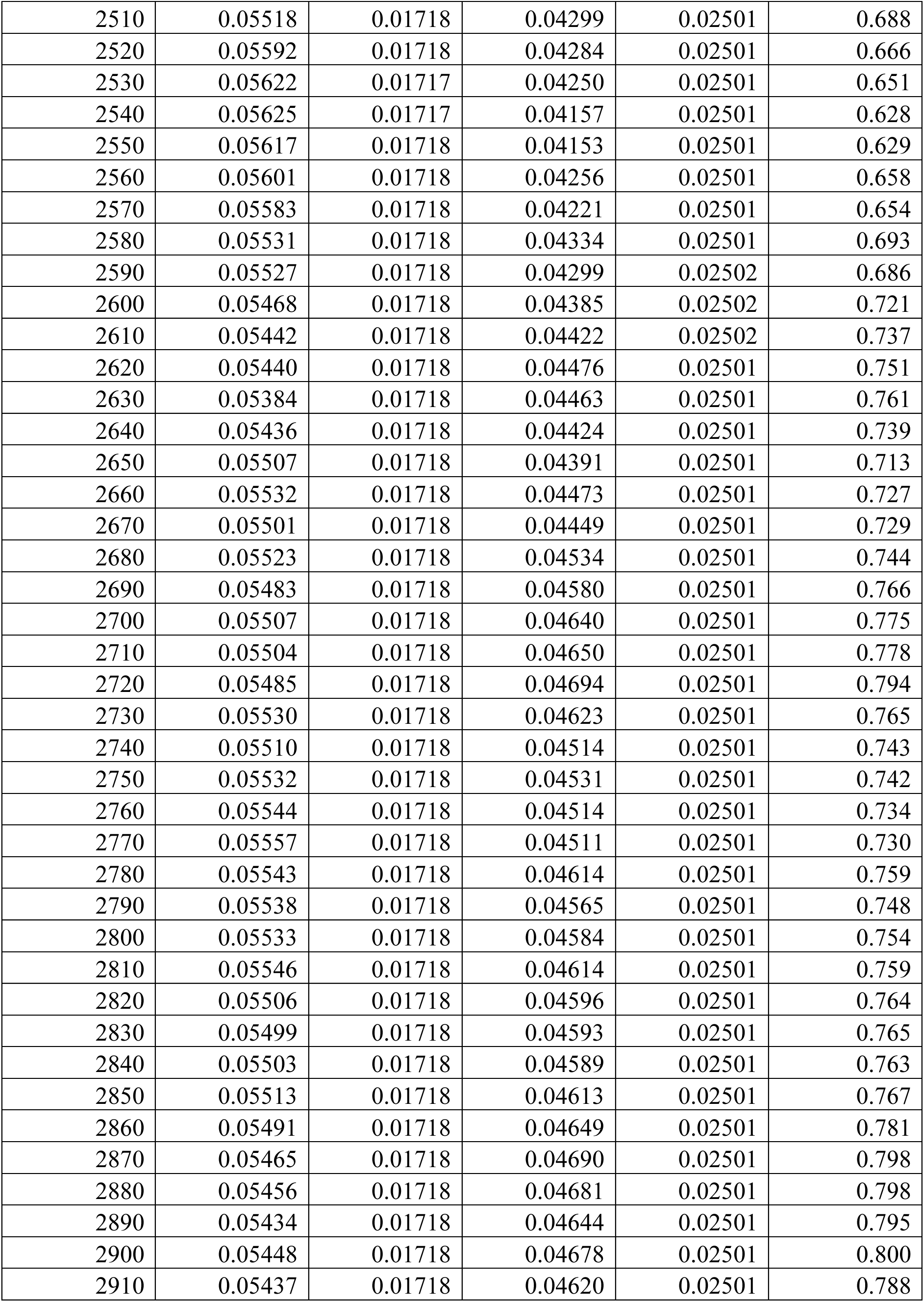
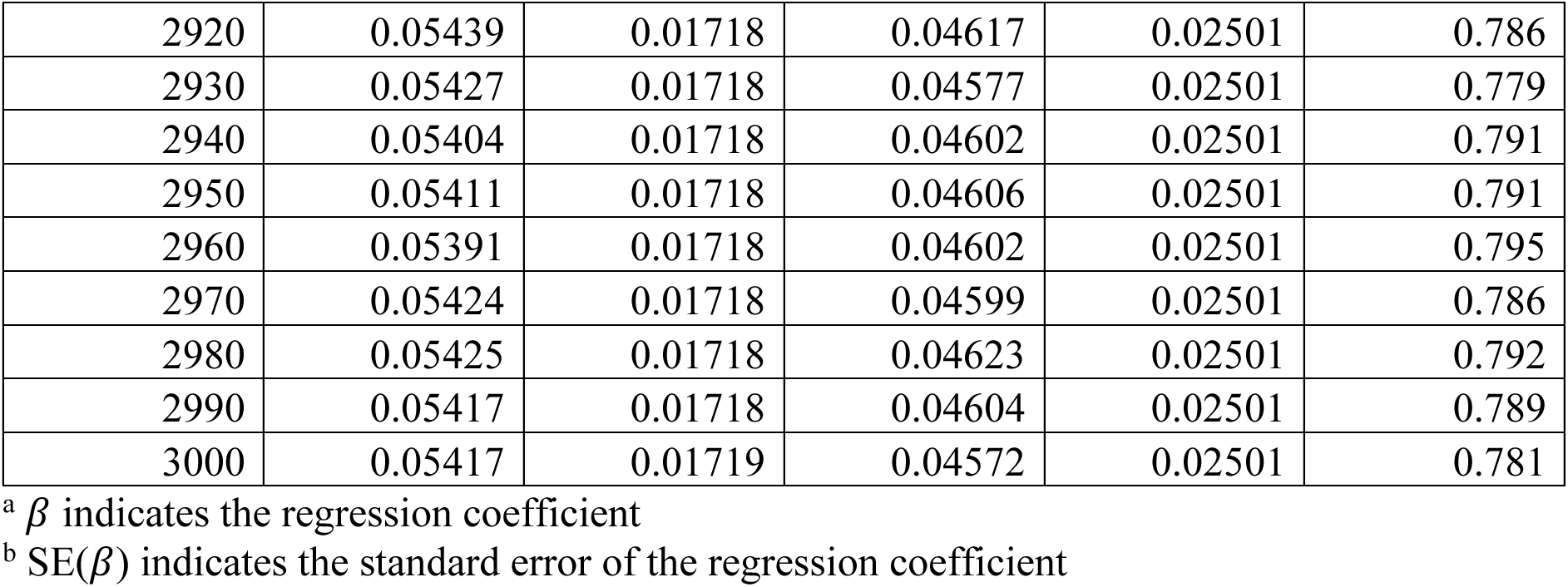
Effect estimates and heterogeneity *P*-values for 3,000 RVGRS <u>determined in the UK Biobank and PROMIS exomes.</u>

| Number of discovery genes | $\beta$ of RVGRS in the UK Biobank<sup>a</sup> | SE($\beta$) of RVGRS in the UK Biobank<sup>b</sup> | $\beta$ of RVGRS in PROMIS<sup>a</sup> | SE($\beta$) of RVGRS in PROMIS<sup>b</sup> | *P*-value for heterogeneity |
| --- | --- | --- | --- | --- | --- |
| 10 | 0.03111 | 0.01649 | 0.04400 | 0.02496 | 0.666 |
| 20 | 0.04693 | 0.01613 | 0.04343 | 0.02496 | 0.906 |
| 30 | 0.05011 | 0.01631 | 0.04604 | 0.02495 | 0.891 |
| 40 | 0.04731 | 0.01654 | 0.03449 | 0.02496 | 0.668 |
| 50 | 0.04084 | 0.01666 | 0.02359 | 0.02498 | 0.566 |
| 60 | 0.04563 | 0.01665 | 0.02760 | 0.02497 | 0.548 |
| 70 | 0.05009 | 0.01667 | 0.03032 | 0.02499 | 0.510 |
| 80 | 0.05650 | 0.01664 | 0.03574 | 0.02498 | 0.489 |
| 90 | 0.05972 | 0.01666 | 0.04700 | 0.02497 | 0.672 |
| 100 | 0.05139 | 0.01676 | 0.03223 | 0.02498 | 0.524 |
| 110 | 0.04912 | 0.01680 | 0.03033 | 0.02498 | 0.533 |
| 120 | 0.05167 | 0.01684 | 0.02687 | 0.02498 | 0.410 |
| 130 | 0.05046 | 0.01685 | 0.02348 | 0.02498 | 0.370 |
| 140 | 0.05660 | 0.01679 | 0.02690 | 0.02498 | 0.324 |
| 150 | 0.05309 | 0.01685 | 0.02690 | 0.02498 | 0.385 |
| 160 | 0.05494 | 0.01685 | 0.02147 | 0.02498 | 0.267 |
| 170 | 0.05780 | 0.01683 | 0.02046 | 0.02499 | 0.215 |
| 180 | 0.05670 | 0.01687 | 0.00828 | 0.02500 | 0.108 |
| 190 | 0.05808 | 0.01687 | 0.00983 | 0.02500 | 0.110 |
| 200 | 0.05554 | 0.01693 | 0.00518 | 0.02500 | 0.095 |
| 210 | 0.05714 | 0.01692 | 0.01289 | 0.02500 | 0.143 |
| 220 | 0.05535 | 0.01693 | 0.01675 | 0.02500 | 0.201 |
| 230 | 0.05315 | 0.01695 | 0.02328 | 0.02499 | 0.323 |
| 240 | 0.05648 | 0.01693 | 0.02587 | 0.02499 | 0.311 |
| 250 | 0.04954 | 0.01699 | 0.02795 | 0.02499 | 0.475 |
| 260 | 0.04949 | 0.01702 | 0.03300 | 0.02498 | 0.585 |
| 270 | 0.04963 | 0.01702 | 0.02950 | 0.02499 | 0.506 |
| 280 | 0.05244 | 0.01701 | 0.03007 | 0.02499 | 0.459 |
| 290 | 0.05916 | 0.01698 | 0.02964 | 0.02499 | 0.329 |
| 300 | 0.05593 | 0.01701 | 0.03120 | 0.02500 | 0.413 |
| 310 | 0.05759 | 0.01699 | 0.03194 | 0.02501 | 0.396 |
| 320 | 0.05457 | 0.01701 | 0.03492 | 0.02500 | 0.516 |
| 330 | 0.05490 | 0.01702 | 0.02675 | 0.02500 | 0.352 |
| 340 | 0.05690 | 0.01700 | 0.03666 | 0.02500 | 0.503 |
| 350 | 0.05698 | 0.01701 | 0.04085 | 0.02501 | 0.594 |
| 360 | 0.06038 | 0.01698 | 0.03957 | 0.02500 | 0.491 |
| 370 | 0.05663 | 0.01700 | 0.03989 | 0.02501 | 0.580 |
| 380 | 0.05790 | 0.01700 | 0.04264 | 0.02500 | 0.614 |
| 390 | 0.05987 | 0.01700 | 0.04213 | 0.02500 | 0.557 |
| 400 | 0.06075 | 0.01700 | 0.03932 | 0.02500 | 0.478 |
| 410 | 0.05797 | 0.01702 | 0.03709 | 0.02500 | 0.490 |
| 420 | 0.05618 | 0.01702 | 0.03457 | 0.02500 | 0.475 |
| 430 | 0.05767 | 0.01702 | 0.03653 | 0.02499 | 0.484 |
| 440 | 0.05927 | 0.01701 | 0.03371 | 0.02499 | 0.398 |
| 450 | 0.05969 | 0.01701 | 0.03220 | 0.02499 | 0.363 |
| 460 | 0.06019 | 0.01702 | 0.03306 | 0.02499 | 0.370 |
| 470 | 0.05752 | 0.01703 | 0.03449 | 0.02499 | 0.446 |
| 480 | 0.05979 | 0.01703 | 0.03515 | 0.02499 | 0.415 |
| 490 | 0.06146 | 0.01702 | 0.03429 | 0.02499 | 0.369 |
| 500 | 0.06082 | 0.01703 | 0.03725 | 0.02499 | 0.436 |
| 510 | 0.06503 | 0.01702 | 0.03765 | 0.02499 | 0.365 |
| 520 | 0.06512 | 0.01702 | 0.03094 | 0.02499 | 0.258 |
| 530 | 0.06363 | 0.01702 | 0.02887 | 0.02499 | 0.250 |
| 540 | 0.05925 | 0.01705 | 0.03263 | 0.02499 | 0.379 |
| 550 | 0.06022 | 0.01705 | 0.03527 | 0.02499 | 0.410 |
| 560 | 0.06395 | 0.01703 | 0.03818 | 0.02499 | 0.394 |
| 570 | 0.06105 | 0.01704 | 0.03994 | 0.02499 | 0.485 |
| 580 | 0.05683 | 0.01706 | 0.04346 | 0.02500 | 0.659 |
| 590 | 0.05738 | 0.01706 | 0.04448 | 0.02499 | 0.670 |
| 600 | 0.05393 | 0.01708 | 0.04433 | 0.02499 | 0.751 |
| 610 | 0.05296 | 0.01707 | 0.04519 | 0.02499 | 0.797 |
| 620 | 0.05355 | 0.01707 | 0.04676 | 0.02500 | 0.822 |
| 630 | 0.05701 | 0.01706 | 0.04696 | 0.02500 | 0.740 |
| 640 | 0.06009 | 0.01705 | 0.05001 | 0.02500 | 0.739 |
| 650 | 0.05933 | 0.01706 | 0.05015 | 0.02500 | 0.762 |
| 660 | 0.05920 | 0.01706 | 0.05461 | 0.02500 | 0.880 |
| 670 | 0.05658 | 0.01707 | 0.05242 | 0.02500 | 0.891 |
| 680 | 0.05867 | 0.01707 | 0.05631 | 0.02500 | 0.938 |
| 690 | 0.06164 | 0.01706 | 0.05263 | 0.02500 | 0.766 |
| 700 | 0.06289 | 0.01706 | 0.05056 | 0.02500 | 0.684 |
| 710 | 0.06342 | 0.01707 | 0.04653 | 0.02500 | 0.577 |
| 720 | 0.06293 | 0.01707 | 0.04330 | 0.02500 | 0.517 |
| 730 | 0.06067 | 0.01708 | 0.04331 | 0.02500 | 0.566 |
| 740 | 0.06239 | 0.01706 | 0.04021 | 0.02500 | 0.464 |
| 750 | 0.06524 | 0.01705 | 0.04524 | 0.02500 | 0.509 |
| 760 | 0.06434 | 0.01706 | 0.04501 | 0.02500 | 0.523 |
| 770 | 0.06314 | 0.01706 | 0.04744 | 0.02500 | 0.604 |
| 780 | 0.06171 | 0.01707 | 0.04482 | 0.02500 | 0.577 |
| 790 | 0.06076 | 0.01707 | 0.04101 | 0.02500 | 0.514 |
| 800 | 0.06424 | 0.01707 | 0.04297 | 0.02500 | 0.482 |
| 810 | 0.06624 | 0.01706 | 0.04385 | 0.02499 | 0.459 |
| 820 | 0.06635 | 0.01705 | 0.04096 | 0.02499 | 0.401 |
| 830 | 0.06604 | 0.01705 | 0.04186 | 0.02500 | 0.424 |
| 840 | 0.06566 | 0.01706 | 0.04152 | 0.02500 | 0.425 |
| 850 | 0.06420 | 0.01707 | 0.04105 | 0.02500 | 0.445 |
| 860 | 0.06550 | 0.01707 | 0.04163 | 0.02500 | 0.430 |
| 870 | 0.06651 | 0.01706 | 0.04376 | 0.02500 | 0.452 |
| 880 | 0.06492 | 0.01707 | 0.04184 | 0.02500 | 0.446 |
| 890 | 0.06616 | 0.01706 | 0.04263 | 0.02501 | 0.437 |
| 900 | 0.06835 | 0.01705 | 0.03994 | 0.02500 | 0.348 |
| 910 | 0.06657 | 0.01705 | 0.04215 | 0.02500 | 0.420 |
| 920 | 0.06514 | 0.01706 | 0.04028 | 0.02500 | 0.411 |
| 930 | 0.06841 | 0.01705 | 0.03962 | 0.02501 | 0.342 |
| 940 | 0.07128 | 0.01705 | 0.04288 | 0.02501 | 0.348 |
| 950 | 0.07256 | 0.01704 | 0.04161 | 0.02501 | 0.307 |
| 960 | 0.07179 | 0.01705 | 0.04291 | 0.02501 | 0.340 |
| 970 | 0.07192 | 0.01705 | 0.04150 | 0.02501 | 0.315 |
| 980 | 0.07246 | 0.01705 | 0.03752 | 0.02500 | 0.248 |
| 990 | 0.06959 | 0.01705 | 0.03636 | 0.02500 | 0.272 |
| 1000 | 0.06847 | 0.01706 | 0.03677 | 0.02501 | 0.295 |
| 1010 | 0.06913 | 0.01706 | 0.03643 | 0.02501 | 0.280 |
| 1020 | 0.06751 | 0.01707 | 0.03623 | 0.02500 | 0.302 |
| 1030 | 0.06558 | 0.01708 | 0.03469 | 0.02500 | 0.308 |
| 1040 | 0.06447 | 0.01708 | 0.03450 | 0.02500 | 0.322 |
| 1050 | 0.06461 | 0.01708 | 0.03179 | 0.02500 | 0.278 |
| 1060 | 0.06452 | 0.01708 | 0.02921 | 0.02500 | 0.244 |
| 1070 | 0.06500 | 0.01708 | 0.03155 | 0.02500 | 0.269 |
| 1080 | 0.06290 | 0.01709 | 0.02944 | 0.02500 | 0.269 |
| 1090 | 0.06255 | 0.01710 | 0.03067 | 0.02500 | 0.293 |
| 1100 | 0.06284 | 0.01710 | 0.03035 | 0.02500 | 0.283 |
| 1110 | 0.06284 | 0.01710 | 0.03025 | 0.02501 | 0.282 |
| 1120 | 0.06369 | 0.01710 | 0.03121 | 0.02501 | 0.284 |
| 1130 | 0.06451 | 0.01710 | 0.03095 | 0.02501 | 0.268 |
| 1140 | 0.06502 | 0.01710 | 0.03309 | 0.02501 | 0.292 |
| 1150 | 0.06475 | 0.01711 | 0.03582 | 0.02501 | 0.340 |
| 1160 | 0.06510 | 0.01711 | 0.03604 | 0.02501 | 0.338 |
| 1170 | 0.06413 | 0.01711 | 0.03826 | 0.02501 | 0.393 |
| 1180 | 0.06440 | 0.01711 | 0.03733 | 0.02501 | 0.372 |
| 1190 | 0.06442 | 0.01711 | 0.03584 | 0.02501 | 0.346 |
| 1200 | 0.06455 | 0.01711 | 0.03598 | 0.02501 | 0.346 |
| 1210 | 0.06340 | 0.01711 | 0.03586 | 0.02502 | 0.364 |
| 1220 | 0.06217 | 0.01712 | 0.03883 | 0.02501 | 0.441 |
| 1230 | 0.06252 | 0.01712 | 0.03788 | 0.02501 | 0.416 |
| 1240 | 0.06268 | 0.01712 | 0.04024 | 0.02502 | 0.459 |
| 1250 | 0.06409 | 0.01712 | 0.04416 | 0.02502 | 0.511 |
| 1260 | 0.06354 | 0.01712 | 0.04565 | 0.02501 | 0.555 |
| 1270 | 0.06340 | 0.01713 | 0.04415 | 0.02501 | 0.525 |
| 1280 | 0.06307 | 0.01713 | 0.04078 | 0.02501 | 0.462 |
| 1290 | 0.06283 | 0.01713 | 0.04148 | 0.02501 | 0.481 |
| 1300 | 0.06287 | 0.01713 | 0.04162 | 0.02501 | 0.483 |
| 1310 | 0.06338 | 0.01713 | 0.04209 | 0.02501 | 0.483 |
| 1320 | 0.06267 | 0.01713 | 0.03999 | 0.02501 | 0.454 |
| 1330 | 0.06004 | 0.01713 | 0.04192 | 0.02501 | 0.550 |
| 1340 | 0.06016 | 0.01713 | 0.04119 | 0.02501 | 0.531 |
| 1350 | 0.06067 | 0.01713 | 0.04034 | 0.02501 | 0.502 |
| 1360 | 0.06237 | 0.01713 | 0.03874 | 0.02501 | 0.436 |
| 1370 | 0.06177 | 0.01714 | 0.03712 | 0.02501 | 0.416 |
| 1380 | 0.06172 | 0.01714 | 0.03591 | 0.02501 | 0.395 |
| 1390 | 0.06113 | 0.01714 | 0.03280 | 0.02501 | 0.350 |
| 1400 | 0.06107 | 0.01714 | 0.03352 | 0.02501 | 0.364 |
| 1410 | 0.06132 | 0.01714 | 0.03189 | 0.02501 | 0.332 |
| 1420 | 0.05925 | 0.01714 | 0.03184 | 0.02501 | 0.366 |
| 1430 | 0.05819 | 0.01715 | 0.03448 | 0.02501 | 0.434 |
| 1440 | 0.05887 | 0.01715 | 0.03331 | 0.02501 | 0.399 |
| 1450 | 0.05826 | 0.01715 | 0.03266 | 0.02501 | 0.399 |
| 1460 | 0.05859 | 0.01715 | 0.03483 | 0.02501 | 0.433 |
| 1470 | 0.05993 | 0.01715 | 0.03499 | 0.02501 | 0.411 |
| 1480 | 0.05852 | 0.01715 | 0.03561 | 0.02501 | 0.450 |
| 1490 | 0.05818 | 0.01715 | 0.03502 | 0.02501 | 0.445 |
| 1500 | 0.05766 | 0.01715 | 0.03637 | 0.02501 | 0.483 |
| 1510 | 0.05909 | 0.01715 | 0.03478 | 0.02501 | 0.423 |
| 1520 | 0.05842 | 0.01715 | 0.03702 | 0.02501 | 0.480 |
| 1530 | 0.05805 | 0.01715 | 0.03419 | 0.02501 | 0.431 |
| 1540 | 0.06034 | 0.01715 | 0.03129 | 0.02501 | 0.338 |
| 1550 | 0.05966 | 0.01715 | 0.03242 | 0.02501 | 0.369 |
| 1560 | 0.05888 | 0.01715 | 0.03415 | 0.02501 | 0.415 |
| 1570 | 0.05815 | 0.01715 | 0.03624 | 0.02501 | 0.470 |
| 1580 | 0.05730 | 0.01716 | 0.03627 | 0.02501 | 0.488 |
| 1590 | 0.05708 | 0.01716 | 0.03675 | 0.02501 | 0.503 |
| 1600 | 0.05815 | 0.01715 | 0.03636 | 0.02501 | 0.472 |
| 1610 | 0.05818 | 0.01716 | 0.03537 | 0.02501 | 0.452 |
| 1620 | 0.05891 | 0.01716 | 0.03378 | 0.02501 | 0.407 |
| 1630 | 0.05886 | 0.01716 | 0.03240 | 0.02501 | 0.383 |
| 1640 | 0.05993 | 0.01716 | 0.03519 | 0.02501 | 0.415 |
| 1650 | 0.06062 | 0.01716 | 0.03402 | 0.02501 | 0.380 |
| 1660 | 0.05929 | 0.01716 | 0.03434 | 0.02501 | 0.411 |
| 1670 | 0.05945 | 0.01717 | 0.03559 | 0.02501 | 0.431 |
| 1680 | 0.06008 | 0.01716 | 0.03362 | 0.02501 | 0.383 |
| 1690 | 0.06044 | 0.01716 | 0.03316 | 0.02501 | 0.369 |
| 1700 | 0.05972 | 0.01716 | 0.03413 | 0.02501 | 0.399 |
| 1710 | 0.05936 | 0.01716 | 0.03416 | 0.02501 | 0.406 |
| 1720 | 0.05901 | 0.01716 | 0.03524 | 0.02501 | 0.433 |
| 1730 | 0.06051 | 0.01716 | 0.03542 | 0.02501 | 0.408 |
| 1740 | 0.06016 | 0.01716 | 0.03562 | 0.02501 | 0.418 |
| 1750 | 0.06030 | 0.01717 | 0.03724 | 0.02501 | 0.447 |
| 1760 | 0.06077 | 0.01716 | 0.03904 | 0.02501 | 0.474 |
| 1770 | 0.05974 | 0.01716 | 0.03866 | 0.02501 | 0.487 |
| 1780 | 0.05987 | 0.01717 | 0.03829 | 0.02501 | 0.477 |
| 1790 | 0.06026 | 0.01717 | 0.03795 | 0.02501 | 0.462 |
| 1800 | 0.05867 | 0.01717 | 0.03996 | 0.02501 | 0.537 |
| 1810 | 0.05916 | 0.01717 | 0.03994 | 0.02501 | 0.526 |
| 1820 | 0.05959 | 0.01717 | 0.04155 | 0.02501 | 0.552 |
| 1830 | 0.05922 | 0.01717 | 0.04136 | 0.02502 | 0.556 |
| 1840 | 0.05857 | 0.01717 | 0.03977 | 0.02502 | 0.536 |
| 1850 | 0.05807 | 0.01717 | 0.03949 | 0.02502 | 0.540 |
| 1860 | 0.05908 | 0.01717 | 0.03961 | 0.02502 | 0.521 |
| 1870 | 0.05956 | 0.01717 | 0.04091 | 0.02502 | 0.539 |
| 1880 | 0.05967 | 0.01717 | 0.03898 | 0.02501 | 0.495 |
| 1890 | 0.05883 | 0.01717 | 0.03788 | 0.02501 | 0.490 |
| 1900 | 0.05828 | 0.01717 | 0.03809 | 0.02501 | 0.506 |
| 1910 | 0.05825 | 0.01717 | 0.03956 | 0.02501 | 0.538 |
| 1920 | 0.05879 | 0.01717 | 0.04100 | 0.02501 | 0.557 |
| 1930 | 0.05908 | 0.01717 | 0.04153 | 0.02501 | 0.563 |
| 1940 | 0.05838 | 0.01717 | 0.04138 | 0.02501 | 0.575 |
| 1950 | 0.05676 | 0.01718 | 0.03945 | 0.02501 | 0.568 |
| 1960 | 0.05671 | 0.01718 | 0.04059 | 0.02501 | 0.595 |
| 1970 | 0.05756 | 0.01718 | 0.04082 | 0.02501 | 0.581 |
| 1980 | 0.05731 | 0.01718 | 0.04020 | 0.02501 | 0.573 |
| 1990 | 0.05865 | 0.01717 | 0.04040 | 0.02501 | 0.548 |
| 2000 | 0.05809 | 0.01718 | 0.04002 | 0.02501 | 0.552 |
| 2010 | 0.05745 | 0.01718 | 0.04008 | 0.02501 | 0.567 |
| 2020 | 0.05736 | 0.01718 | 0.04040 | 0.02501 | 0.576 |
| 2030 | 0.05748 | 0.01718 | 0.03906 | 0.02501 | 0.544 |
| 2040 | 0.05733 | 0.01718 | 0.03974 | 0.02501 | 0.562 |
| 2050 | 0.05738 | 0.01718 | 0.04054 | 0.02501 | 0.579 |
| 2060 | 0.05655 | 0.01718 | 0.04007 | 0.02501 | 0.587 |
| 2070 | 0.05629 | 0.01718 | 0.03901 | 0.02501 | 0.569 |
| 2080 | 0.05604 | 0.01719 | 0.03856 | 0.02502 | 0.565 |
| 2090 | 0.05511 | 0.01719 | 0.03824 | 0.02502 | 0.578 |
| 2100 | 0.05445 | 0.01718 | 0.04062 | 0.02502 | 0.649 |
| 2110 | 0.05503 | 0.01719 | 0.04002 | 0.02502 | 0.621 |
| 2120 | 0.05515 | 0.01719 | 0.04001 | 0.02502 | 0.618 |
| 2130 | 0.05514 | 0.01719 | 0.04039 | 0.02502 | 0.627 |
| 2140 | 0.05553 | 0.01718 | 0.04119 | 0.02502 | 0.636 |
| 2150 | 0.05544 | 0.01718 | 0.04166 | 0.02502 | 0.650 |
| 2160 | 0.05573 | 0.01718 | 0.04182 | 0.02502 | 0.647 |
| 2170 | 0.05530 | 0.01718 | 0.04297 | 0.02502 | 0.685 |
| 2180 | 0.05559 | 0.01718 | 0.04413 | 0.02502 | 0.706 |
| 2190 | 0.05492 | 0.01718 | 0.04410 | 0.02502 | 0.721 |
| 2200 | 0.05455 | 0.01718 | 0.04438 | 0.02502 | 0.738 |
| 2210 | 0.05458 | 0.01718 | 0.04554 | 0.02502 | 0.766 |
| 2220 | 0.05472 | 0.01718 | 0.04480 | 0.02502 | 0.744 |
| 2230 | 0.05531 | 0.01718 | 0.04446 | 0.02502 | 0.721 |
| 2240 | 0.05432 | 0.01718 | 0.04462 | 0.02502 | 0.749 |
| 2250 | 0.05350 | 0.01718 | 0.04326 | 0.02502 | 0.736 |
| 2260 | 0.05342 | 0.01718 | 0.04360 | 0.02502 | 0.746 |
| 2270 | 0.05313 | 0.01718 | 0.04349 | 0.02502 | 0.751 |
| 2280 | 0.05429 | 0.01718 | 0.04464 | 0.02502 | 0.751 |
| 2290 | 0.05371 | 0.01718 | 0.04511 | 0.02502 | 0.777 |
| 2300 | 0.05350 | 0.01718 | 0.04487 | 0.02502 | 0.776 |
| 2310 | 0.05377 | 0.01718 | 0.04451 | 0.02502 | 0.760 |
| 2320 | 0.05386 | 0.01718 | 0.04329 | 0.02502 | 0.728 |
| 2330 | 0.05415 | 0.01718 | 0.04338 | 0.02502 | 0.723 |
| 2340 | 0.05413 | 0.01718 | 0.04389 | 0.02502 | 0.736 |
| 2350 | 0.05430 | 0.01718 | 0.04435 | 0.02502 | 0.743 |
| 2360 | 0.05390 | 0.01719 | 0.04474 | 0.02501 | 0.763 |
| 2370 | 0.05467 | 0.01718 | 0.04405 | 0.02501 | 0.726 |
| 2380 | 0.05517 | 0.01718 | 0.04307 | 0.02501 | 0.690 |
| 2390 | 0.05480 | 0.01718 | 0.04303 | 0.02501 | 0.698 |
| 2400 | 0.05526 | 0.01718 | 0.04231 | 0.02501 | 0.670 |
| 2410 | 0.05524 | 0.01718 | 0.04248 | 0.02501 | 0.674 |
| 2420 | 0.05514 | 0.01718 | 0.04231 | 0.02501 | 0.672 |
| 2430 | 0.05490 | 0.01718 | 0.04262 | 0.02501 | 0.686 |
| 2440 | 0.05472 | 0.01718 | 0.04234 | 0.02501 | 0.683 |
| 2450 | 0.05480 | 0.01718 | 0.04197 | 0.02501 | 0.672 |
| 2460 | 0.05413 | 0.01718 | 0.04251 | 0.02501 | 0.702 |
| 2470 | 0.05418 | 0.01718 | 0.04283 | 0.02501 | 0.708 |
| 2480 | 0.05462 | 0.01718 | 0.04283 | 0.02501 | 0.698 |
| 2490 | 0.05436 | 0.01718 | 0.04262 | 0.02501 | 0.699 |
| 2500 | 0.05471 | 0.01718 | 0.04269 | 0.02501 | 0.692 |
| 2510 | 0.05518 | 0.01718 | 0.04299 | 0.02501 | 0.688 |
| 2520 | 0.05592 | 0.01718 | 0.04284 | 0.02501 | 0.666 |
| 2530 | 0.05622 | 0.01717 | 0.04250 | 0.02501 | 0.651 |
| 2540 | 0.05625 | 0.01717 | 0.04157 | 0.02501 | 0.628 |
| 2550 | 0.05617 | 0.01718 | 0.04153 | 0.02501 | 0.629 |
| 2560 | 0.05601 | 0.01718 | 0.04256 | 0.02501 | 0.658 |
| 2570 | 0.05583 | 0.01718 | 0.04221 | 0.02501 | 0.654 |
| 2580 | 0.05531 | 0.01718 | 0.04334 | 0.02501 | 0.693 |
| 2590 | 0.05527 | 0.01718 | 0.04299 | 0.02502 | 0.686 |
| 2600 | 0.05468 | 0.01718 | 0.04385 | 0.02502 | 0.721 |
| 2610 | 0.05442 | 0.01718 | 0.04422 | 0.02502 | 0.737 |
| 2620 | 0.05440 | 0.01718 | 0.04476 | 0.02501 | 0.751 |
| 2630 | 0.05384 | 0.01718 | 0.04463 | 0.02501 | 0.761 |
| 2640 | 0.05436 | 0.01718 | 0.04424 | 0.02501 | 0.739 |
| 2650 | 0.05507 | 0.01718 | 0.04391 | 0.02501 | 0.713 |
| 2660 | 0.05532 | 0.01718 | 0.04473 | 0.02501 | 0.727 |
| 2670 | 0.05501 | 0.01718 | 0.04449 | 0.02501 | 0.729 |
| 2680 | 0.05523 | 0.01718 | 0.04534 | 0.02501 | 0.744 |
| 2690 | 0.05483 | 0.01718 | 0.04580 | 0.02501 | 0.766 |
| 2700 | 0.05507 | 0.01718 | 0.04640 | 0.02501 | 0.775 |
| 2710 | 0.05504 | 0.01718 | 0.04650 | 0.02501 | 0.778 |
| 2720 | 0.05485 | 0.01718 | 0.04694 | 0.02501 | 0.794 |
| 2730 | 0.05530 | 0.01718 | 0.04623 | 0.02501 | 0.765 |
| 2740 | 0.05510 | 0.01718 | 0.04514 | 0.02501 | 0.743 |
| 2750 | 0.05532 | 0.01718 | 0.04531 | 0.02501 | 0.742 |
| 2760 | 0.05544 | 0.01718 | 0.04514 | 0.02501 | 0.734 |
| 2770 | 0.05557 | 0.01718 | 0.04511 | 0.02501 | 0.730 |
| 2780 | 0.05543 | 0.01718 | 0.04614 | 0.02501 | 0.759 |
| 2790 | 0.05538 | 0.01718 | 0.04565 | 0.02501 | 0.748 |
| 2800 | 0.05533 | 0.01718 | 0.04584 | 0.02501 | 0.754 |
| 2810 | 0.05546 | 0.01718 | 0.04614 | 0.02501 | 0.759 |
| 2820 | 0.05506 | 0.01718 | 0.04596 | 0.02501 | 0.764 |
| 2830 | 0.05499 | 0.01718 | 0.04593 | 0.02501 | 0.765 |
| 2840 | 0.05503 | 0.01718 | 0.04589 | 0.02501 | 0.763 |
| 2850 | 0.05513 | 0.01718 | 0.04613 | 0.02501 | 0.767 |
| 2860 | 0.05491 | 0.01718 | 0.04649 | 0.02501 | 0.781 |
| 2870 | 0.05465 | 0.01718 | 0.04690 | 0.02501 | 0.798 |
| 2880 | 0.05456 | 0.01718 | 0.04681 | 0.02501 | 0.798 |
| 2890 | 0.05434 | 0.01718 | 0.04644 | 0.02501 | 0.795 |
| 2900 | 0.05448 | 0.01718 | 0.04678 | 0.02501 | 0.800 |
| 2910 | 0.05437 | 0.01718 | 0.04620 | 0.02501 | 0.788 |
| 2920 | 0.05439 | 0.01718 | 0.04617 | 0.02501 | 0.786 |
| 2930 | 0.05427 | 0.01718 | 0.04577 | 0.02501 | 0.779 |
| 2940 | 0.05404 | 0.01718 | 0.04602 | 0.02501 | 0.791 |
| 2950 | 0.05411 | 0.01718 | 0.04606 | 0.02501 | 0.791 |
| 2960 | 0.05391 | 0.01718 | 0.04602 | 0.02501 | 0.795 |
| 2970 | 0.05424 | 0.01718 | 0.04599 | 0.02501 | 0.786 |
| 2980 | 0.05425 | 0.01718 | 0.04623 | 0.02501 | 0.792 |
| 2990 | 0.05417 | 0.01718 | 0.04604 | 0.02501 | 0.789 |
| 3000 | 0.05417 | 0.01719 | 0.04572 | 0.02501 | 0.781 |

<sup>a</sup> $\beta$ indicates the regression coefficient
<sup>b</sup> SE( $\beta$ ) indicates the standard error of the regression coefficient

**Supplementary Table 10:** Effect estimates of RVGRS950 and RVGRS950*^LDLR-^* on CAD across tertiles of FRS in the UK Biobank.

| FRS tertile<sup>a</sup> | OR (95% CI) | P-value | P<sub>interaction</sub> |
| --- | --- | --- | --- |
| RVGRS950 |  |  |  |
| 1 | 1.16 (1.06-1.27) | 8.4x10<sup>-4</sup> | 0.015 |
| 2 | 1.10 (1.03-1.17) | 2.3x10<sup>-3</sup> |  |
| 3 | 1.03 (0.99-1.08) | 0.17 |  |
| RVGRS950<sup>LDLR-</sup> |  |  |  |
| 1 | 1.15 (1.05-1.26) | 2.0x10<sup>-3</sup> | 0.019 |
| 2 | 1.09 (1.02-1.16) | 6.2x10<sup>-3</sup> |  |
| 3 | 1.03 (0.98-1.08) | 0.29 |  |

<sup>a</sup> Effect estimates and P-values for each tertile were determined using age, age<sup>2</sup>, sex, and the first 20 principle components as covariates

**Supplementary Table 11:** Effect estimates of CVGRS on tertiles on CAD across tertile of FRS in the UK Biobank.

| FRS tertile<sup>a</sup> | OR (95% CI) | P-value | P<sub>interaction</sub> |
| --- | --- | --- | --- |
| 1 | 1.20 (1.14-1.26) | 5.3x10<sup>-12</sup> | 0.65 |
| 2 | 1.14 (1.11-1.18) | 2.2x10<sup>-14</sup> |  |
| 3 | 1.17 (1.13-1.20) | < 2.2x10<sup>-16</sup> |  |

<sup>a</sup> Effect estimates and P-values for each tertile were determined using age, age<sup>2</sup>, sex, and the first 20 principle components as covariates

**Supplementary Table 12:**
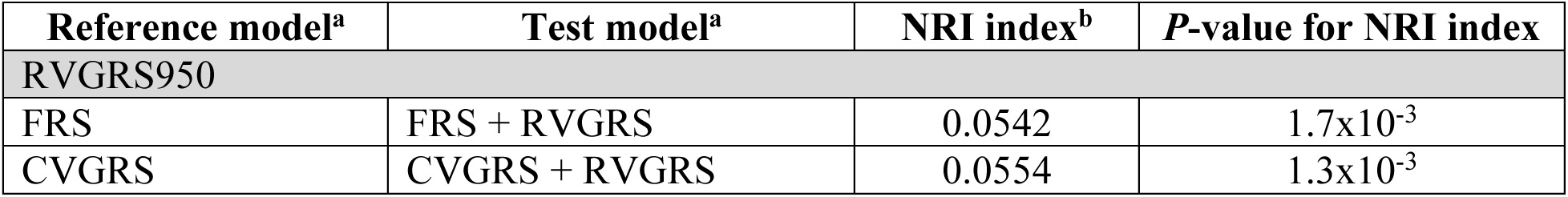
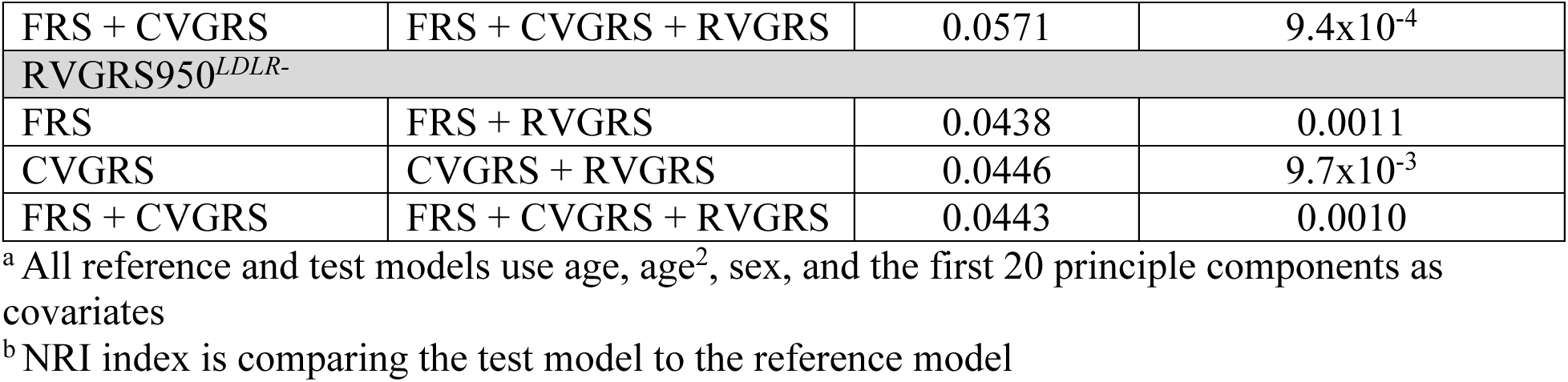
Net reclassification improvement index when incorporating RVGRS950 and RVGRS950*^LDLR-^* to risk models.

---

**Supplementary Figure 1:**
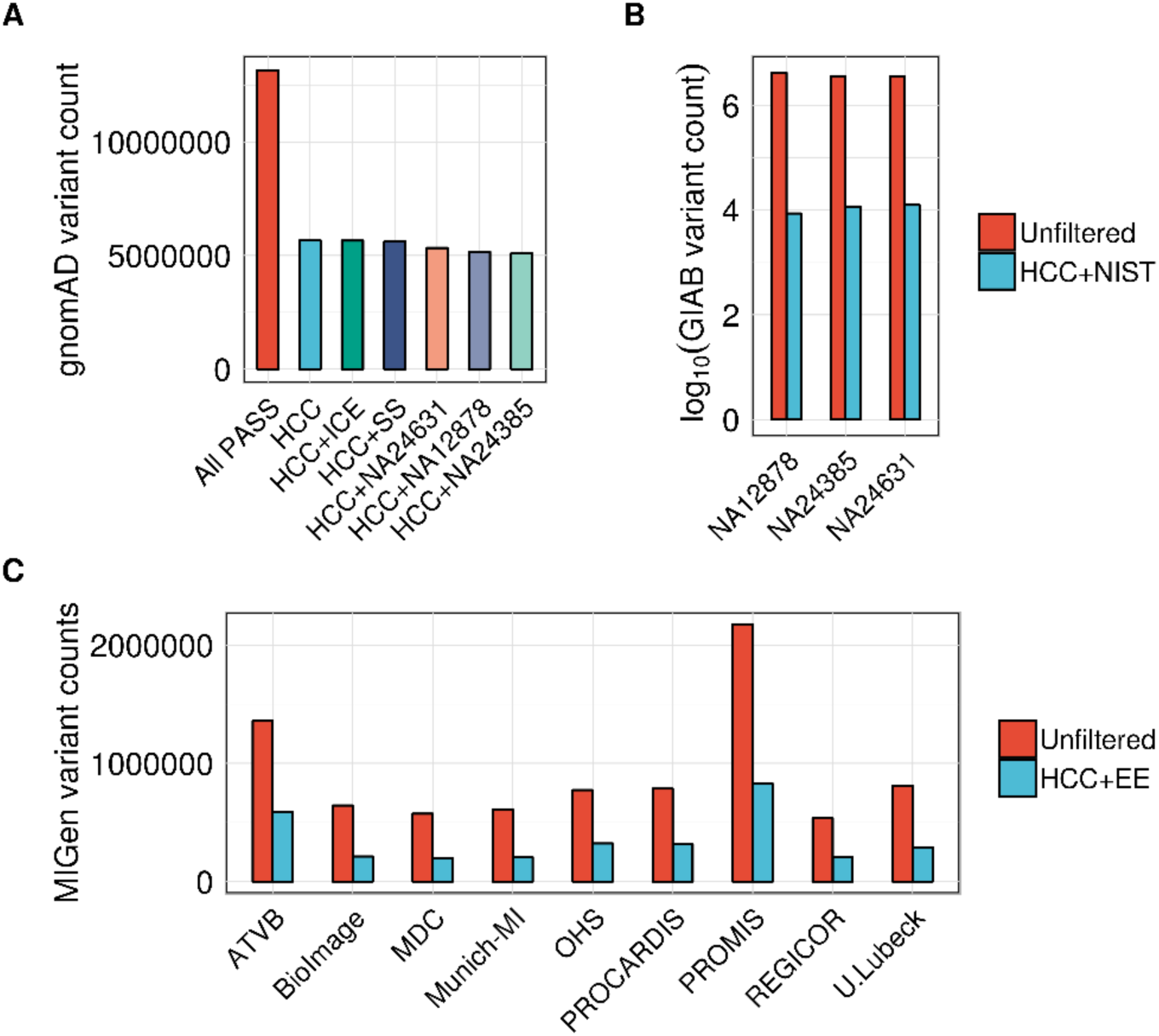
Number of variant sites remaining in (A) gnomAD, (B) GIAB, and (C) MIGen after intersection with different region intervals. For (A), HCC+ICE: intersection sites between high-coverage coding sites (defined in Supplementary Section 9B) and the ICE-capture exome enrichment kit; HCC+SS: intersection between HCC sites and SureSelect (SS) Human All Exon v.2 Kit; HCC+NA24631: intersection between HCC and NIST high-cofidence sites for GIAB sample NA24631; HCC+NA12878: intersection between HCC and NIST high-confidence sites for GIAB sample NA12878; HCC+NA24385: intersection between HCC and NIST high-confidence sites for GIAB sample NA24631. For (B), HCC+NIST: intersection between HCC sites and each respective NIST high-confidence interval site for each GIAB sample. For (C), HCC+EE: intersection between HCC sites and intervals from the exome enrichment (EE) kit used for a given MIGen cohort. HCC indicates high coverage coding sites, SS indicates SureSelect, NIST indicates National Institute of Standards and Technology, and GIAB indicates Genome In A Bottle.

**Supplementary Figure 2:**
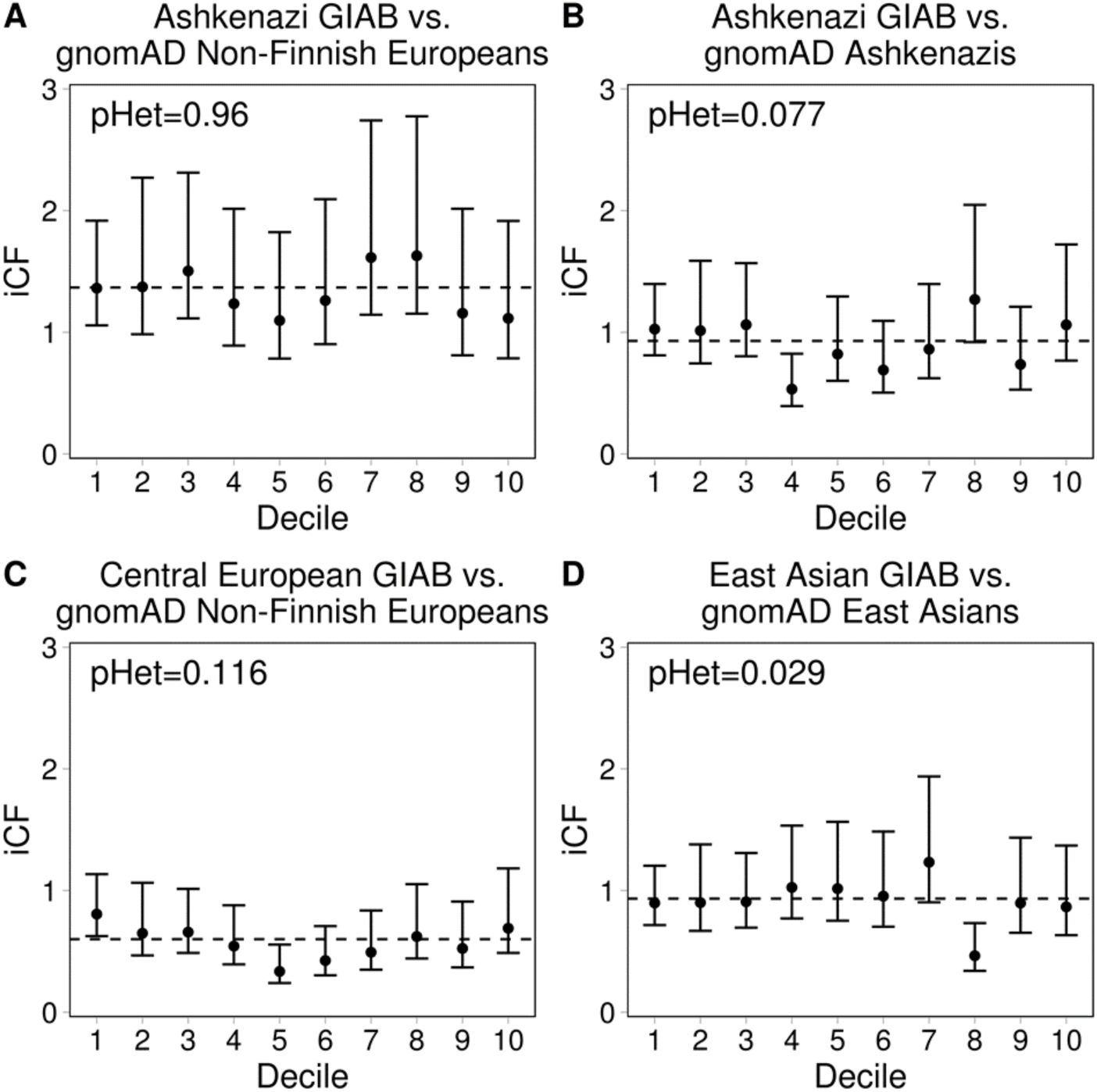
GIAB iCF values for gene groups stratified by decile of gene constraint score. Ethnic comparisons include the Ashkenazi GIAB sample (NA24385) with gnomAD non-Finnish Europeans and gnomAD Ashkenazis (**A** and **B**), the Central European GIAB sample (NA12878) with gnomAD Non-Finnish Europeans (**C**), and the East Asian GIAB sample (NA24631) with gnomAD East Asians population (**D**). All genes were grouped by decile according to their missense constraint metric (Lek *et al*., 2016). iCF values were calculated for each decile according to equation 6. Error bars depict 95% CI and dashed lines represent the iCF across all deciles (i.e. exome-wide iCF). *P*-values for heterogeneity (pHet) across deciles were calculated using a fixed effect model. Any *P*-value < 0.0125 (0.05/4) was considered significant.

**Supplementary Figure 3:**
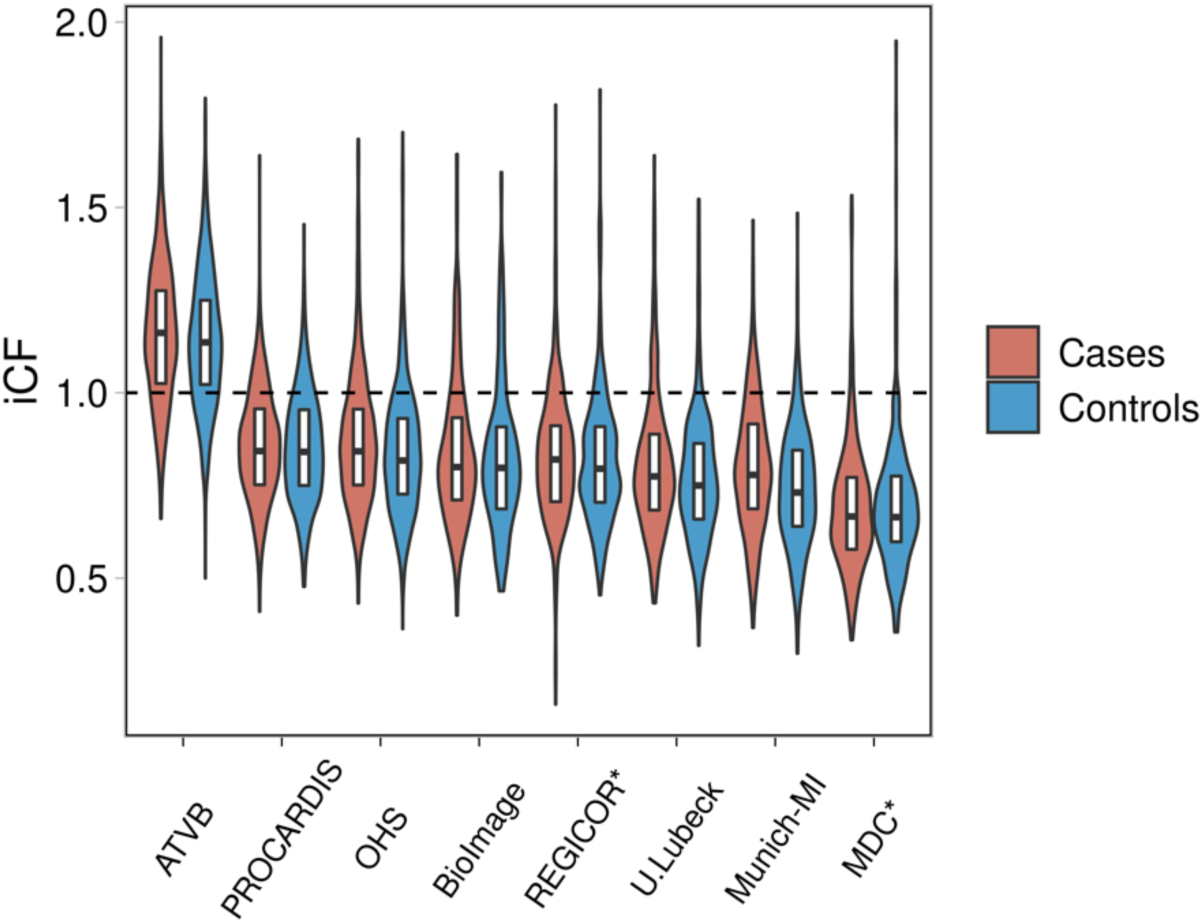
iCF values for cases and controls across all discovery MIGen cohorts. Distribution of iCF values are shown for case (red) and control (blue) participants across 8 MIGen cohorts. Violins demonstrate the spread of iCF values. The horizontal line in each boxplot indicate the median iCF value while the top and bottom lines represent the 75^th^ and 25^th^ percentiles of the iCF distribution, respectively. Length of boxplot represents the inter-quartile range of iCF values. Cohorts marked with an asterisk were found to have a significantly different (P<0.05) iCF distribution between cases and controls after adjusting for sex and the first 20 principal components (age was not available as a phenotype in MIGen). The dashed line represents an equal mutation load between MIGen cases or controls versus gnomAD non-Finnish Europeans. iCF indicates individual correction factor.

**Supplementary Figure 4:**
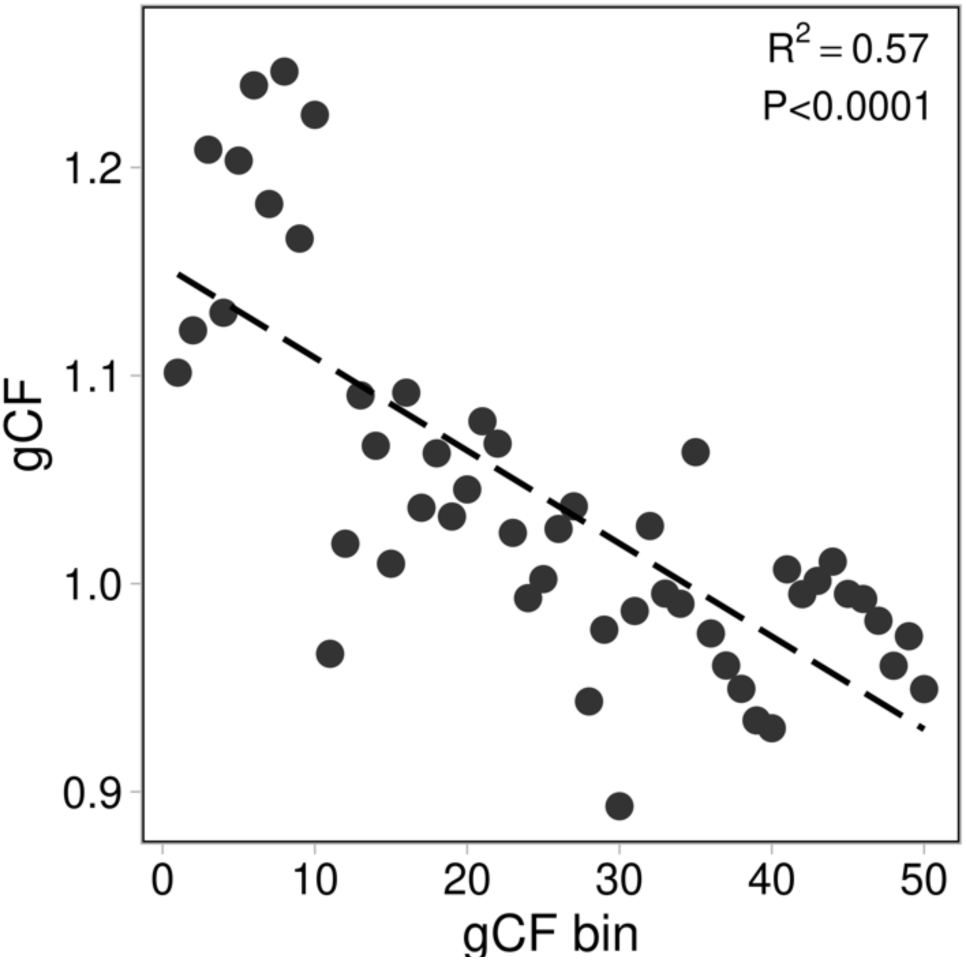
gCF values across 50 gene bins derived from 3,352 healthy MIGen controls. gCF values were computed as the ratio of observed count to iCF-adjusted expected count across all protein-coding genes that were organized into one of 50 gene bins according to quintile of iCF-adjsuted expected count and decile of P-value obtained from the rare variant association test using burden of rare pathogenic alleles in the ranking cohort consisting of the remaining 2,730 healthy control participants in MIGen as “cases” and gnomAD non-Finnish Europeans as controls. R^2^ and significance was evaluated through a linear regression model. gCF indicates gene correction factor.

**Supplementary Figure 5:**
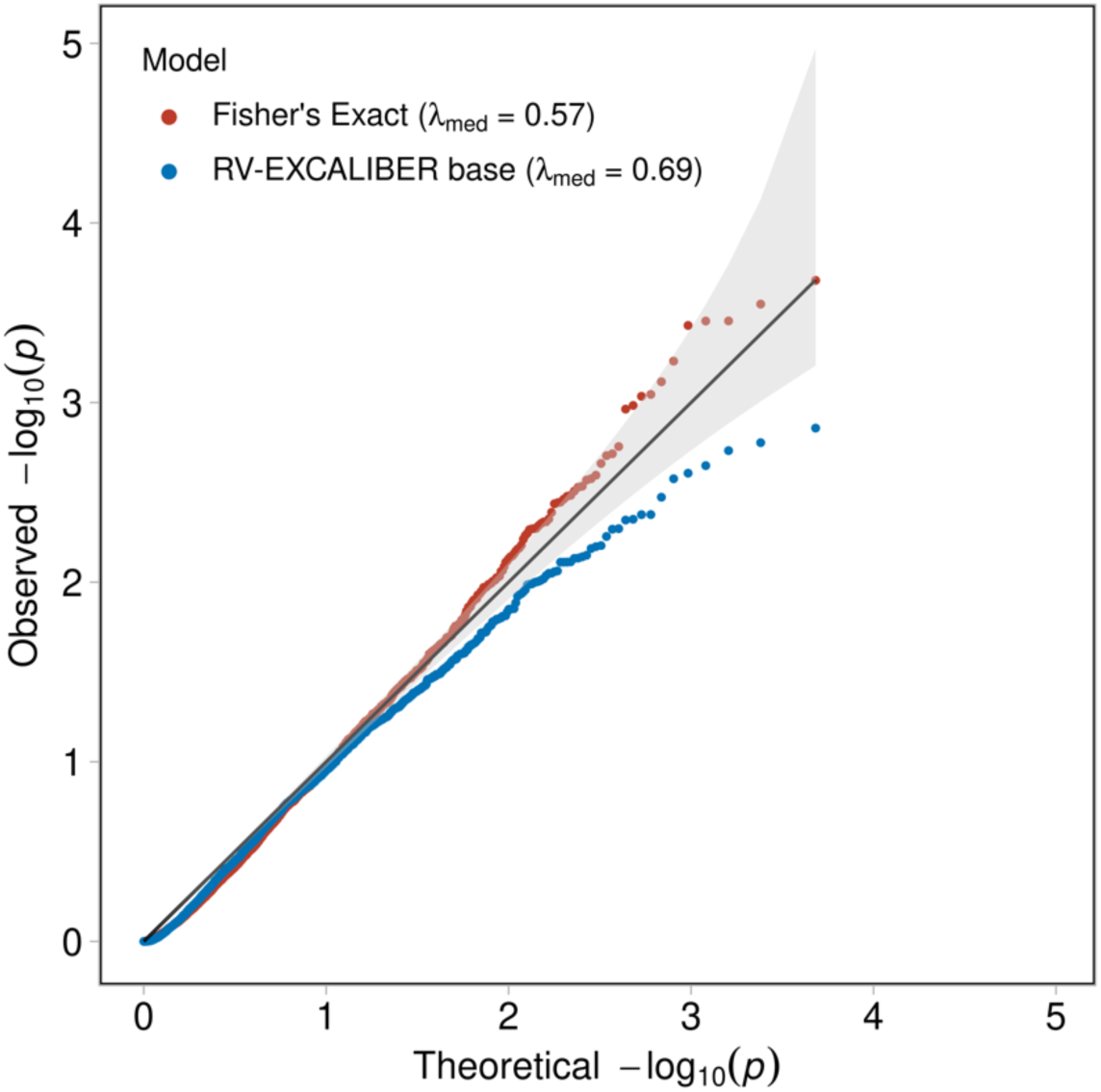
Quantile-quantile plots for gene-based test statistics generated using RV-EXCALIBER base and a Fisher’s Exact test for 3,352 healthy control participants in MIGen. RV-EXCALIBER base only accounts for LD between rare variants does not incorporate ‘iCF’ or ‘iCF and gCF’-adjusted expected counts. A total of 4,815 protein-coding genes with observed counts ≥1 in MIGen and expected counts ≥10 in gnomAD non-Finnish Europeans were used to conduct a gene burden test using rare pathogenic alleles according to both RV-EXCALIBER base and a Fisher’s Exact tests. A total of 3,352 healthy control participants were used as “cases” in MIGen and gnomAD non-Finnish Europeans were used as controls. Genomic inflation estimates were evaluated at the median (*λ_med_*). The solid black line indicates an expected uniform distribution of *P*-values under the null and the shaded region represent 95% confidence interval of the expected distribution.

**Supplementary Figure 6:**
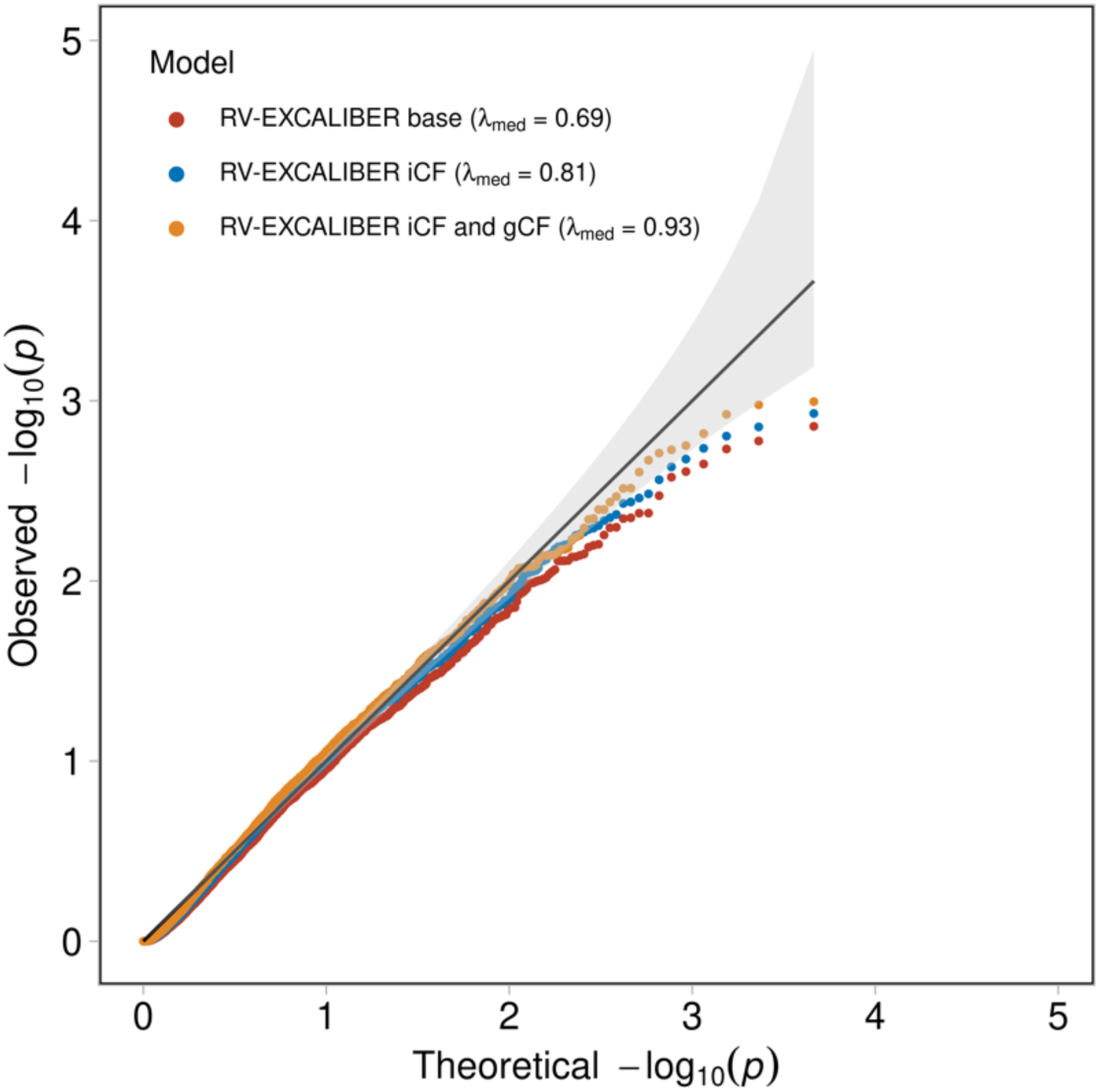
Quantile-quantile plots for gene-based test statistics generated using RV-EXCALIBER base, RV-ECALIBER iCF and RV-EXCALIBER iCF+gCF (i.e. fully adjusted model) for 3,352 healthy control participants in MIGen. RV-EXCALIBER base only accounts for LD between rare variants does not incorporate ‘iCF’ or ‘iCF and gCF’-adjusted expected counts, which are used in RV-EXCALIBER ‘iCF’ and RV-EXCALIBER ‘iCF and gCF’, respectively. A total of 4,815 protein-coding genes with observed counts ≥1 in MIGen and expected counts ≥10 in gnomAD non-Finnish Europeans were used to conduct a rare variant association test. A gene burden test using rare pathogenic alleles was implemented using 3,352 healthy control participants as “cases” in MIGen and gnomAD non-Finnish Europeans as controls. The remaining 2,730 healthy control participants were used as the ranking cohort to generate gCF values. Genomic inflation estimates were evaluated at the median (*λ_med_*). Solid black line indicates an expected uniform distribution of *P*-values under the null and the shaded region represent 95% confidence interval of the expected distribution.

**Supplementary Figure 7:**
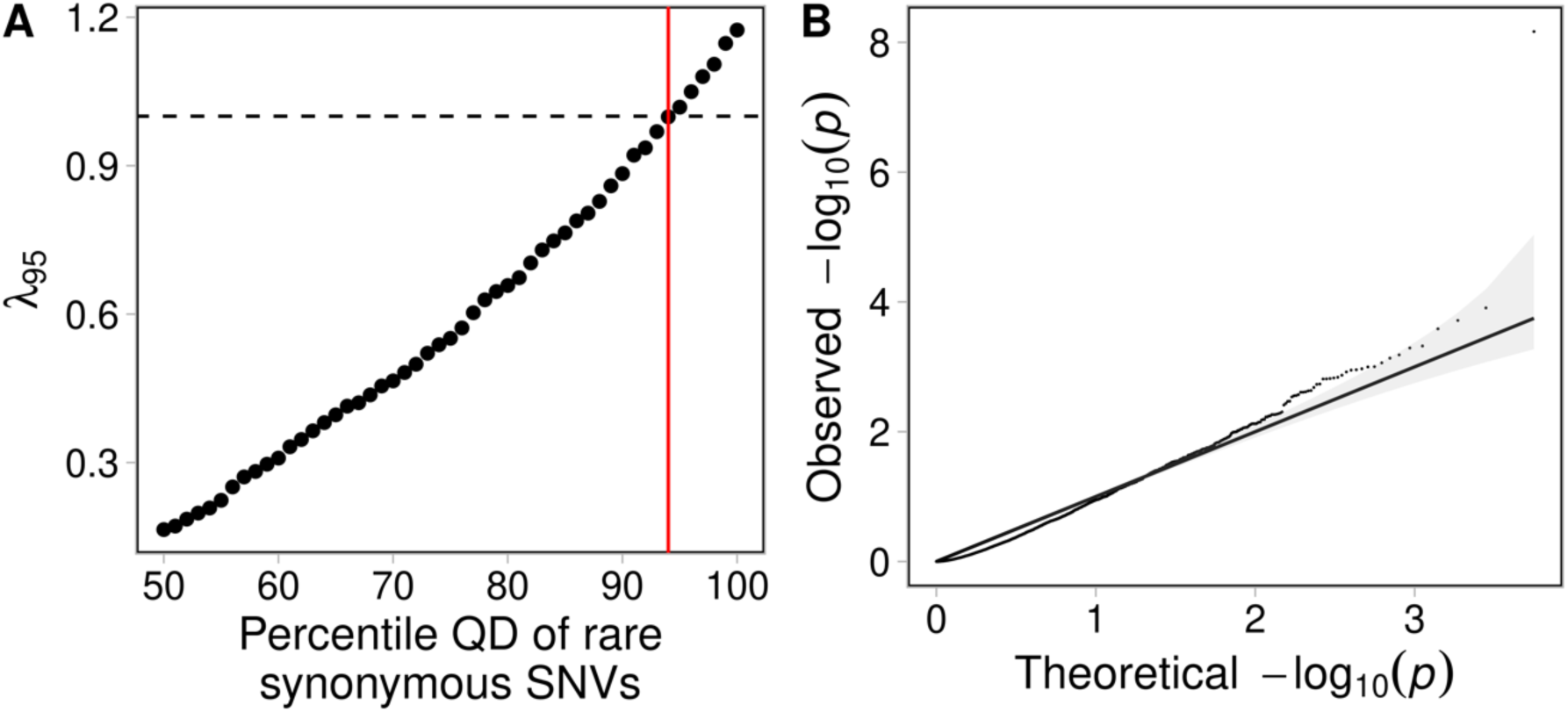
Calibration of gene-based test statistics using quality by depth for rare synonymous SNVs according to the TRAPD algorithm for 3,352 healthy control participants in MIGen. *λ*_95_ estimates were calculated as previously described and were based on the distribution of gene-based test statistics from a gene burden test of rare synonymous SNVs in 3,352 healthy controls from MIGen, which were used as “cases” and non-Finnish Europeans in gnomAD, used as controls. Percentiles were determined based on distribution of quality-by-depth (QD) values for rare synonymous SNVs in MIGen healthy controls **(A)**. These QD scores were compared to the 90th percentile of QD scores for rare synonymous SNVs in gnomAD. Dashed line represents expected value of genomic inflation and the red line indicates the 94^th^ percentile QD from rare synonymous SNVs in MIGen healthy controls, which achieved a *λ*_95_ closest to 1. A quantile-quantile plot demonstrating the distribution of gene-based test statistics based on rare synonymous SNVs corresponding to the 94^th^ percentile of QD scores in MIGen healthy controls and 90^th^ percentile in gnomAD **(B)**. The Solid black line indicates an expected uniform distribution of *P*-values under the null and the shaded region represent 95% confidence interval of the expected distribution.

